# From local seafloor imagery to global patterns in benthic habitat states: contribution of citizen science to habitat classification across latitudes

**DOI:** 10.1101/2024.02.18.580891

**Authors:** Clément Violet, Aurélien Boyé, Stanislas Dubois, Graham J. Edgar, Elizabeth S. Oh, Rick D. Stuart-Smith, Martin P. Marzloff

## Abstract

**Aim:** The aim of this study was to define reef benthic habitat states and explore their spatial and temporal variability at a global scale using an innovative clustering pipeline.

**Location:** The study uses data on the transects surveyed on shallow (< 20m) reef ecosystems across the globe. Time period: Transects sampled between 2008 and 2021. Major taxa studied: Macroalgae, sessile invertebrates, hydrozoans, seagrass, corals.

**Methods:** Percentage cover was estimated for 24 functional groups of sessile biota and substratum from annotated underwater photoquadrats taken along 6,554 transects by scuba divers contributing to the Reef Life Survey dataset. A clustering pipeline combining a non-linear dimension-reduction technique (*UMAP*), with a density-based clustering approach (*HDBSCAN*), was used to identify benthic habitat states. Spatial and temporal variation in habitat distribution was then explored across ecoregions.

**Results:** The *UMAP-HDBSCAN* pipeline identified 17 distinct clusters representing different benthic habitats and gradients of ecological state. Certain habitat states displayed clear biogeographic patterns, predominantly occurring in temperate regions or tropical waters. Notably, some reefs dominated by turf algae were ubiquitous across latitudinal zones. Transition zones between temperate and tropical waters emerged as spatial hotspots of habitat state diversity. Temporal analyses revealed changes in the proportion of certain states over time, notably an increase in turf algae occurrence.

**Main Conclusions:** The *UMAP-HDBSCAN* clustering pipeline effectively characterised fine-scale benthic habitat states at a global scale, confirming known broader biogeographic patterns, including the importance of temperate-tropical transition zones as hotspots of habitat state diversity. This fine-scale, yet broadly-scalable habitat classification could be applied as a standardised template for tracking benthic habitat change across space and time at a global scale. The *UMAP-HDBSCAN* pipeline has proven to be a powerful and versatile approach for analysing complex biological datasets and can be applied in various ecological domains.

## Introduction

Benthic habitats host diverse species and communities (Sunday et al. 2017) and contribute to marine coastal ecosystems functioning and the services they provide to people (Barbier et al. 2011). These services include shoreline protection (Barbier 2017), carbon sequestration (Fourqurean et al. 2012), commercial fisheries (Barbier 2017). As modifiers to abiotic substrates, foundation species, such as kelp, seagrass, and corals engineer biogenic habitats that contribute to specific functions of coastal ecosystems (Elith and Leathwick 2009). For instance, the tridimensional structure of coral reefs can shelter fish assemblages from predators (Hixon and Beets 1993); seaweed or mussel beds can buffer environmental conditions (Jurgens, Ashlock, and Gaylord 2022; Whitaker et al. 2023); and kelp forests are both habitat and food sources for various fish and invertebrate species (Graham J. Edgar, N. S. Barrett, and Morton 2004). Thus, changes in coastal benthic habitats have direct cascading consequences on marine ecosystem structure, functioning and services.

Also hotspots of human activity, coastal ecosystems can be adversely affected by multiple anthro-pogenic stressors (Bowler et al. 2020; Halpern et al. 2019), including global climate change (Burrows et al. 2014; Bowler et al. 2020). The impact of these multiple stressors on benthic communities and ecosystems are frequently mediated by the response of biogenic habitats like kelp, seagrass, or coral (Harley et al. 2006; J. C. Rocha, Peterson, and Biggs 2015). For example, in the vicinity of urban areas, eutrophication can induce the replacement of kelp forests by turf algae (Filbee-Dexter and Wernberg 2018; Pessarrodona et al. 2021), marine heatwaves can lead to coral bleaching, and their intensification in magnitude and frequency can induce long-term decline in tropical coral reefs (Bellwood et al. 2004), and overfishing of herbivorous fish in these reefs can lead to the overgrowth of macroalgae on top of coral reefs (Hughes et al. 2007). Hence, habitat changes are amongst the greatest symptoms of anthropogenic impacts on shallow marine systems, with large consequences for marine biodiversity (J. Rocha et al. 2015). As anthropogenic stressors on marine ecosystems increase and diversify (Halpern et al. 2019), habitat changes will likely become more frequent in the world’s seas (Conversi et al. 2015). Detecting and anticipating future habitat changes in benthic ecosystems requires a thorough understanding of the current state and distribution of benthic habitats and characterising of underlying drivers across multiple scales. Currently, detailed knowledge of habitat distribution is mostly local, i.e. at scales ranging from study sites (10 m - 100 m) or bay (100 m - 10 km) up to regions (10 km - 100 km) (e.g. Robert et al. (2015); Wicaksono, Aryaguna, and Lazuardi (2019)). At larger scales, habitat distribution maps are largely based on physical, geomorphological and biogeochemical ocean properties (e.g. Brown et al. (2011); Lecours et al. (2015); Sonnewald et al. (2020)). At such scales, habitat maps either disregard biogenic habitats or focus on a small number of specific habitat-formers (Assis et al. 2020; McKenzie et al. 2020) and rarely provide information on community composition. Large-scale seafloor habitat maps of either abiotic, or biogenic features also tend to integrate data over large timescales (e.g. decades). Knowledge of benthic habitat changes thus remains highly regional (e.g. Cattano et al. (2020)). In that context several global studies have collated heterogeneous regional monitoring data to document changes in emblematic habitat-formers, such as seagrass spp. (Waycott et al. 2009; Dunic et al. 2021), kelp beds (Krumhansl et al. 2016; Filbee-Dexter and Wernberg 2018) or coral reefs (Eddy et al. 2021). Yet, these independent studies on specific habitat-formers are not sufficient to gain a comprehensive understanding of how the seafloor habitat mosaic has changed through time at a global scale in the face of anthropogenic pressures. Our understanding of current changes in seafloor habitat mosaic is impeded by the lack of large-scale, standardised, data-driven definition and maps of benthic habitat and their potential states.

Identifying changes in benthic habitat states through space and time requires a standardised workflow from data collection through to systematic statistical discrimination between habitat states. In this study, we aim to develop a data-driven pipeline that distinguishes different iconic benthic habitats observed spatially, and apply this to characterise stepwise changes in habitat ecological states through time. Because scientific monitoring programmes are often expensive (Graham J. Edgar, Bates, et al. 2016) and restricted in their spatial and/or temporal coverage (Rhodes et al. 2015), participatory science programmes have emerged as valuable means to increase monitoring programme coverage and resolution. In this study, we leverage the benefits of a citizen science program to characterise benthic habitat states at the global scale and overcome the limitations of traditional scientific programs. The *Reef Life Survey* (*RLS*) relies on standardised diver-based 50-metre-long transects to estimate fish and invertebrate species abundance as well as image-based percentage cover of coastal benthic habitats (Graham J Edgar and Rick D Stuart-Smith 2014). Estimates of habitat percentage cover have already proven useful for defining habitat states at a regional scale through the use of unsupervised machine learning techniques (Cresswell et al. 2017; Pelletier et al. 2020). However, the methods proposed in these previous studies come with a number of limitations when upscaling these approaches at a global scale. In particular, the occurrence and abundance of habitat-forming species are expected to show non-linear responses to environmental changes (Oksanen and Minchin 2002), especially across large environmental gradients. Still, the clustering algorithm used for Cresswell et al. (2017) and Pelletier et al. (2020) are not adapted to take into account the non-linear nature of the dominance patterns between different habitat-forming species.

Hence, we applied a new workflow, combining two algorithms to overcome these challenges: (1) *Uniform Manifold Approximation and Projection* (*UMAP*) a novel dimension reduction technique preserving complex nonlinear structures and patterns (McInnes, Healy, and Melville 2020), (2) and the *Hierarchical Density-Based Spatial Clustering of Applications with Noise algorithm* (*HDBSCAN*) that can identify clusters of varying shapes and sizes while filtering out outlier noise (Campello, Moulavi, and Sander 2013; McInnes, Healy, and Astels 2017). While previous ecological studies have successfully applied both UMAP for dimension reduction (Milošević et al. 2022) and *HDBSCAN* for food web classification (Ohlsson and Eklöf 2020), our study represents a novel application to coastal marine habitats. We interpret classification results by combining the latest *SHapley Additive exPlanations* (*SHAP*) (Lundberg and Lee 2017) framework with visual inspections of photoquadrats associated with the most representative transects of the different clusters.

Therefore, the aim of this study is to characterise coastal benthic habitat states using a *UMAP-HDBSCAN* pipeline on the RLS habitat dataset. Using this pipeline, we identify and classify benthic habitat states at a global scale and characterise their spatial and temporal variability across biogeo-graphical gradients as well as within bioregions.

## Materials & Methods

We used a *UMAP-HDBSCAN* pipeline to cluster the global *RLS* benthic habitat dataset. In the following sections, we sequentially describe : (1) the data used in this study, (2) the clustering pipeline, (3) the interpretation of the identified clusters.

### Data

#### *Reef Life Survey* photoquadrat dataset

The *RLS* (http://www.reeflifesurvey.com/) is a hybrid citizen science/professional researcher program monitoring reef communities around the world using scuba-diving visual census. Details about the survey methods, including protocols, diver training, data quality assurance and data management, are covered by Graham J Edgar and Rick D Stuart-Smith (2014). Here, we used estimates of relative cover of benthic habitats derived from in situ digital photoquadrats: along standardised 50 m transect, 20 photoquadrats, which each approximately covers 0.3 m × 0.3 m, are collected every 2.5 m (Graham J. Edgar, Cooper, et al. 2020). Images are then annotated using point counts on the Squidle+ (https://squidle.org/) platform to estimate the percentage covers of about 50 substratum types and functional groups, based on the *CATAMI* benthic imagery classification scheme (Althaus et al. (2015); for further details, see Graham J. Edgar, Cooper, et al. (2020)). Based on *RLS* specialists’ expertise, these 50 original benthic habitat categories (see Appendix A, Table 1 in Supporting Informations) were grouped into 24 broader categories (Table 1) that more consistently capture the range of dominant coastal substratum available along *RLS* transects at the global scale.

We extracted the *RLS* photoquadrat dataset on 24 January 2023. From the original 8,154 transects, we removed partially scored transects. For transects annotated multiple times on *Squidle+* across various research projects, mean percentage cover estimates were considered. After fully curating the dataset, the photoquadrat dataset consisted of 6,554 transects across 2,249 sites over the world. All subsequent analyses were performed at the transect level to consider local-scale variation in the state of benthic habitats.

**Table 1.**
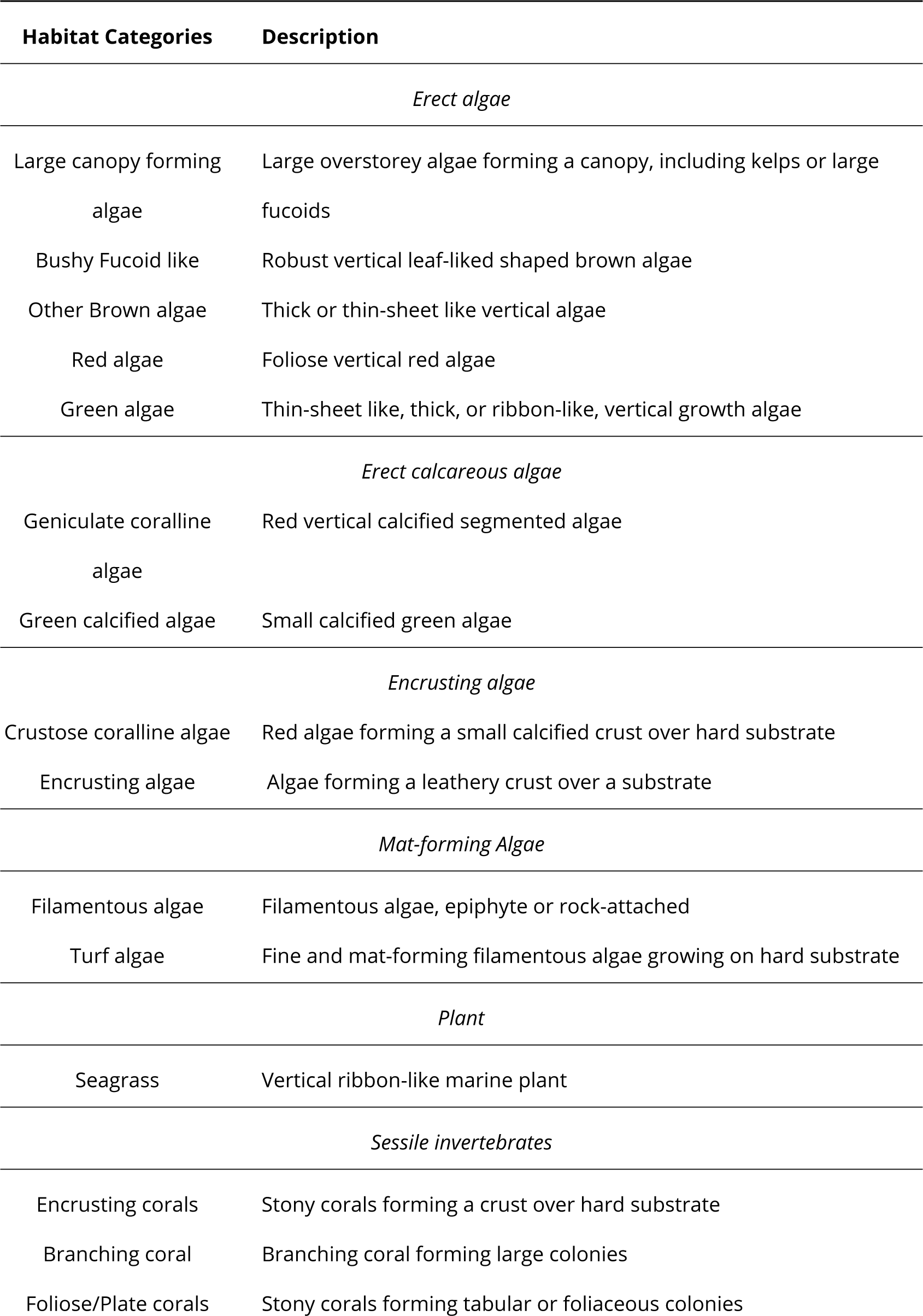
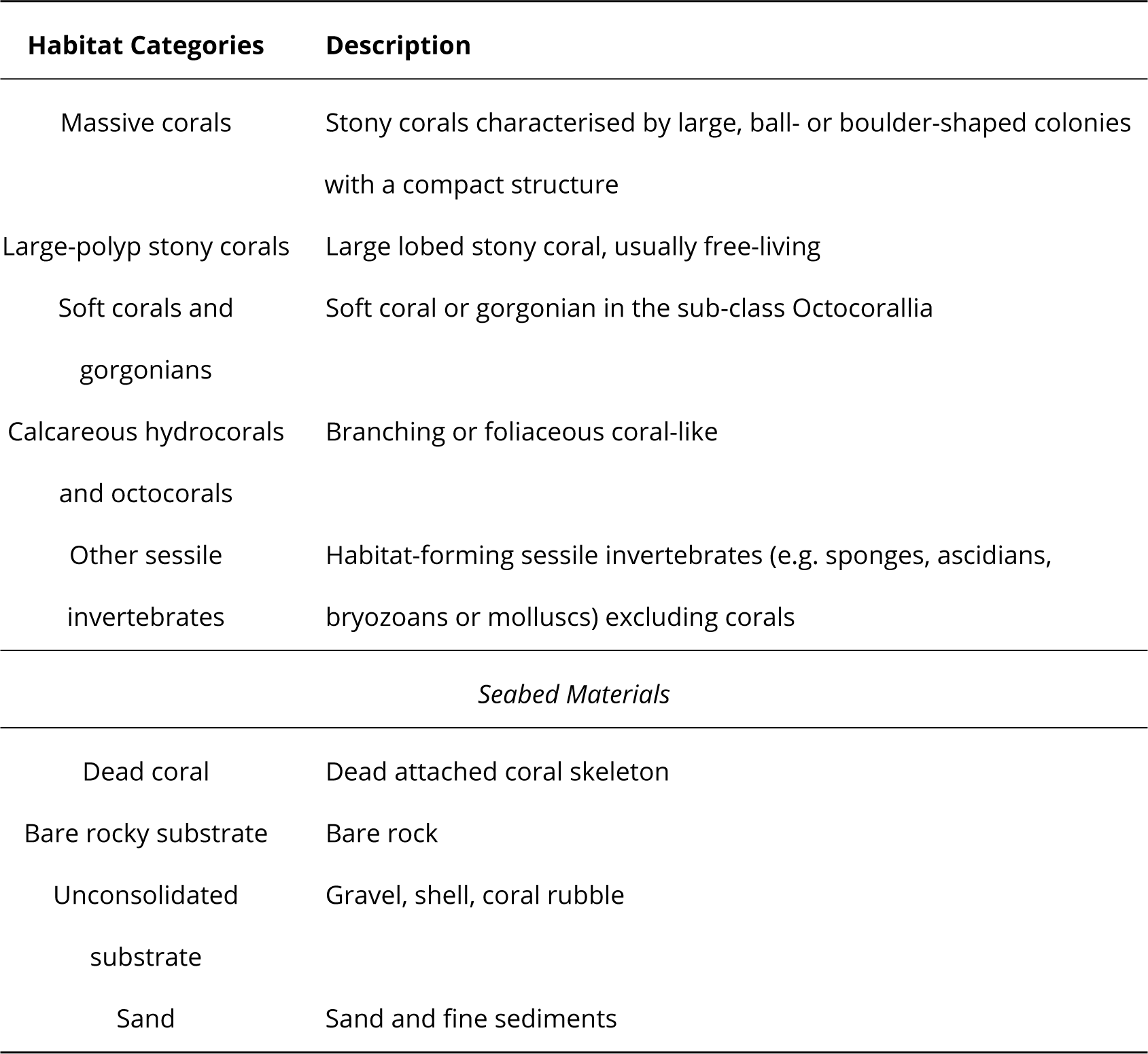
Description of the 24 categories used in this study to overall capture the diversity of habitat types sampled by *Reef Life Surveys* worldwide. The 50 original *RLS* categories were lumped into these 24 categories that represent ecologically-consistent groups associated with different levels of structural complexity.

### Clustering pipeline

To account for non-linear, high-dimensional and complex nature of the ecological data, we combined a graph theoretical dimension reduction technique and a density-based classification technique, which has successfully identified ecoprovinces using biogeochemical ocean data at a global scale (Sonnewald et al. 2020). Among the set of methods available, we have chosen the *UMAP* algorithm (McInnes, Healy, and Melville 2020) and the *HDBSCAN* algorithm (Campello, Moulavi, and Sander 2013; McInnes, Healy, and Astels 2017) for dimension reduction and clustering, respectively.

### Dimension reduction - *UMAP*

The *UMAP* algorithm is a non-linear reduction technique (McInnes, Healy, and Melville 2020). Unlike more traditional methods applied in ecology such as *Principal Component Analysis* (*PCA*), *UMAP* preserves both the local structure (preserving the distance between neighbouring points) and the global structure (preserving the distances between the most different points) of the raw dataset (McInnes, Healy, and Melville 2020). These two key properties have been proven useful for reducing the dimension of complex genomic (Dorrity et al. 2020), or ecological (Milošević et al. 2022) data prior to clustering. *UMAP* reduces the dimensionality of a dataset by first creating a high-dimensional graph that connects each data point to its k-nearest neighbours. Then, *UMAP* produces a low-dimensional representation of this high-dimensional graph that reflects the original dataset (McInnes, Healy, and Melville 2020). *UMAP* requires a distance matrix to construct the initial k-nearest-neighbour graph. Here, we applied the Chord transformation to standardise percentage cover data as relative cover per transect before computing euclidean distances between transects (Legendre and Gallagher 2001). In addition to the choice of a suitable distance metric, two *UMAP* hyperparameters can influence dimension reduction. The first one is the number of neighbours (*n_neighbors*) to consider when creating the k-nearest neighbour graph. Low *n_neighbors* values will allow the embedding to preserve more of the local structure of the original distance matrix and larger ones will preserve more of the global structure (McInnes, Healy, and Melville 2020). The second parameter is *min_dist*, which controls the packing density at which UMAP is allowed to clump similar points in the reduced dimensional space. A high value of *min_dist* will tend to preserve the overall topological structure of the data, while a low value allows UMAP to clump closely similar points on the embedding. The value of n_neighbors has been tuned in this study, while the value of *min_dist* has been set to 0.0, since this value allows densification of the low-dimensional representation of the dataset, which is important before using a density-based classification algorithm (Vermeulen et al. 2021).

### Clustering - *HDBSCAN*

After embedding our data into a two-dimensional space, we clustered the generated projections of the data with the unsupervised hierarchical density-based clustering *HDBSCAN* algorithm that can provide both hard (i.e. samples are exclusively assigned to a single cluster) and soft (i.e. samples are assigned probabilities of belonging to the different clusters) clustering solutions. In addition to identifying clusters of various shapes and density from a dendrogram, this algorithm comes with several advantages in ecology both in terms of classification and interpretation: it can exclude noisy observations, which do not get assigned to any clusters, and can also highlight most representative members of each cluster (Campello, Moulavi, and Sander 2013; McInnes, Healy, and Astels 2017). The *HDBSCAN* clustering algorithm involves a few core steps. First, it computes the core distance for the k-nearest neighbours for all points in the dataset. Then, it computes the extended minimum spanning tree from a weighted graph, where the edges are weighted by the distance between two points while taking into account the density of points around them. Then, *HDBSCAN* builds a hierarchy from the extended minimum spanning tree by cutting it at different levels of density. If the cut results in the creation of clusters smaller than the minimal number of observations set by the user *min_cluster_size*, all points members of these clusters are declared as noise by the algorithm. The algorithm stops when it declares all points as noise and returns to the user a tree-like structure where each node corresponds to a cluster varying in shape and density (Campello, Moulavi, and Sander 2013; McInnes, Healy, and Astels 2017). In this study, we tuned only one parameter for *HDBSCAN*: the *minimum_cluster_size*, controlling for the minimal number of observations required to form a cluster and used the default parameters otherwise.

### Evaluation of the clustering output

For this pipeline, we search the best combination of hyperparameters for both *UMAP* (*n_neighbors*) and *HDBSCAN* (*minimum_cluster_size*) using a complete grid search. We exhaustively explored results sensitivity to the two hyperparameters from 10 to 500 resulting in 241,081 models evaluated. The best combination was found by optimising both the quality of the embedding and the clustering, using two criteria. The *UMAP* embedding was evaluated with the trustworthiness metric (Venna and Kaski 2001), ranging from 0 to 1 (the higher the index the more the local structure of the original data is preserved). The quality of the clustering was evaluated with the *DBCV*, which measures both compactness within and separations between clusters (Moulavi et al. 2014). The *DBCV* index, which ranges between −1 and 1, is appropriate to assess the quality clusters with varying shapes and densities (Moulavi et al. 2014).

A previous fine-scale analysis by Cresswell et al. (2017) on a regional subset of this dataset yielded nine groups of habitats. We expected to find at least that many groups at the global scale and thus restrained our search of the best hyperparameter combinations to the solution yielding at least the same number of cluster than Cresswell et al. (2017). Among these solutions, we select the best combination of hyperparameter (*n_neighbors* = 400; *min_cluster_size* = 74) yielding the best performance in terms of both their trustworthiness and DBCV scores, while having the maximal number of clusters for a finer granularity.

### Interpretation of the clusters

To interpret individual clusters identified with *UMAP-HDBSCAN*, we computed the mean percentage cover of each habitat in each cluster. Then we used the *SHAP* framework to further explore how potential nonlinear interactions between variables may determine clustering outcomes (Lundberg and Lee 2017). Because of the computational cost of applying *SHAP* to our complete pipeline, we used a classification tree (Breiman et al. 1984) to approximate the clustering pipeline (i.e. predict label cluster membership based on the raw percentage cover variables) before applying the *SHAP* framework (Lundberg and Lee 2017). In order to train our classification tree, we used a stratified train-test split to ensure that the relative frequency of each cluster label is preserved in the train and test fold. The training and the test sets contain 80% and 20% of the data, respectively. Then, we used a minimal cost-complexity pruning algorithm to avoid overfitting of our classification tree (Breiman et al. 1984) estimated classification error rates using the F1-score (Van Rijsbergen 1979). The classification error rates were satisfactory, F1-score of 0.99 and 0.94 on the train and test sets respectively. Based on the *SHAP* values that estimate the influence of each variable to cluster definition, we examined potential interactions between the two most characteristic variables for each cluster by performing a piecewise linear interpolation of the *SHAP* values. Finally, we completed interpretation by extracting the photoquadrats for these transects considered by HDBSCAN as the most representative members of their cluster.

### Spatio-temporal distribution of benthic habitat states

We first explored the latitudinal distribution of each cluster. We also summarised their occurrence within each of the Marine Ecoregions of the World (*MEOW*; Spalding et al. (2007)) sampled by the *RLS*. In addition to examining dominant clusters per ecoregion, we also computed the proportion of transects classified as noise, as well as the Gini-Simpson diversity index. We chose this diversity index because it focuses on changes in dominance patterns, more indicative of changes in landscapes and is more robust to low sampling issues than other diversity indices (Lande, DeVries, and Walla 2000).

Finaly, We investigated temporal trends at the site level by comparing the proportion transects classified into the different habitat states at five different sites temperate Australian sites already monitored by Rick D. Stuart-Smith et al. (2022). We were interested to determine the extent to which the proportion of transects classified according to the different habitat states at a site could be an indicator of the overall ecological status of it.

## Results

Based on extensive exploring of hyperparameter space, both trustworthiness score of 0.98 ± 0.002 (mean ± sd) and *DBCV* score of 0.46 ± 0.08 (mean ± sd) for solution containing at least 9 group suggest that these solutions yield reliable clustering of the *RLS* photoquadrat dataset (Annexe B Fig. S1 in Supporting Informations). Across these best solutions, optimal number of clusters varied between 9 and 184 (22.81 ± 18.37; mean ± sd) while mean number of points classified as noise was 2, 207.33 ± 364.95 (mean ± sd). Hereafter, out of theses, we chose to focus on the single solution yielding the highest resolution (i.e. the greatest number of clusters; see Fig. 1 for a description of the habitat states uncovered), and the smaller number of transects classified as noise (1,464 transects) possible. This solution has a trustworthiness score of 0.98 for *UMAP* and a *DBCV* score of 0.60 for *HDBSCAN*. The number of clusters identified by this set of hyperparameters is 17 (Fig. 1).

**Figure 1.**
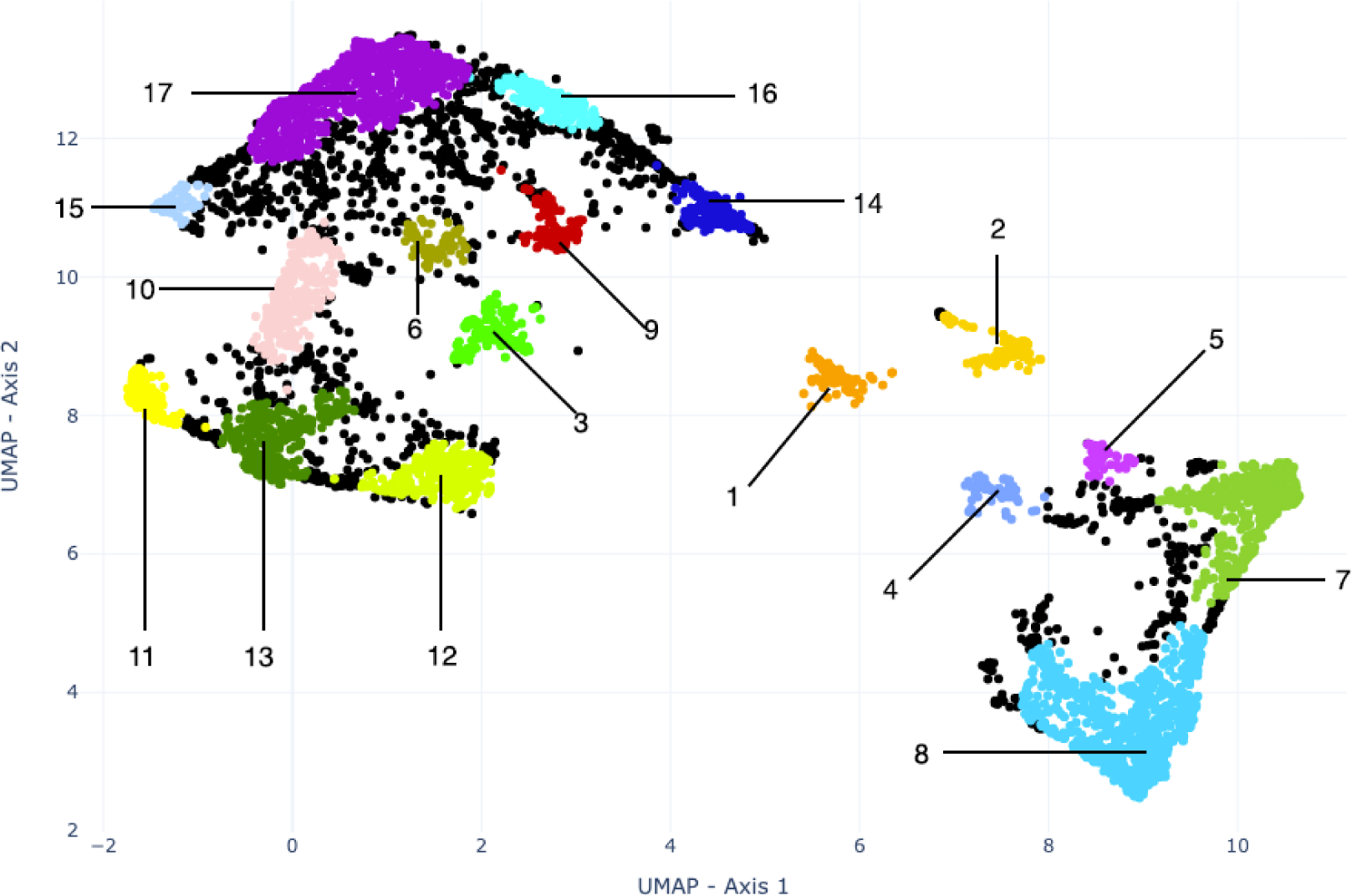
Two-dimensional *UMAP* embedding of the benthic cover data of the 6,554 *RLS* transects. Each point corresponds to an *RLS* transect, coloured according to membership for the selected *UMAP-HDBSCAN* pipeline. Black dots represent points classified as noise (n=1464). The 17 clusters can be interpreted as follows (see Fig. 2 and S20-36): 1. *Foliose brown algae* (n=148) 2. *Filamentous algae* (n=208) 3. *Other Sessile invertebrates* (n=185) 4. *Foliose red algae* (n=123) 5. *Seagrass* (n=83) 6. *Soft coral and gorgonians* (n=98) 7. *Bushy fucoids* (n=577) 8. *Large Canopy forming algae* (n=894) 9. *Unconsolidated substrate* (n=151) 10. *Crustose coralline and turf algae* (n=286) 11. *Green calcified algae* (n=166) 12. *Bare substrates* (n=329) 13. Crustose coralline algae (n=409) 14. *Sand* (n=220) 15. *Branching coral* (n=110) 16. *Turf and sand* (n=207) 17. *Turf algae* (n=897)

The 17 clusters identified can be summarised hereafter according to four broad groups (Fig. 2; Fig. 3, see Fig S2-S19 for their distribution on the globe and S20-36 for their interpretation with *SHAP* framework in Supporting Informations): (1) temperate habitats, (2) subtropical and tropical habitats, (3) broadly-distributed habitats and (4) opportunistic habitats (i.e. habitats with documented ecologi-cal dysfunctions - and therefore often habitats under strong anthropogenic influence, characterised by the presence of filamentous algal species or turf).

**Figure 2.**
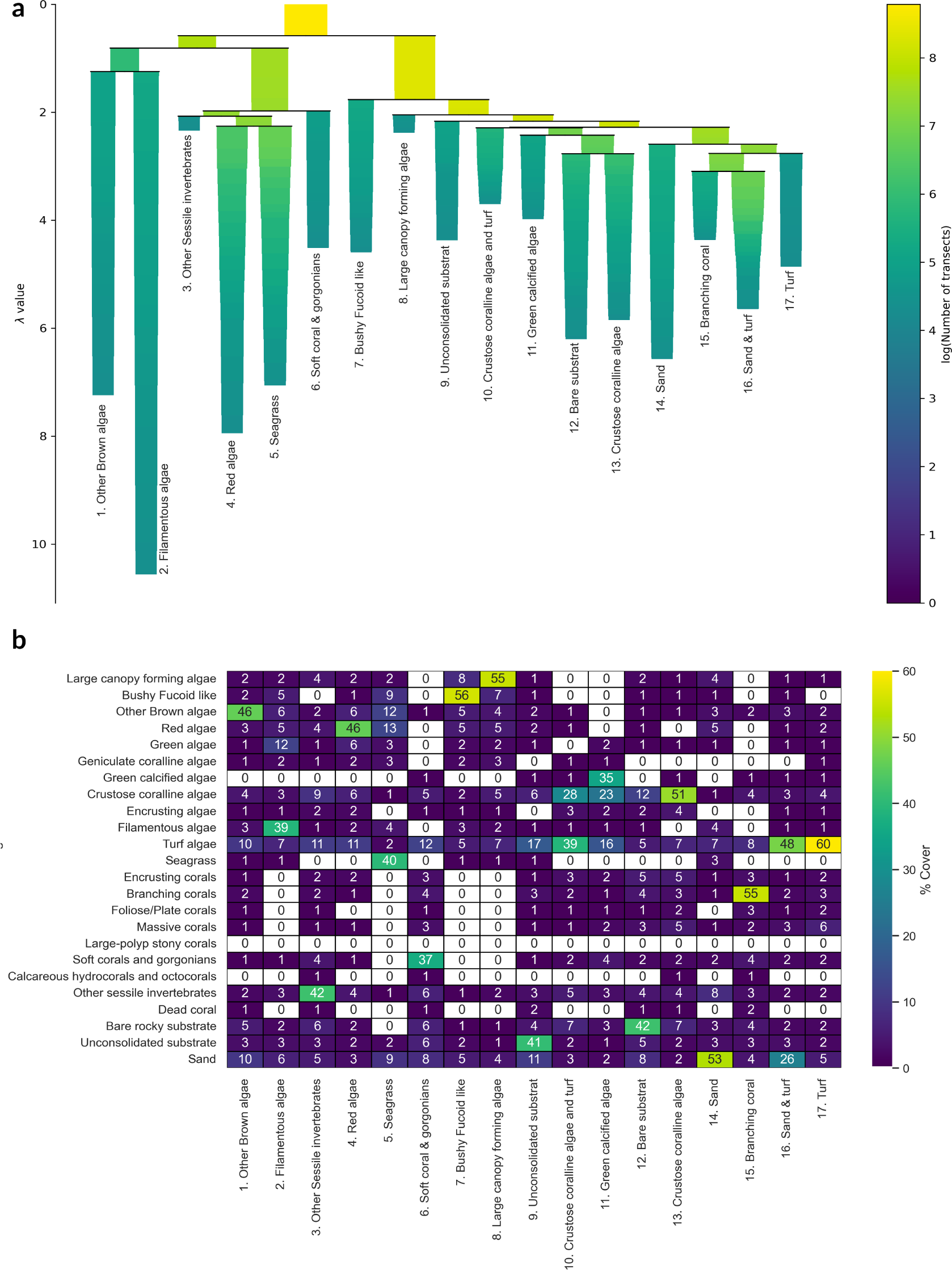
a. *HDBSCAN* condensed clustering tree of the *UMAP* 2D embedding b. Heatmap of the mean substrate coverage (rounded to the nearest integer) for each cluster identified by the *UMAP-HDBSCAN* pipeline.

**Figure 3.**
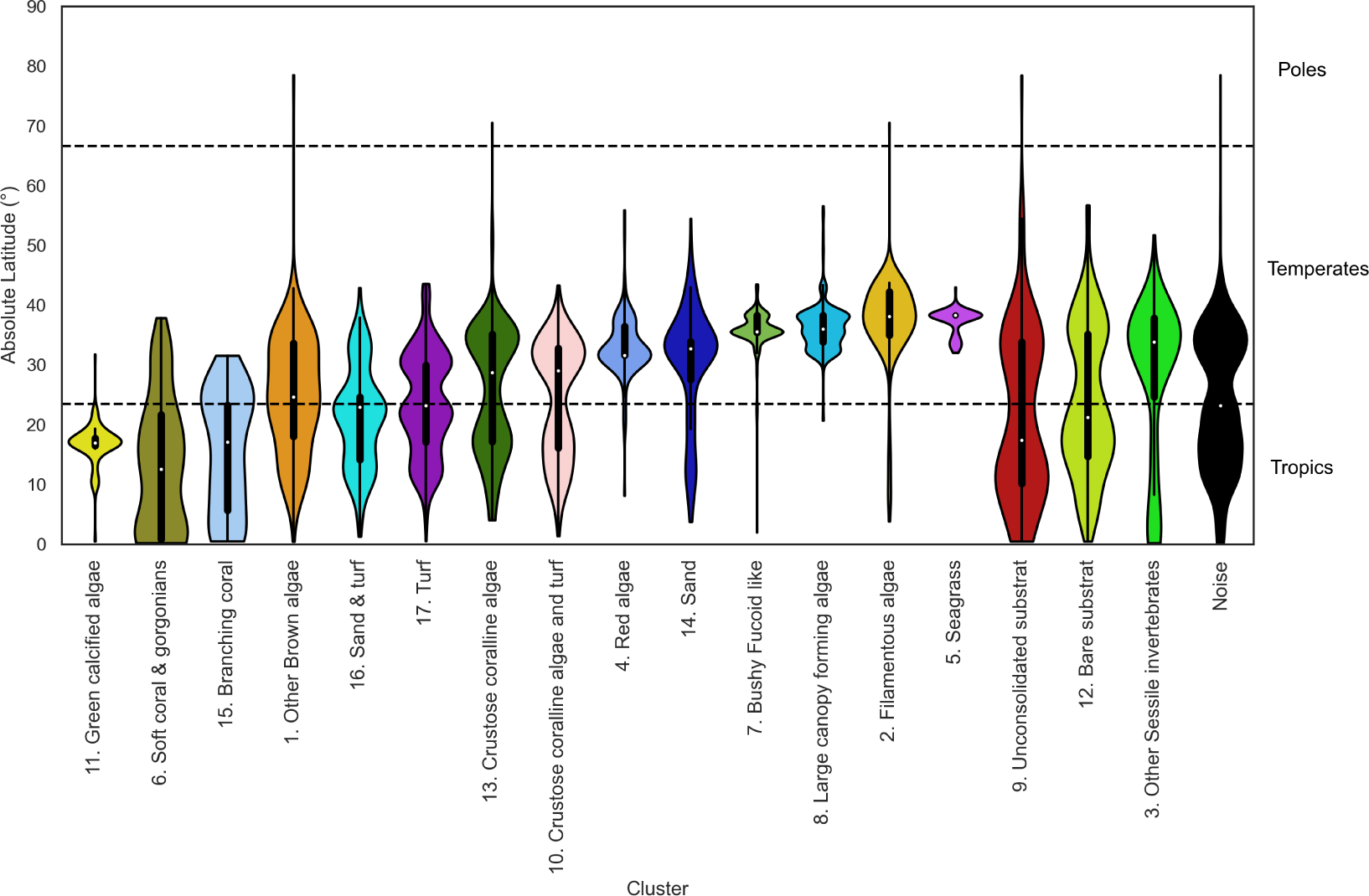
Violin plot of the absolute latitudinal distribution of the different hard cluster solutions.

Transects within temperate regions can be classified according to five major clusters associated with contrasted dominance of sessile invertebrates, foliose red algae, seagrass, bushy foliose algae and canopy-forming algae, as follows: cluster 3 is dominated by at least 30% and on average 42% of sessile invertebrates. Cluster 4 is dominated by at least 40% coverage of foliose red algae. Cluster 5 is dominated by at least 30% and on average 40% seagrass. Cluster 7 is dominated by at least 20% coverage and an average of 56% fucoid bushy algae and an absence of canopy forming algae. Cluster 8 is characterised by a cover of at least 20% and an average of 55% of canopy forming algae with an absence of fucoid bushy algae.

Three clusters correspond to tropical and sub-tropical habitat types. Cluster 6 which is charac-terised by at least 30% and on average 37% of soft corals and gorgonians. Cluster 11 is composed of 20% coverage and an average of 35% green calcified algae. Finally cluster 15 is composed of at least 35% and on average 55% branching coral. Interestingly, this is the only group of corals identified in the dataset given the four categories of colony-forming corals.

Five clusters correspond to broadly-distributed habitats that can occur across both temperate and tropical latitudes. Cluster 1 is dominated by at least 30% and on average 46% brown foliose algae. Cluster 9 is dominated by the presence of at least 30% and on average 41% unconsolidated substrate. Cluster 12 has at least 30% and on average 42% bare substrate. Cluster 13 is characterised by 40% and on average 51% of crustose coralline algae with an absence of turf algae. Cluster 14 has at least 30% and an average of 53% sand without turf algae.

Finally, four clusters correspond to opportunistic habitats. Cluster 2 is in that respect dominated by at least 30% coverage and an average of 39% filamentous algae. Clusters 10, 11 and 17 are all dominated by turf algae. Cluster 10 is composed of at least 30% and on average 39% of turf algae and at least 20% and on average 28% of crustose coralline algae. Cluster 16 is characterised by the presence of at least 30% and on average 48% turf algae and a minimum coverage of 20% and on average 26% sand. Cluster 17 is composed of at least 40% and on average 60% turf algae with an absence of crustose coralline.

The clusters identified by the *UMAP-HDBSCAN* pipeline show a marked latitudinal gradient (Fig. 3). *Red algae*, *filamentous algae*, *fucoids*, *large canopy-forming algae* and *seagrass* are essentially distributed overall in the temperate zones across latitudes higher than 25° (Fig. 3). Conversely, four habitat states, namely *soft corals and gorgonians*, *green calcified algae*, *sand and turf* and *branching coral* essentially occur in tropical latitudes (lower than 25°) (Fig. 3). However, some groups are relatively ubiquitous across all surveyed latitudes such as those associated with transects classified as *bare substrate* and *unconsolidated substrate*, *brown algae*, *crustose coralline algae* with and without *turf algae* and *turf algae* (Fig. 3). It should also be noted that the transects considered to be noisy are also evenly distributed across all latitudes (Fig. 3).

The spatial distribution of transects sampled by RLS volunteers is particularly concentrated in Australia (Fig. 4 a). However, other areas such as the Caribbean, the Azores, and French Polynesia have also been extensively surveyed with more than 50 transects (Fig. 4 a). Globally, three habitat types dominate in terms of occurrences across all surveyed ecoregions, namely *bare substrate* (n = 20), *turf algae* (n = 17), and *large canopy-forming algae* (n = 11). These three habitat types dominate in 37% of the ecoregions sampled by the *RLS* (Fig. 4 b). Two habitat types identified by the *UMAP-HDBSCAN* pipeline, *seagrass* and *red algae*, are not dominant in any of the world’s ecoregions. The patterns of dominance of the different clusters also vary along the latitudinal gradient (Fig. 4 b), in line with the latitudinal distribution of each cluster (Fig. 3). These latitudinal variations of dominance are visible both at a global scale, but also along certain regions. For instance, a decrease in prevalence of sites in the *large canopy-forming algae* cluster accompanies an increase in sites in the *turf algae* cluster along the coastline from southern to northern Australia (Fig. 4 b).

**Figure 4.**
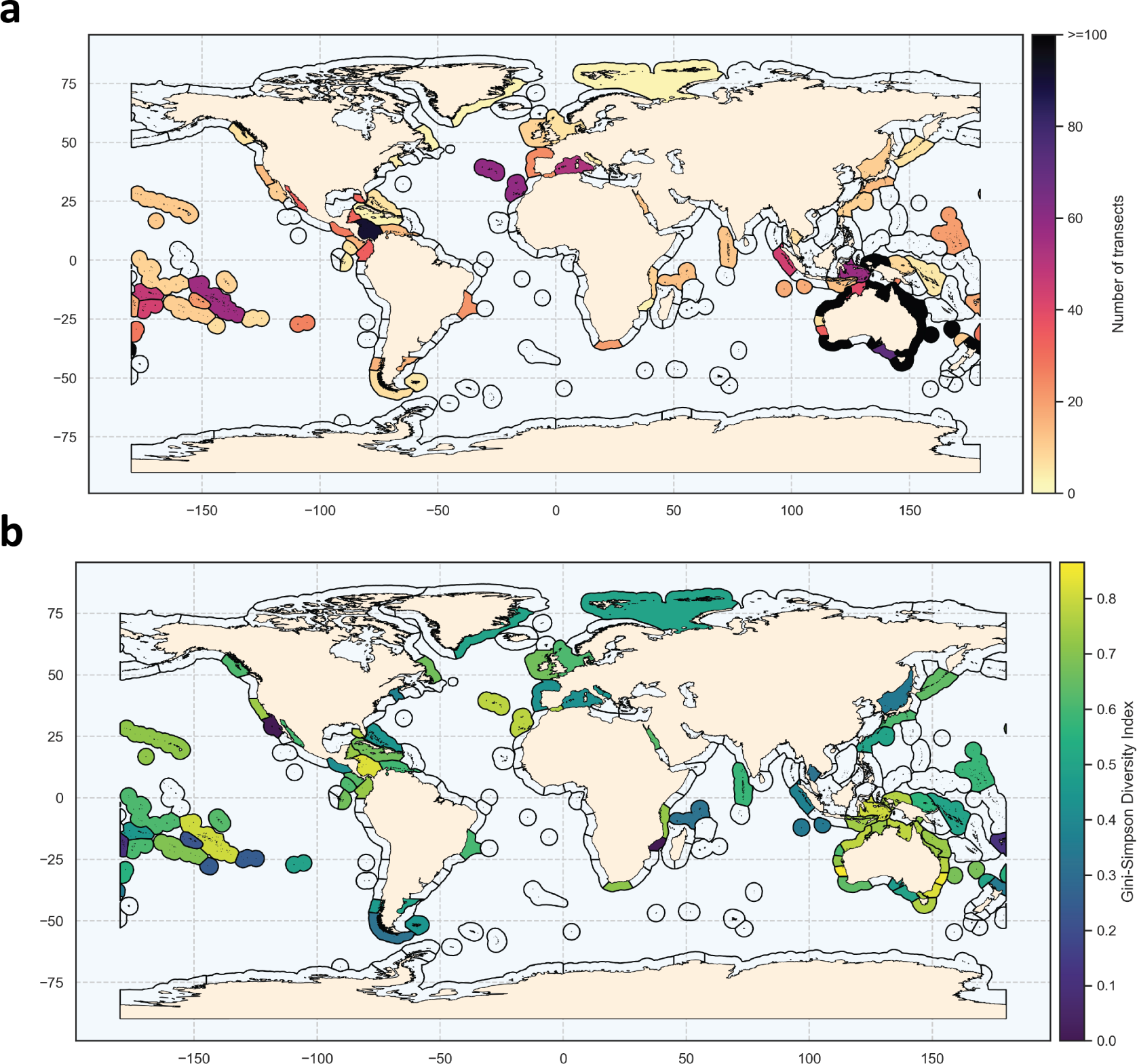
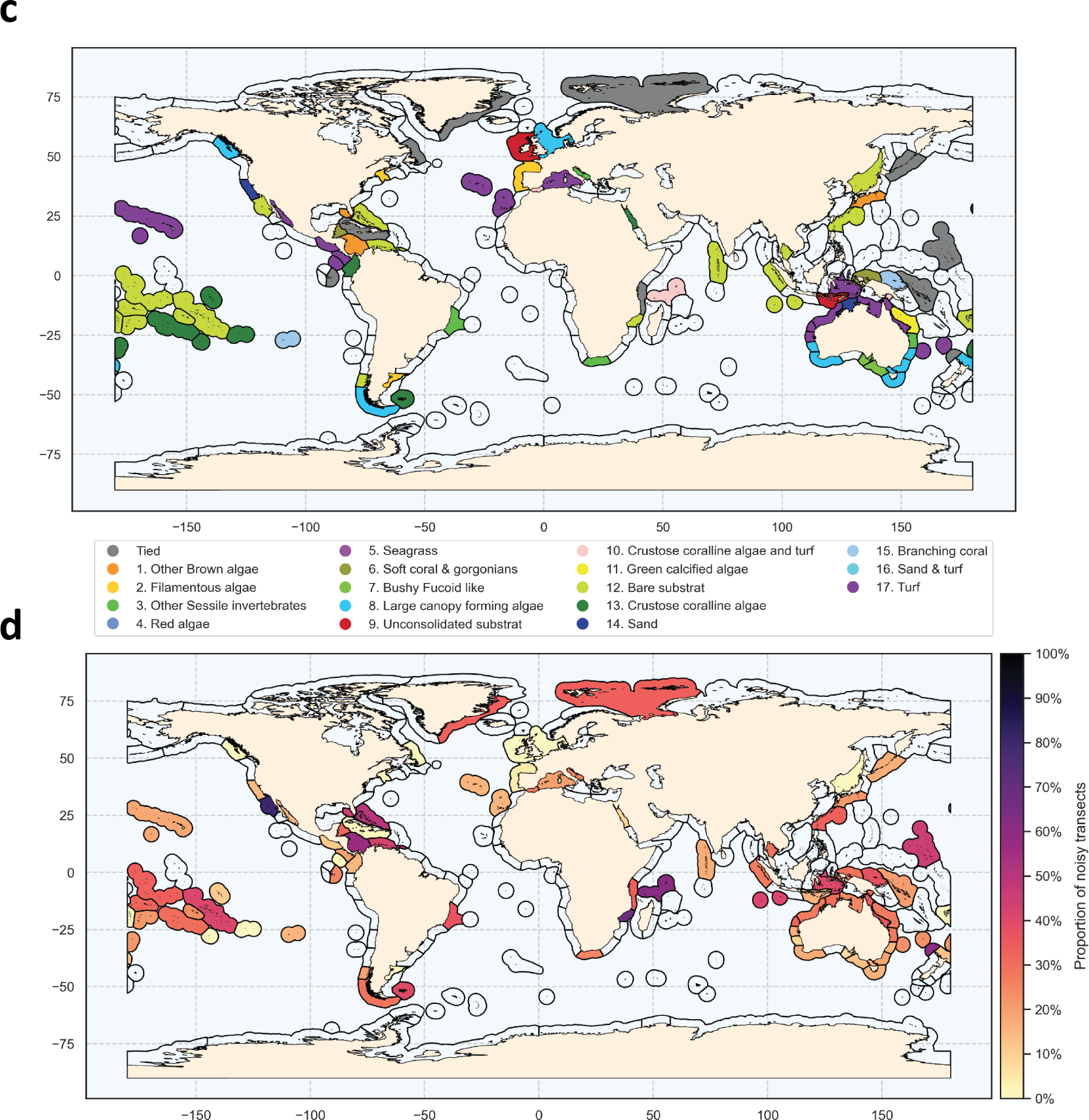
a. Spatial distribution of reef surveys from the Reef Life Survey database used for analyses. b. Map of dominant clusters in each MEOW ecoregion. Dominant clusters were determined as the greatest count of transect labels in each ecoregion. c. Spatial distribution of the proportion of transects classified as noise in each ecoregion. d. Gini-Simpson diversity index calculated by the occurrence of clusters in each ecoregion of the world.

The proportion of noisy transects is highly heterogeneous across the globe (Fig. 4 c). Noisy transects represent 23% of all transects analysed, but are present in some areas more than in others. For example, in the Southern California Bight (western USA), Bight of Sofala/Swamp Coast (Eastern Africa), the Seychelles, and in Three Kings-North Cape (northern New Zealand), at least 60% of transects are classified as noisy (Fig. 4 c). While these four ecoregions share in common a low number of transects sampled (Fig. 4 a), no significant correlation was found between the proportion of transects classified as noisy and the number of transects done in each ecoregion (*τ_Kendall_* = 0.05, p = 0.54; Fig. S37 Supporting Information). Moreover, 12 ecoregions sampled out of the 83 by the RLS had no transects that were classified as noisy (Fig. 4 c).

Areas with the highest diversity of habitat types, based on both the number of clusters occurring and on their relative proportions in the ecoregions, are concentrated in eastern and western Australia, as well as in the Caribbean and the Tuamotus (Fig. 4 d). Areas with the lowest Gini-Simpson values are the Southern California Bight (western USA) and Bight of Sofala/Swamp (eastern Africa) Coast with a Gini index of 0 (Fig. 4 d). It should be noted, nevertheless, that there is a weak correlation between the Gini-Simpson index and the number of transects carried out in the ecoregion (*τ_Kendall_* = 0.29, p < 0.001; Fig. S38 in Supporting Information).

At *Reef Life Survey* site level, temporal changes in the occurrence of the different clusters can provide useful indicators of ecological changes (Fig. 5). For instance, at a given site, changes in yearly proportions of transects classified as *large canopy forming algae* tend to match with annual mean percentage cover of *large canopy forming* algae estimated across transects (Fig. 5 and Fig. S39). Moreover, at certain sites where the cover of *large canopy forming algae* decreased, changes in the dominance patterns of habitat states offer further insights about ongoing ecological changes. At Beware Reef (Fig. 5 b), *large canopy forming algae* disappeared in favour of *other sessile invertebrates* whereas at Port Phillips Heads (Fig. 5 d), decrease in the proportion of transects classified as canopy forming algae between 2013 and 2017 was counterbalanced by an increase in the proportion of transects classified as *bushy fucoids* or *filamentous algae*. This site also experienced a moderate inter-annual variability in the proportion of transects classified as *noise*. A long-term decrease in the proportion of transects classified as *noise* was however observed at other sites (e.g. Batemans or Beware Reef; Fig. 5 a and b), where a turnover through time in the dominating habitat state occurred.

**Figure 5.**
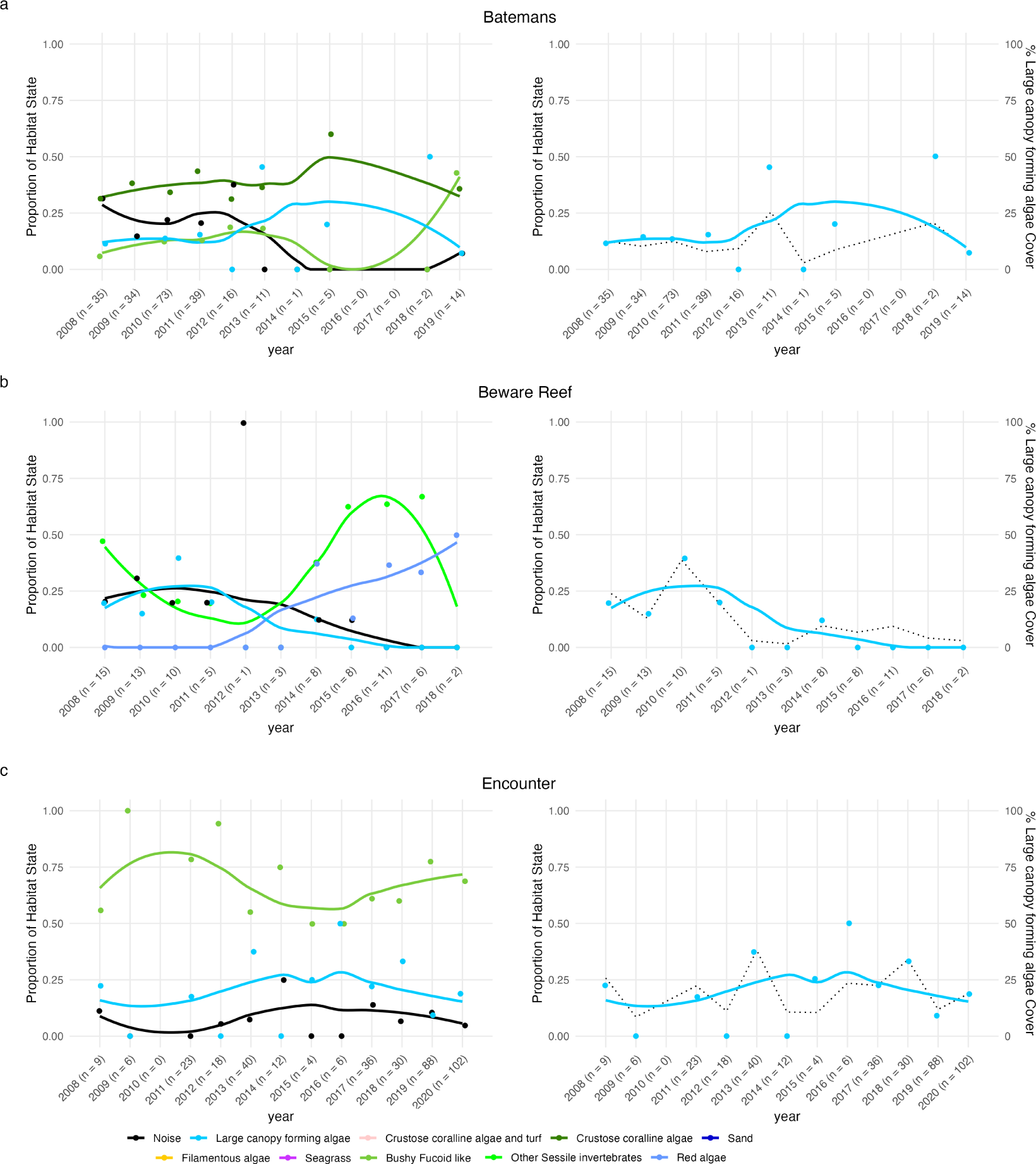
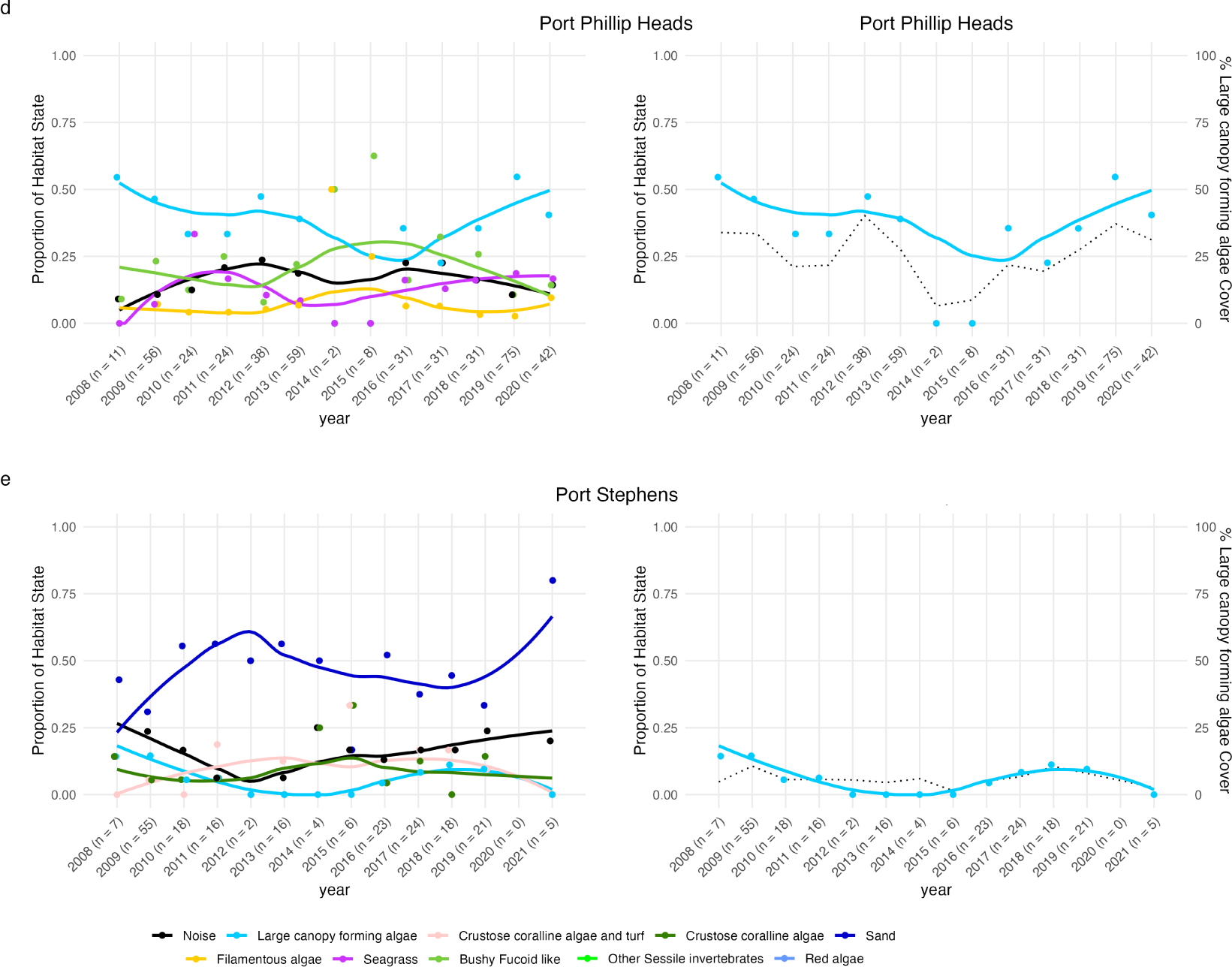
Temporal evolution of the proportion of transects classified into the different habitat states at different sites: a. Batemans, b. Beware Reef, c. Encounter, d. Port Phillips Heads, e. Port Stephens. Habitat states are colour-coded as follows: light blue for large canopy-forming algae; pink for crustose coralline and turf; deep green for crustose coralline algae; deep blue for sand; yellow for filamentous algae; light green for bushy fucoid; flashy green for other sessile invertebrates; light blue for red algae; purple for turf. Noise appears in black). The dots represent the proportion of transects in each category in each year and the trend lines are LOESS regression models weighted by the number of transects per year. For visual clarity, the left-hand column only shows the subset of groups, which proportion most varied over time (see Fig. S40 for the full version). The right-hand column focuses solely on the yearly proportion of transects classified as Canopy forming algae (blue line) in comparison with the annual mean percentage cover of Canopy forming algae estimated across all transects (black dotted line).

## Discussion

The *UMAP-HDBSCAN* clustering pipeline identified 17 distinct clusters within all the *RLS* transects performed globally across a range of coastal temperate and tropical regions. Within these groups, we found different biogenic habitats whose distribution patterns match with current biogeographic knowledge of benthic ecosystems: for example, *bushy fucoid algae*, and *large canopy-forming algae* predominantly occur in temperate waters (Assis et al. 2020; Jayathilake and Costello 2020), while soft corals and gorgonians, and branching coral are more frequent in tropical waters (Jones et al. 2019; Wirabuana et al. 2019). Our analysis also highlights habitat types that occur across the globe, including (1) different granulometric facies like *sand*, *unconsolidated substrate*, and *bare substrate*, as well as (2) different habitat types dominated by low-profile algae, such as *crustose coralline algae* or *turf algae*. The latter are known to occur across the globe and can dominate benthic substrates in diverse conditions (Connell, Foster, and Airoldi 2014; Liu et al. 2018).

In addition, this classification also distinguishes between different ecological states of these habitats (hereafter refers to as “habitat state”), including known alternative successional stages, or different degradation states of these habitats (Fig. 6). For example, the clusters *crustose coralline algae*, *crustose coralline algae and turf* and *turf* provide an interesting template to describe the habitat transitions described in Cornwall et al. (2023), which suggests that a shift from *crustose coralline algae* to *turf* domination reduces reef carbonate production. Similarly, the clusters *branching coral*, *turf and sand* and *turf* can be used to describe and quantify in a standardised manner the transitions between corals and turf dominated habitats that are occurring more frequently due to anthropogenic pressures (Jouffray et al. 2015). Fig. 5 b also illustrates the occurrence along the southeastern Australian coastline of alternative ecological states on temperate reefs, where dense macroalgal canopies dominated by *Ecklonia radiata* (here *large canopy-forming algae*), can shift to extensive barrens (here *bare substrate* or *crustose coralline algae*) following destructive grazing by the long-spined sea urchin *Centrostephanus rodgersii* (Ling 2008). Thus, our approach can classify reef cover data collected across the globe with the *RLS* protocol into an ecologically sound template to explore common reef habitat transitions under anthropogenic pressures (Donovan et al. 2018). Some of the habitat states identified here on the global *RLS* dataset match well with the habitat states previously identified by Cresswell et al. (2017), who applied another clustering approach to an Australian subset of the *RLS* dataset. In particular, some of our groups (i.e. *large canopy forming algae*, *turf algae*, *filamentous algae* and *branching coral*), match well with four out of the nine habitat states identified by Cresswell et al. (2017) (i.e. “Canopy algae”, “Turf”, “Epiphytic filamentous algae–caulerpa” and “Coral”; see Table 1 in Cresswell et al. (2017) for a detailed description of their habitat states and Fig. 2 b in this paper for comparison). Furthermore, we identified more finely-resolved habitat states here relative to the classification proposed by Cresswell et al. (2017). The clusters *crustose coralline algae* and *bare substrate* identified here are amalgamated into a single “Barren” cluster in Cresswell et al. (2017), while the clusters *red algae* and *other brown algae* further detail what Cresswell et al. (2017) identified as one single “Foliose algae” group. Thus, our large-scale spatial approach overall confirms the habitat types defined by Cresswell et al. (2017) while to some extent providing a more nuanced distinction between similar habitats. Nevertheless, our analysis only managed to capture one coral reef habitat state, contrary to what we might have expected.

**Figure 6.**
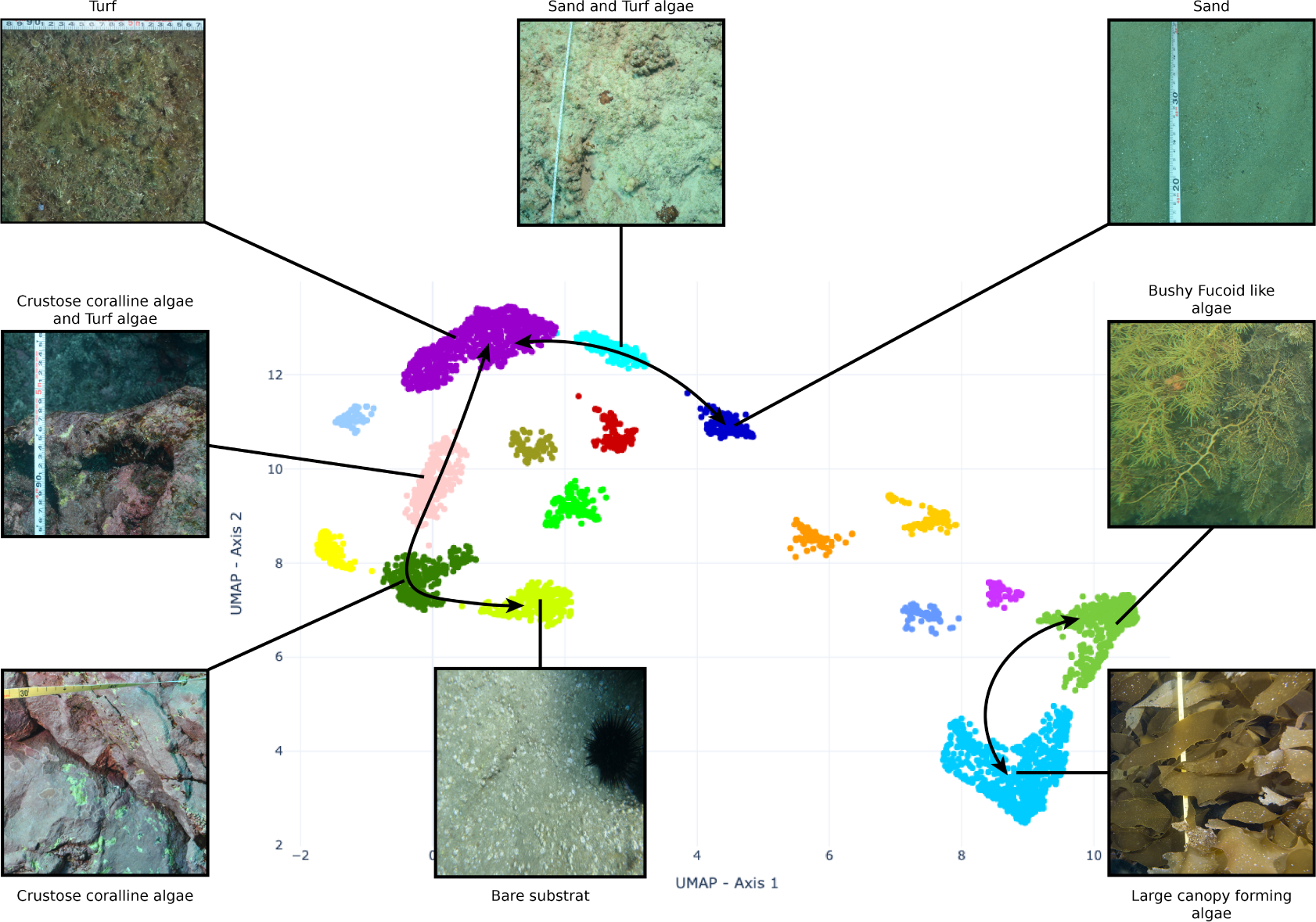
Two-dimensional *UMAP* embedding of the 5,090 clustered RLS transects. Each picture is representative of its cluster. The arrows indicate potential transitions as identified in Jouffray et al. (2015) and Cornwall et al. (2023).

There may be several reasons for only distinguishing a single coral reef state. First, the area covered by coral exhibits significant variability across reefs, as indicated by (De’ath et al. 2012), which can make it difficult to characterise coral reef state from cover data alone. Additionally, the morphological diversity of these reefs is extensive, with variations in surface areas (Zawada, Dornelas, and Madin 2019). Such variability might elucidate why a singular group of corals, *branching coral* is found as a group, since some species of *Acropora spp.* are able to establish colonies with expansive surface areas. Hence, our data-driven approach yields a finely resolved classification that comprises typical benthic habitats (e.g. seagrass meadows, coral reefs, kelp forests) that are common across all major seafloor habitat classification scheme (e.g. *European Nature Information System*; Bajjouk et al. (2015)).

Our global classification of the *RLS* data highlights hotspots of diversity in terms of benthic habitats and habitat states. Four ecoregions in particular, the Eastern (Manning-Hawkesbury ecore-gion) and Western Australia (Houtman ecoregion), the Caribbean, and the Tuamotus Archipelago, showed a high diversity of habitat types (considering both richness and evenness). In the transition zones between temperate and tropical waters, such as the Manning-Hawkesbury or the Houtman ecoregions in eastern and western Australia respectively, the high diversity of benthic habitat types we observe could be explained by a high diversity of foundation species. Indeed, high biodiversity is typical of transitional environmental conditions where ecological niches, which are overall disjointed, overlap (Ferro and Morrone 2014). This phenomenon is well known for multiple taxa such as birds (Altamirano et al. 2020), plants (Lemessa, Mewded, and Alemu 2023) or reef fish (Pinheiro et al. 2018) and also seems to apply to a certain extent to biogenic habitats like coralline red algae (Sissini et al. 2022). Such subtropical or warm temperate zones are also identified as regions where both mobile fauna (Vergés et al. 2014) or sessile habitat-forming species assemblages (Marzloff et al. 2018) are likely to undergo tropicalisation, which implies that native temperate assemblages can co-occur and being in competition with warmer species assemblages undergoing poleward climate-driven range shifts that could replace indigenous species. Our finely resolved classification could be modelled against environmental predictors in future work to understand and predict the state of benthic habitats under current and future conditions (e.g. Belanger et al. (2012)).

Beyond exploring spatial patterns of benthic biodiversity, our classification of the *RLS* dataset offers a new perspective to explore temporal changes in benthic habitat states at site level. In fact, the proportion of transects classified in the different groups seems to be an interesting indicator for researchers and managers, providing a metric for expressing the dominant ecological state, less affected by the inherent variability that transects can present. This metric can be particularly useful for identifying changes in the structure of ecosystems. One of the sites surveyed by (Rick D. Stuart-Smith et al. 2022), Beware Reef underwent a major ecological change following an overgrazing event in 2013 by Centrostephanus rodgersii sea urchins (N. Barrett et al. 2014). Our metric allows us to observe that the regime change towards a urchin barren state is not complete. In fact, this species of sea urchin is omnivorous, capable of attacking sessile invertebrates (Byrne and Andrew 2013), but showing a clear preference for algae, in particular *Ecklonia radiata*, a species belonging to the *large canopy forming algae* group (Hill et al. 2003). Our metric indicates that many transects at this site are classified as *red algae*, or o*ther sessile invertebrates*, showing that the proliferation of these urchins appears to be contained, in agreement with in-situ observations (Environmental Sustainability 2021).

Overall benthic habitat changes may reflect a range of processes, including ecological ones such as temporal variability in the cover of habitat-forming species (e.g. Wernberg et al. (2016)) in relation to climate-driven environmental changes (i.e. tropicalisation of tropical-temperate transition zones (Horta e Costa et al. 2014), marine heatwaves Wernberg et al. (2016)) or to trends in human stressors (i.e nutrients and organic pollution runoffs, impacts from coastal human populations; Halpern et al. (2019)), as well as methodological ones, such as variability in transect location or in sampling effort through time (e.g. Stuble et al. (2021)). Identifying the processes driving the observed habitat transitions could help better characterise the impact of anthropogenic activities on benthic habitats (see for example Donovan et al. (2018) for a similar approach at a finer spatial scale). Our classification could thus provide an interesting template to further explore changes in benthic habitats across the world with expanded monitoring efforts (Graham J. Edgar, Rick D. Stuart-Smith, et al. 2023).

Nonetheless, not all expected transitions between habitats, or alternative ecological states, come out in the different clusters. Some transitory states may be classified as noise if they are too scarcely observed in the dataset to constitute a cluster of their own. Understanding the drivers behind the transects classified as noise can reveal valuable information about the factors influencing habitat variability and the ecological processes driving shifts between different states. This includes deciphering the reasons for a noise classification, such as variations in environmental conditions, biotic interactions, or anthropogenic disturbances. By investigating these aspects, researchers can gain crucial insights into the dynamics and transitions occurring between habitat states and alternative ecological states.

The *UMAP-HDBSCAN* clustering pipeline presented in this study demonstrates remarkable robust-ness and versatility, leveraging global data to identify fine-scale patterns within coastal temperate and tropical ecosystems. Because of its hierarchical structure, this pipeline aligns well with estab-lished classification standards and facilitates a first data-driven description of global patterns in habitat states, which constitutes a valuable database to explore the influence of local and global drivers of benthic habitat states. Additionally, the pipeline’s capability to handle non-linear data and accommodate noise underscores its adaptability to various ecological contexts and data sources (e.g. citizen science).

## Supporting information

Supplementary Informations

## Acknowledgments

We are grateful to all volunteers and staff involved in the *Reef Life Survey* monitoring programme. The authors would also like to acknowledge the Pôle de Calcul et de Données Marines (PCDM) for providing DATARMOR storage and computational resources for this study. https://pcdm.ifremer.fr. MPM is the recipient of an ANR early career grant ANR-21-CE02-0006.

## Author Contributions

Clément Violet: Conceptualisation (lead), Methodology (lead), Visualisation (lead), Writing - original draft (lead), Writing - Review & Editing (lead)

Aurélien Boyé: Conceptualisation (lead), Methodology (equal), Writing - original draft (equal), Writing - Review & Editing (equal)

Stanislas Dubois: Conceptualisation (equal), Writing - original draft (equal), Writing - Review & Editing (equal)

Graham J. Edgar: Investigation (lead), Data curation(lead), Writing - original draft (equal), Writing - Review & Editing (equal)

Elizabeth S. Oh: Investigation (lead), Data curation(lead), Writing - original draft (equal), Writing - Review & Editing (equal)

Rick D. Stuart-Smith: Investigation (lead), Data curation (lead) Writing - original draft (equal), Writing - Review & Editing (equal)

Martin P. Marzloff: Conceptualisation (lead), Methodology (equal), Writing - original draft (equal)

