## Supplementary Informations for "From local seafloor imagery to global patterns in benthic habitat states: contribution of citizen science to habitat classification across latitudes"

### Appendix A

Table 1: Correspondence of the functional groups of substrates and the habitat groups used in this study.

| Photoquadrat Original Categories | Final Classification |
| --- | --- |
| Ahermatypic corals | Sessile invertebrates |
| Ascidians | Sessile invertebrates |
| Ascidians (stalked) | Sessile invertebrates |
| Ascidians (unstalked) | Sessile invertebrates |
| Bare Rock | Bare substrate |
| Barnacles | Sessile invertebrates |
| Black & Octocorals | Soft corals and gorgonians |
| Bottlebrush Acropora corals | Branching corals |
| Bottlebrush Acropora corals_Bleached | Branching corals |
| Branching Acropora | Branching corals |
| Branching Acropora_Bleached | Branching corals |
| Branching corals | Branching corals |
| Branching corals_Bleached | Branching corals |
| Branching Pocillopora | Branching corals |
| Branching Pocillopora_Bleached | Branching corals |
| Bryozoa | Sessile invertebrates |
| Bryozoan (hard) | Sessile invertebrates |
| Bryozoan (soft) | Sessile invertebrates |
| Caulerpa | Green algae |
| Cnidaria | Sessile invertebrates |
| Cobble | Unconsolidated substrate |
| Colonial Anemones, Zoanthids and Corallimorphs | Sessile invertebrates |
| Columnar corals | Massive corals |
| Columnar corals_Bleached | Massive corals |
| Coral rubble | Unconsolidated substrate |
| Coral rubble with turf/encrusting algae | Turf |
| Corymbose Acropora corals | Branching corals |
| Corymbose Acropora corals_Bleached | Branching corals |
| Crustose coralline algae | Crustose coralline algae |
| Dead coral | Bare substrate |
| Desmarestia and Himantothallus | Canopy forming algae |
| Digitate corals | Massive corals |
| Digitate corals_Bleached | Massive corals |
| Durvillaea | Canopy forming algae |
| Ecklonia radiata | Canopy forming algae |
| Encrusting corals | Encrusting corals |
| Encrusting corals_Bleached | Encrusting corals |
| Encrusting leathery algae | Encrusting leathery algae |
| Filamentous algae_epiphyte | Filamentous algae |
| Filamentous brown algae_epiphyte | Filamentous algae |
| Filamentous green algae_epiphyte | Filamentous algae |
| Filamentous red algae_epiphyte | Filamentous algae |
| Filamentous rock-attached algae | Filamentous algae |
| Foliose/Plate corals | Foliose/Plate corals |
| Foliose/Plate corals_Bleached | Foliose/Plate corals |
| Geniculate coralline algae | Geniculate coralline algae |
| Green calcified algae (Halimeda) | Green calcified algae (Halimeda) |
| Heliopora coerulea (blue coral) | Calcareous hydrocorals and octocorals |
| Hydrocoral | Calcareous hydrocorals and octocorals |
| Hydrocoral_Bleached | Calcareous hydrocorals and octocorals |
| Hydroids | Sessile invertebrates |
| Large brown laminarian kelps | Canopy forming algae |
| Large-polyp stony corals (free-living) | Large-polyp stony corals (free-living) |
| Large-polyp stony corals (free-living)_Bleached | Large-polyp stony corals (free-living) |
| Macroalgae | Canopy forming algae |
| Macroalgae_canopy forming | Canopy forming algae |
| Macrocystis | Canopy forming algae |
| Massive corals | Massive corals |
| Massive corals_Bleached | Massive corals |
| Medium foliose brown algae | Brown algae |
| Medium foliose green algae | Green algae |
| Medium foliose red algae | Red algae |
| Molluscs | Sessile invertebrates |
| Organ-pipe coral (Tubipora) | Calcareous hydrocorals and octocorals |
| Other fucoids | Canopy forming algae |
| Pebbles/gravel/shell | Unconsolidated substrate |
| Phyllospora | Canopy forming algae |
| Polychaete | Sessile invertebrates |
| Sand | Unconsolidated substrate |
| Seagrass (Halophila) | Seagrass |
| Seagrass (straplike) | Seagrass |
| Seagrasses | Seagrass |
| Sessile bivalves | Sessile invertebrates |
| Sessile gastropods | Sessile invertebrates |
| Slime (not trapping sediment) | Bare substrate |
| Small <2cm foliose algal cover (not trapping sediment) | Turf |
| Soft corals and gorgonians | Soft corals and gorgonians |
| Solitary Anemones | Sessile invertebrates |
| Sponges | Sessile invertebrates |
| Sponges (encrusting) | Sessile invertebrates |
| Sponges (erect) | Sessile invertebrates |
| Sponges (hollow) | Sessile invertebrates |
| Sponges (massive) | Sessile invertebrates |
| Stony corals | Encrusting corals |
| Stony corals_Bleached | Encrusting corals |
| Sub-massive corals | Massive corals |
| Sub-massive corals_Bleached | Massive corals |
| Substrate | Bare substrate |
| Tabular Acropora corals | Foliose/Plate corals |
| Tabular Acropora corals_Bleached | Foliose/Plate corals |
| Turfing algae (<2 cm high algal/sediment mat on rock) | Turf |
| Worms | Sessile invertebrates |

### Appendix B

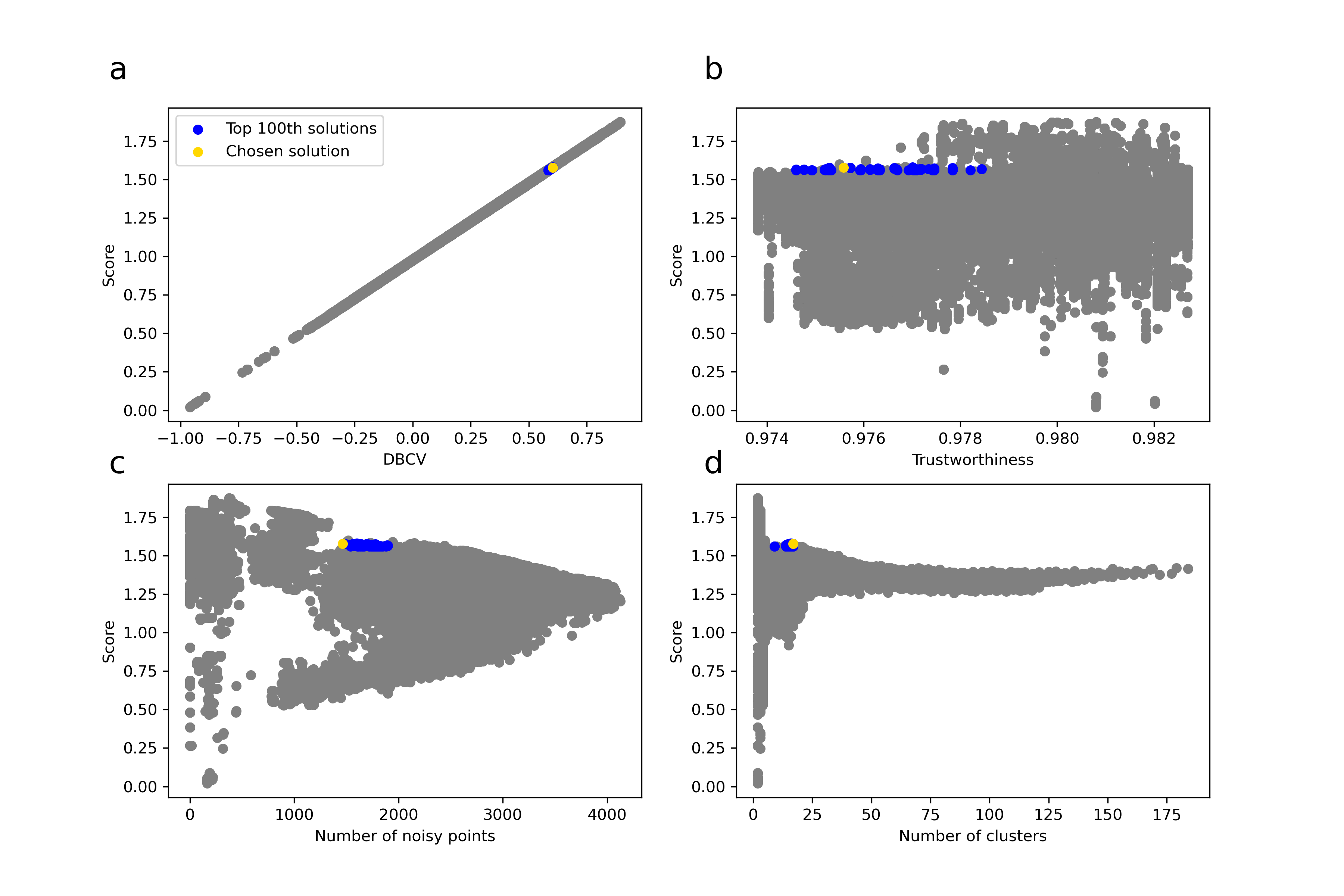

Figure 1: Plot of the evolution of the score (i.e. sum of the DBCV value and the Trustworthiness) according to the different parameters measured (a. DBCV; b. Trustworthiness; c. Number of noisy points; d. Number of clusters). Each of the 241,100 dots represent the evaluation of a unique combination of hyperparameter values for UMAP (n_neighbors) and HDBSCAN (min_cluster_size). Blue dots represent the top 100th solution, while the yellow dot represents the best solution found according to our criteria. All other grey dots represent the other evaluated solutions

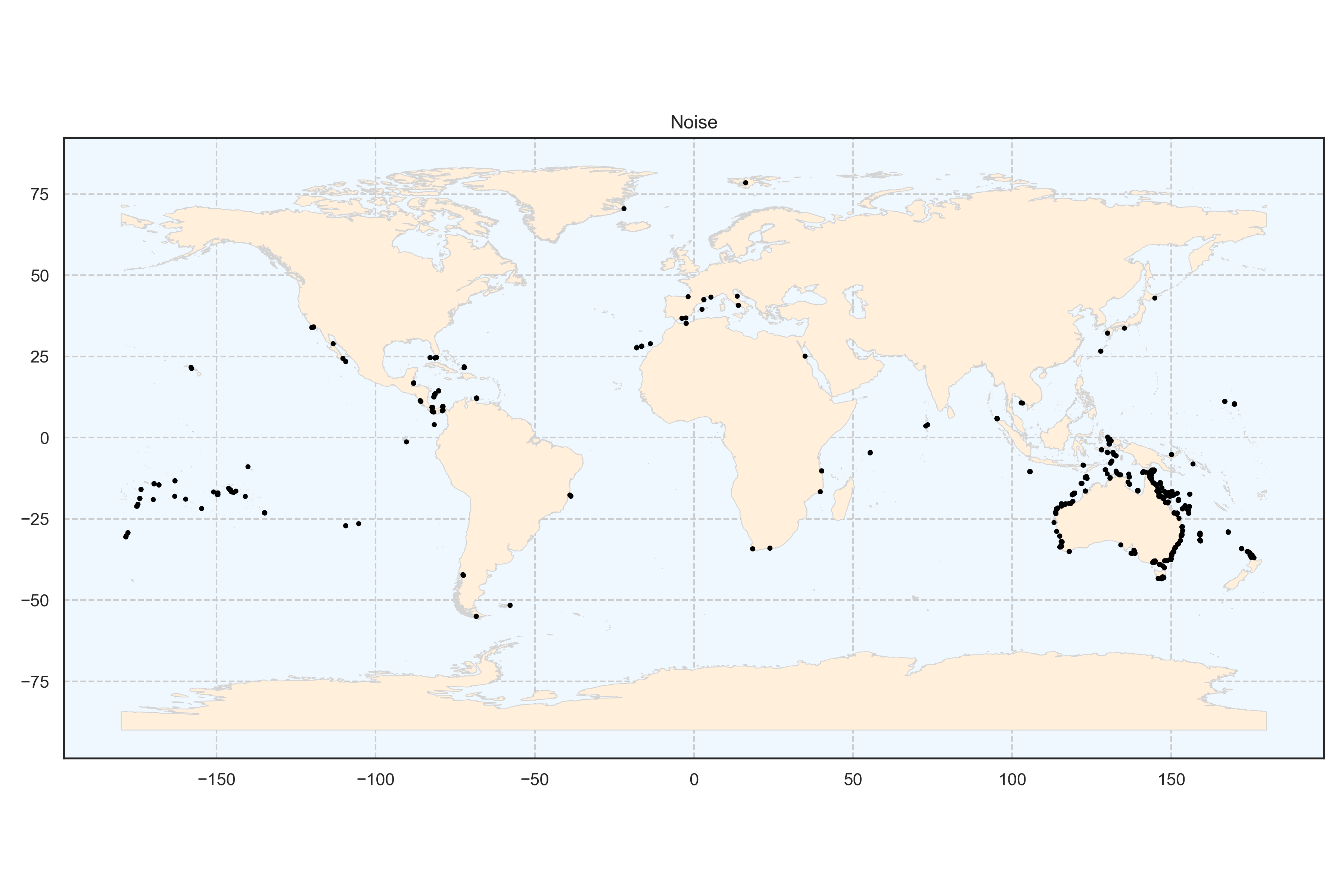

Figure 2: Spatial distribution of the cluster noise at the global scale. Each point represents a transect.

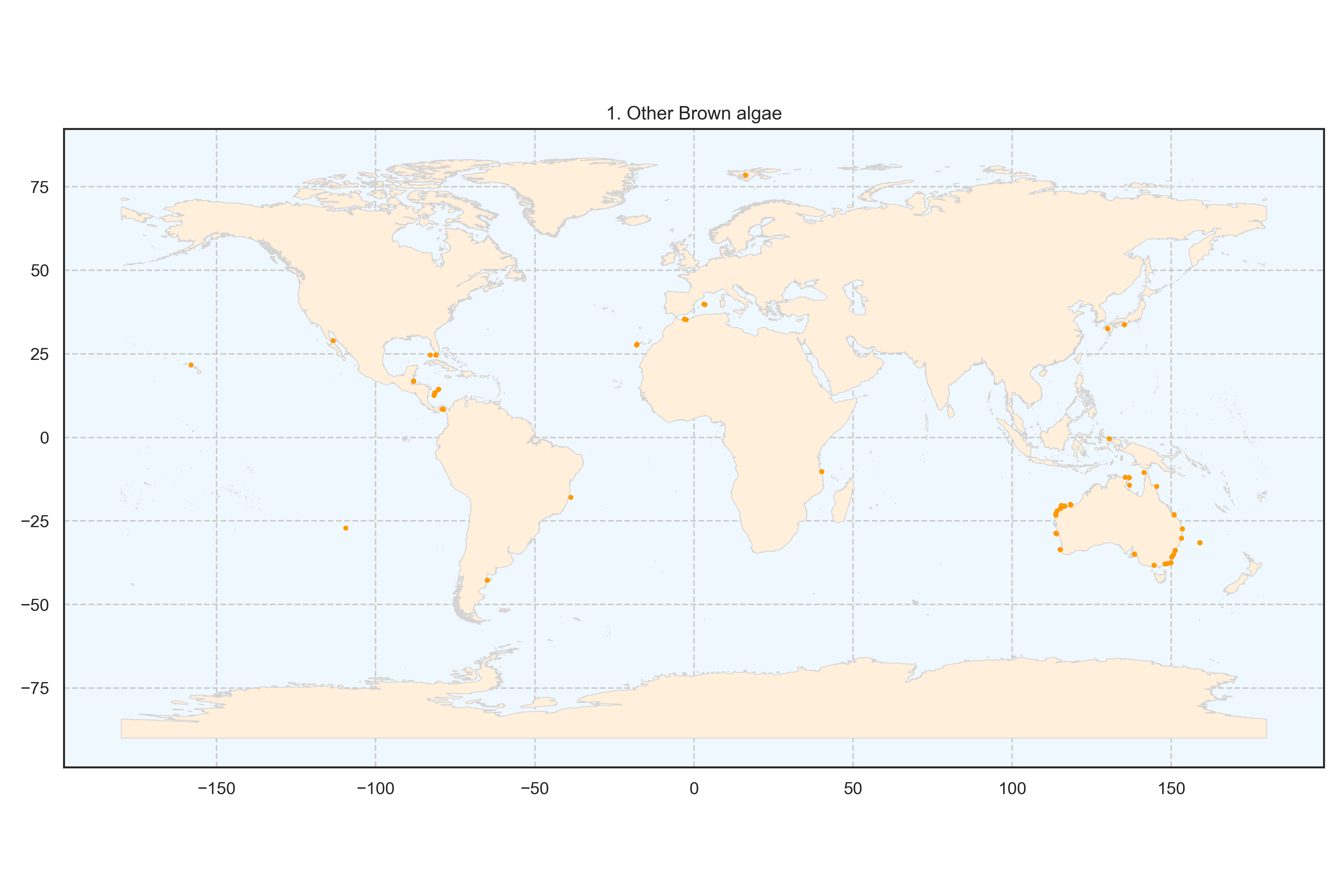

Figure 3: Spatial distribution of the cluster foliose brown algae at the global scale. Each point represents a transect.

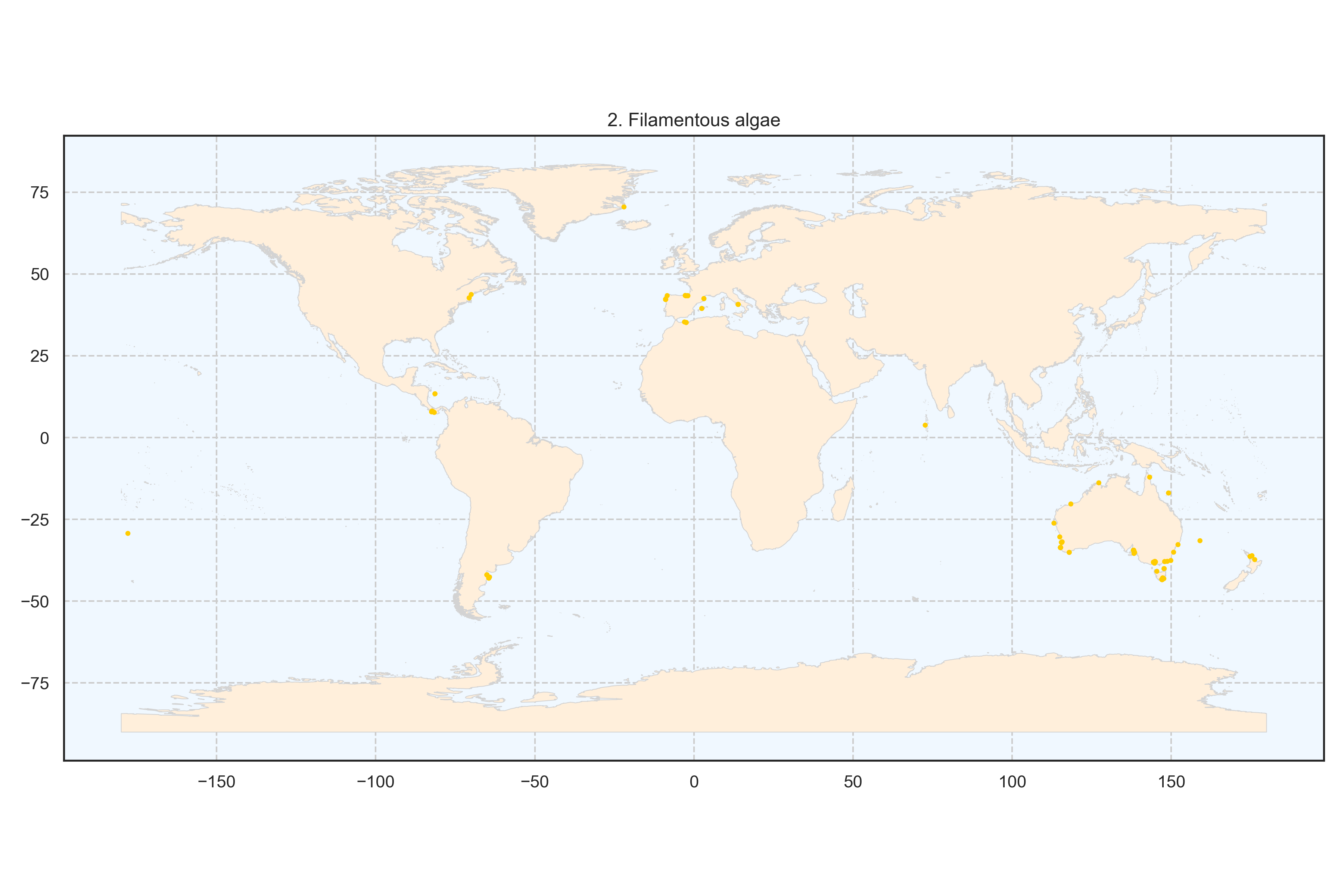

Figure 4: Spatial distribution of the cluster filamentous algae at the global scale. Each point represents a transect.

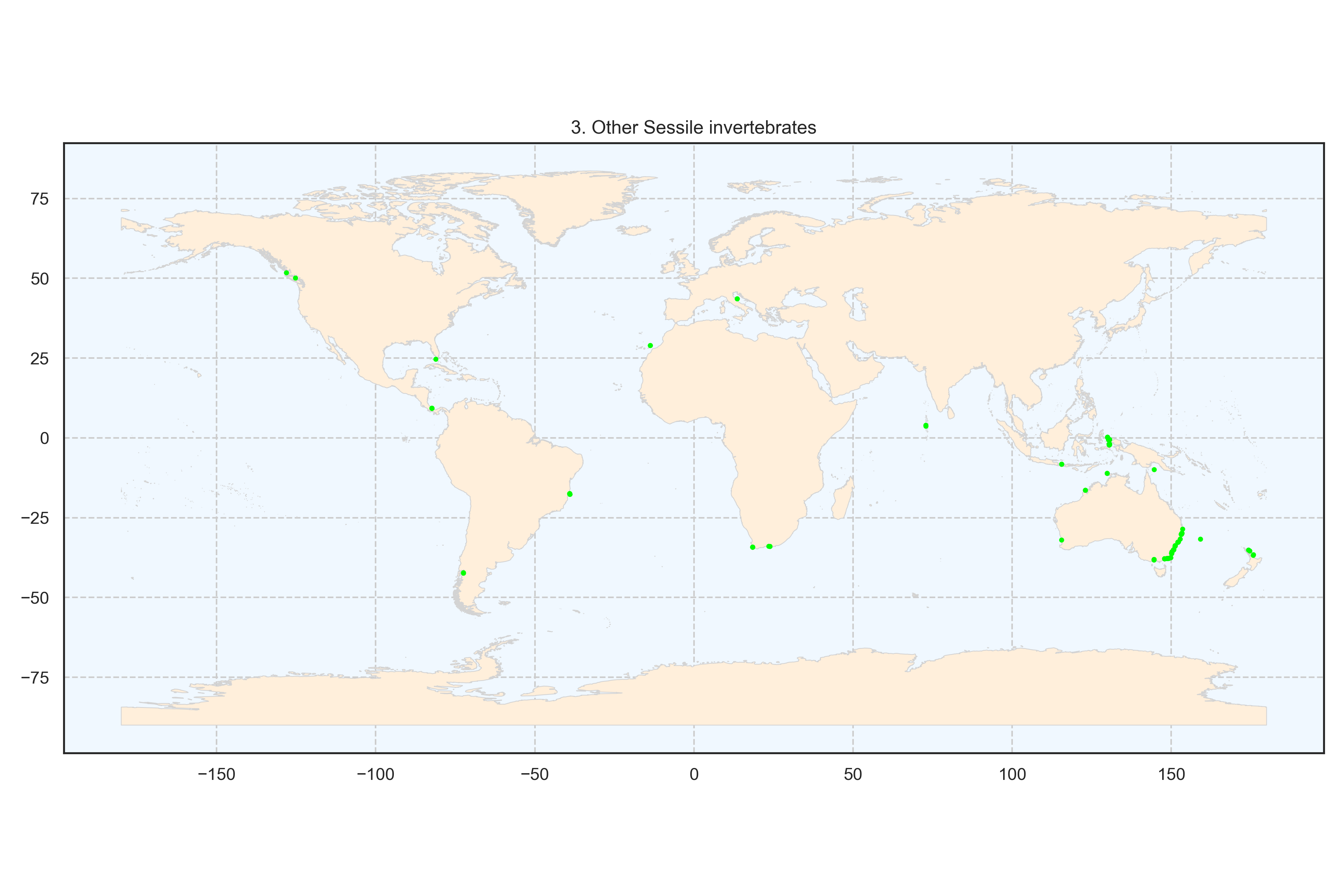

Figure 5: Spatial distribution of the cluster other Sessile invertebrates at the global scale. Each point represents a transect.

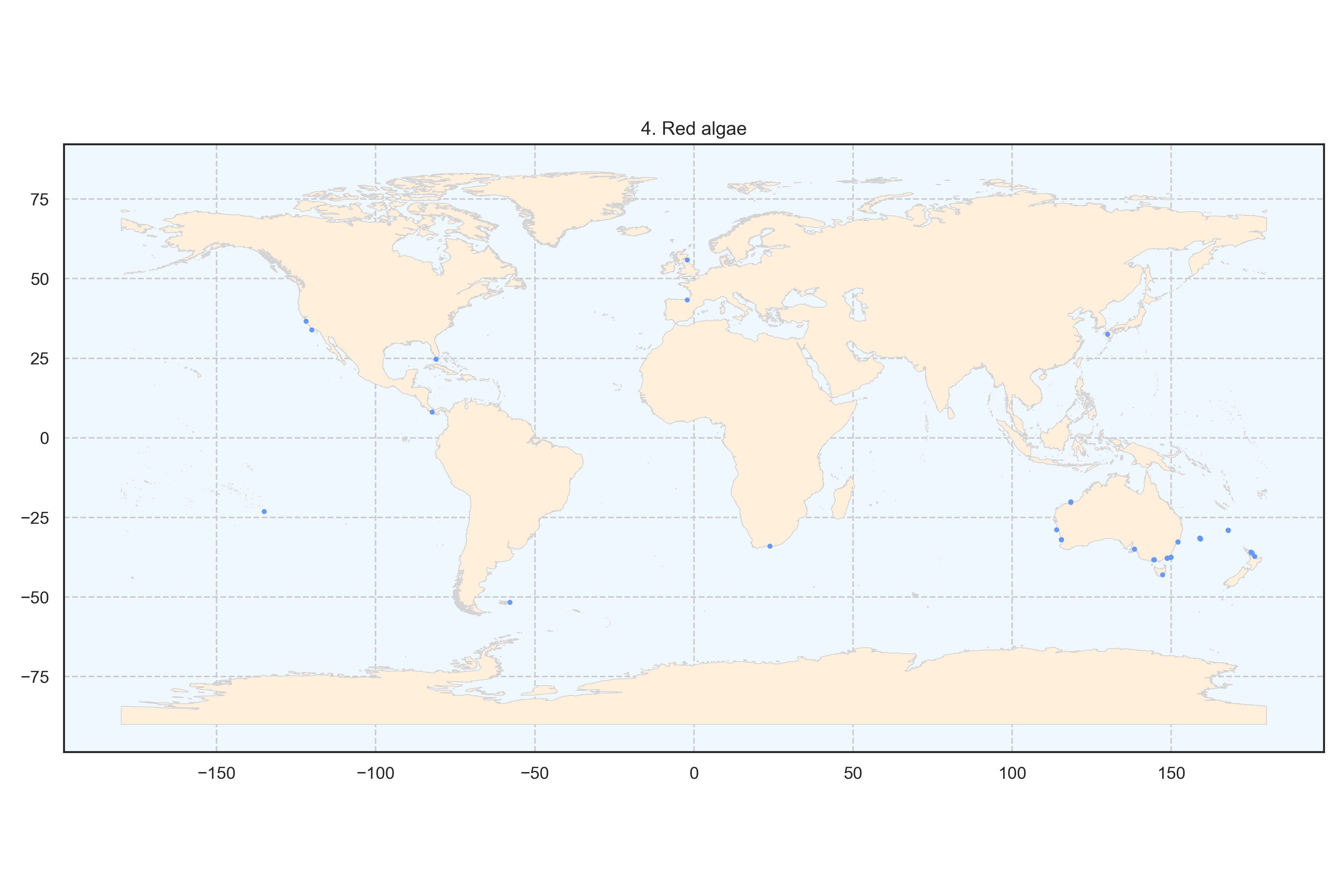

Figure 6: Spatial distribution of the cluster red algae at the global scale. Each point represents a transect.

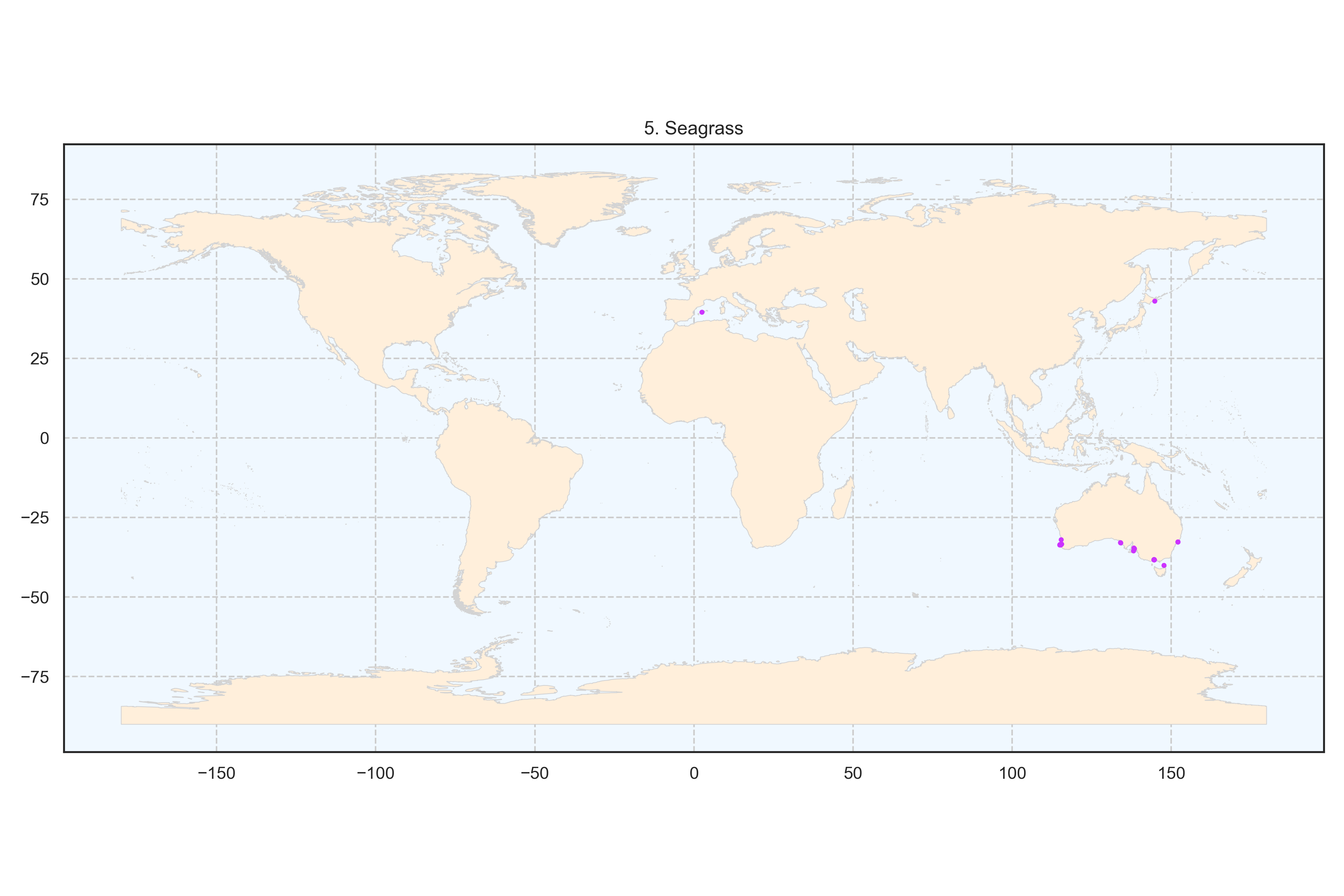

Figure 7: Spatial distribution of the cluster seagrass at the global scale. Each point represents a transect.

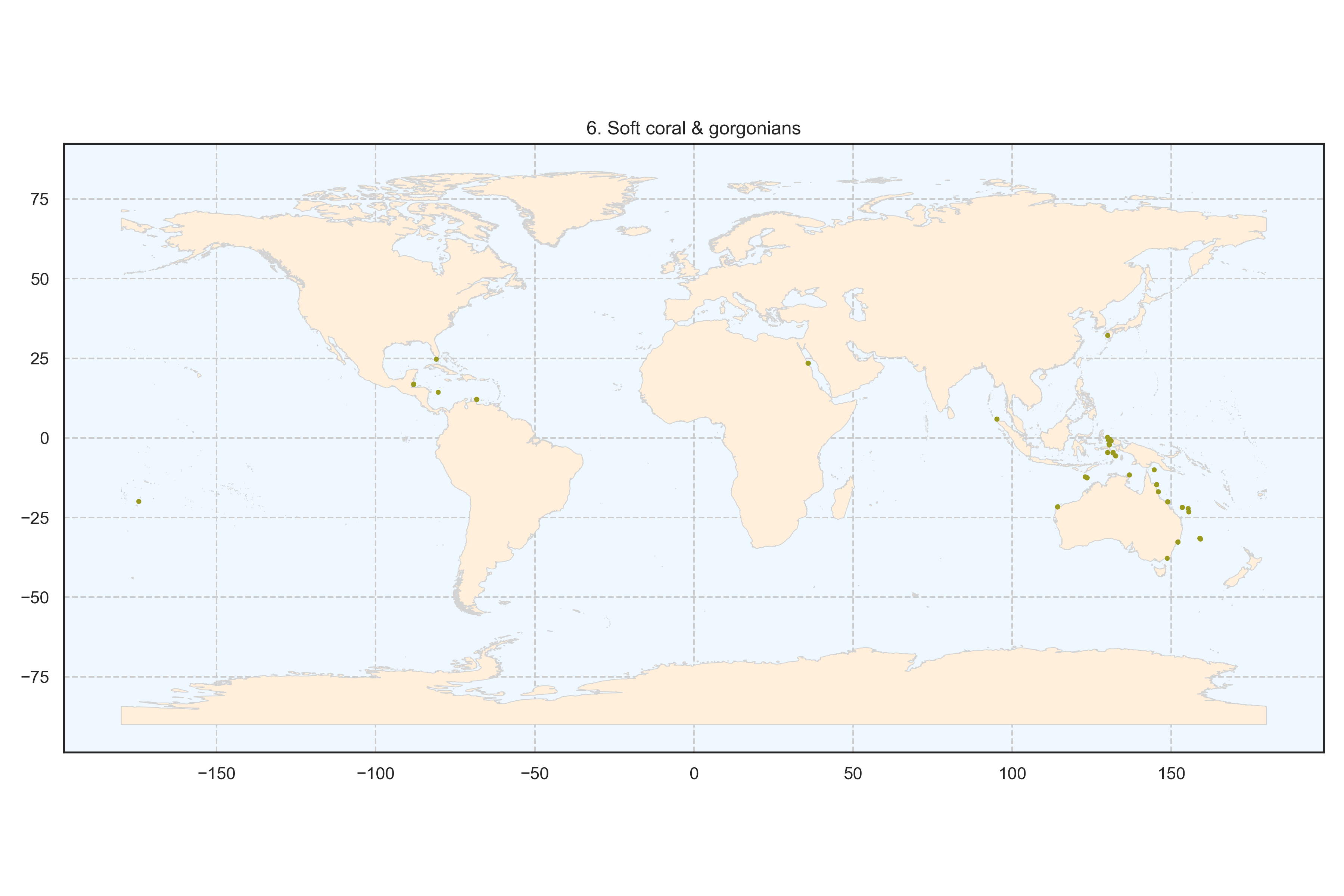

Figure 8: Spatial distribution of the cluster soft coral and gorgonians at the global scale. Each point represents a transect.

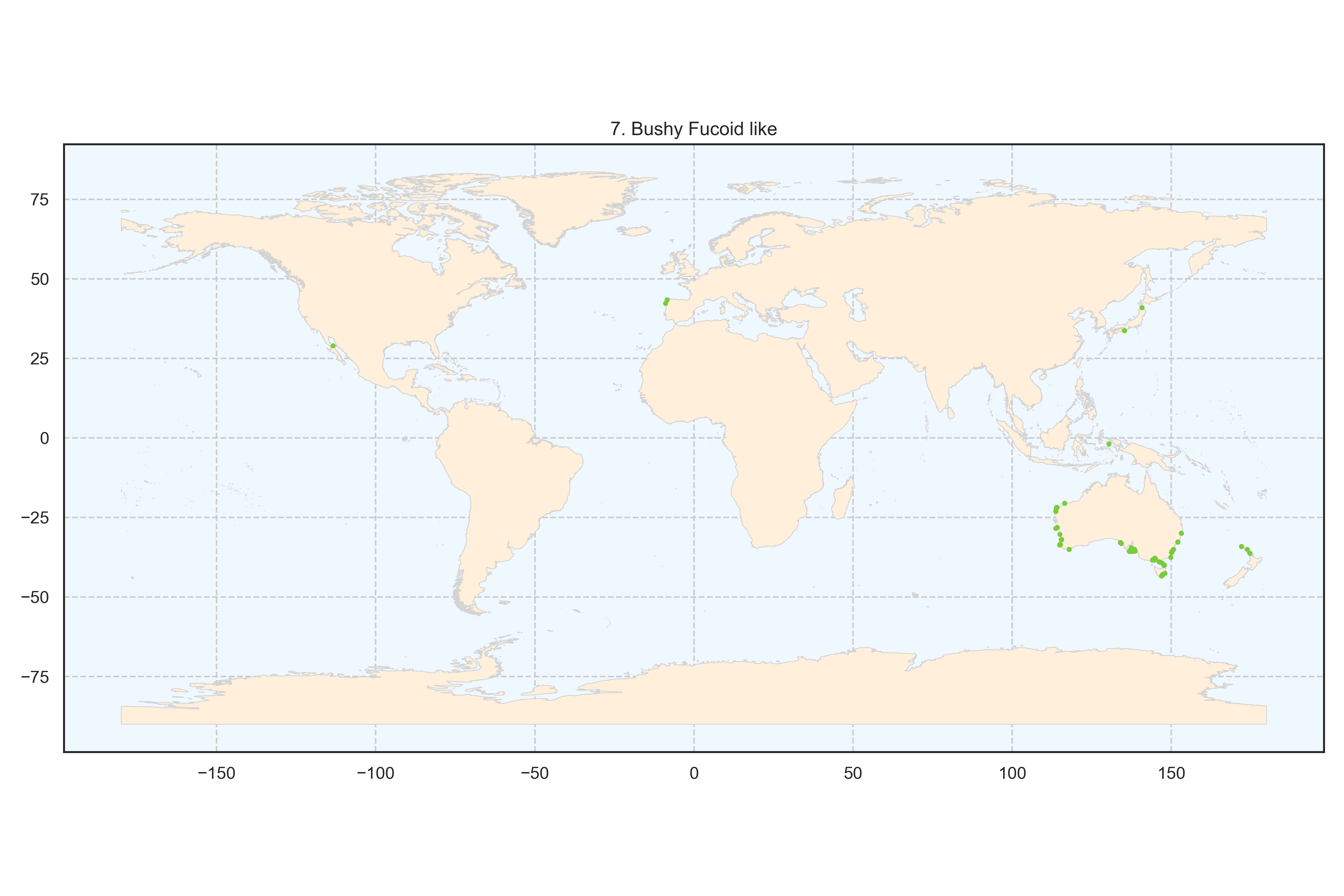

Figure 9: Spatial distribution of the cluster bushy fucoid-like algae at the global scale. Each point represents a transect.

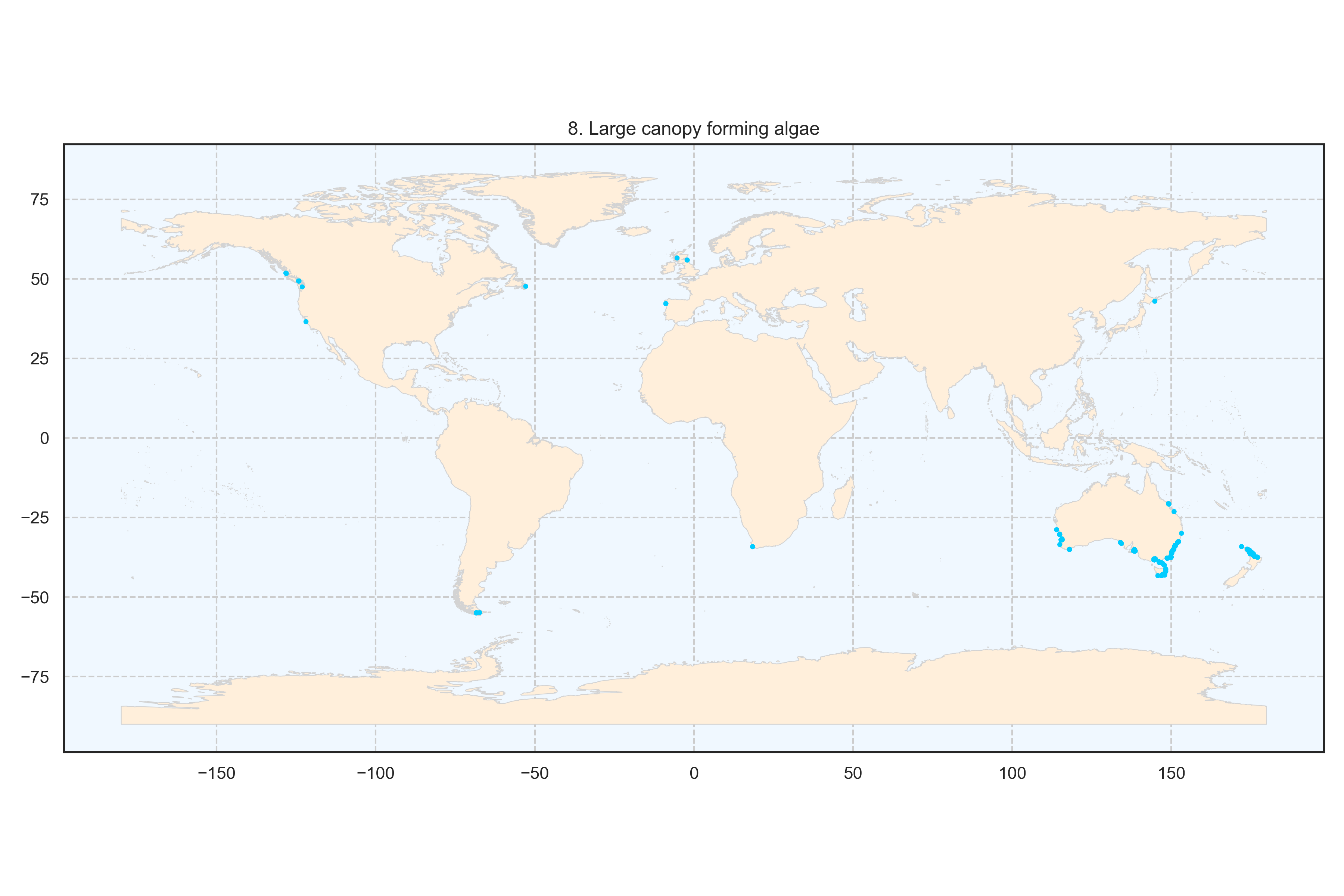

Figure 10: Spatial distribution of the cluster large canopy forming algae at the global scale. Each point represents a transect.

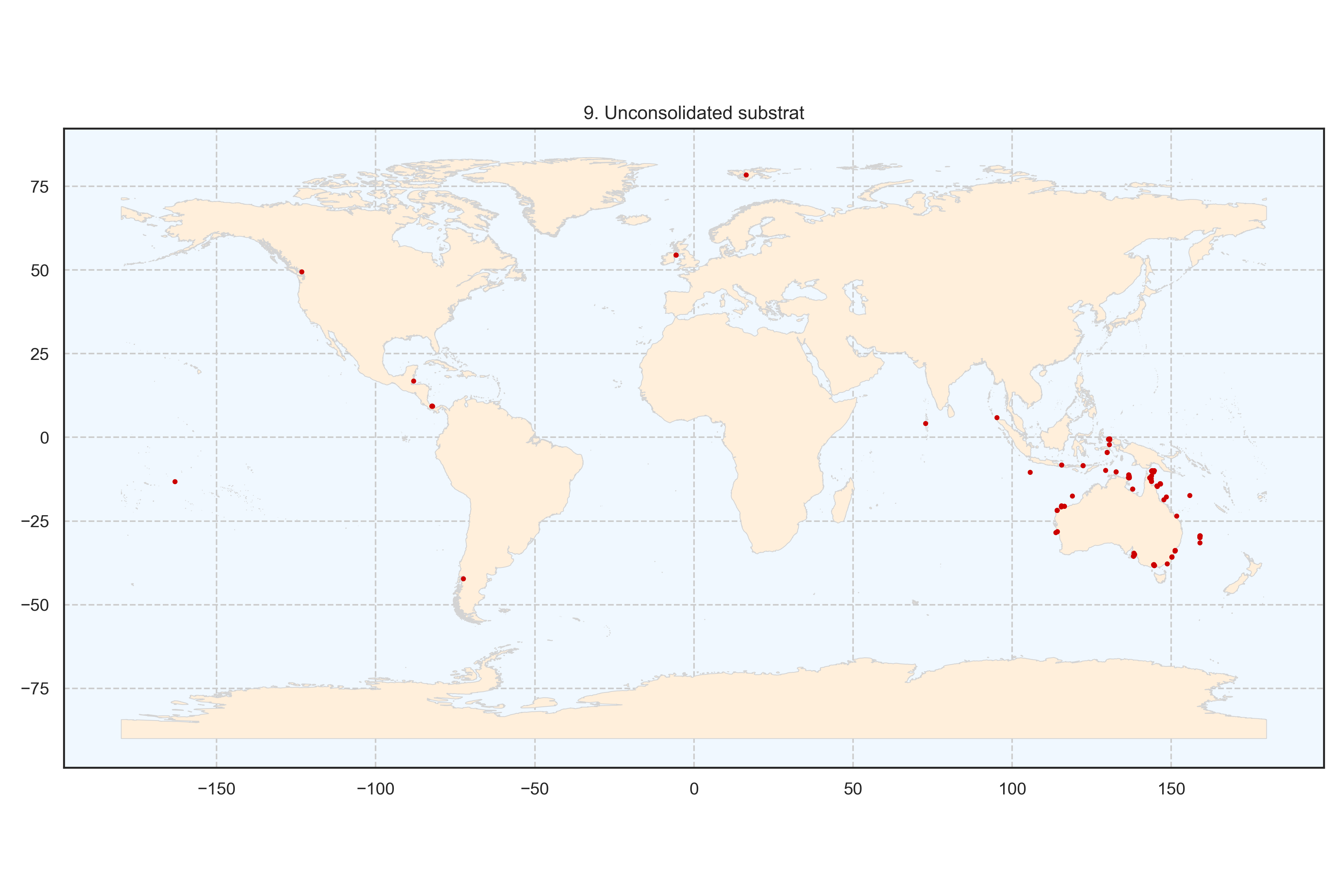

Figure 11: Spatial distribution of the cluster unconsolidated substrate at the global scale. Each point represents a transect.

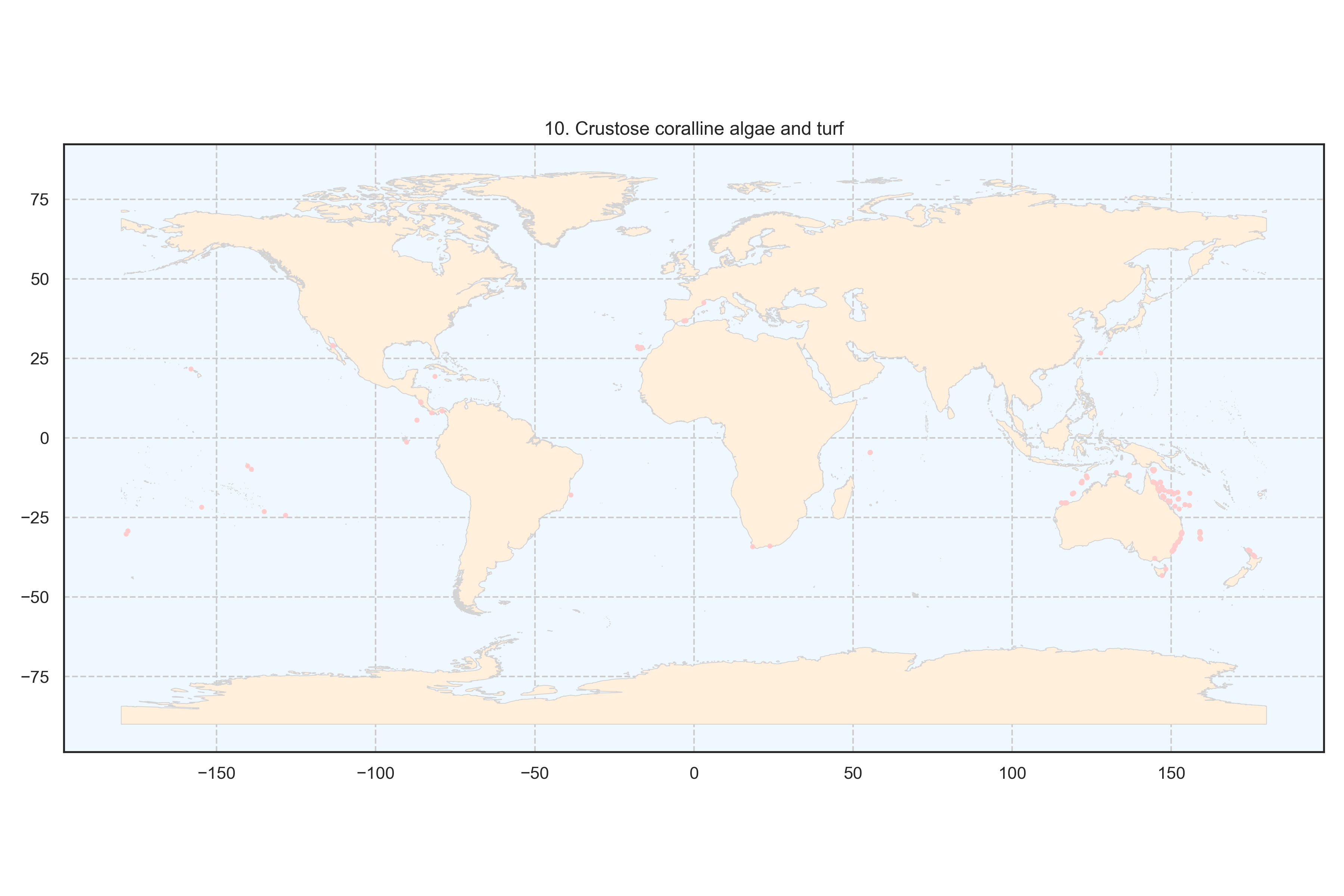

Figure 12: Spatial distribution of the cluster crustose coralline algae and turf at the global scale. Each point represents a transect.

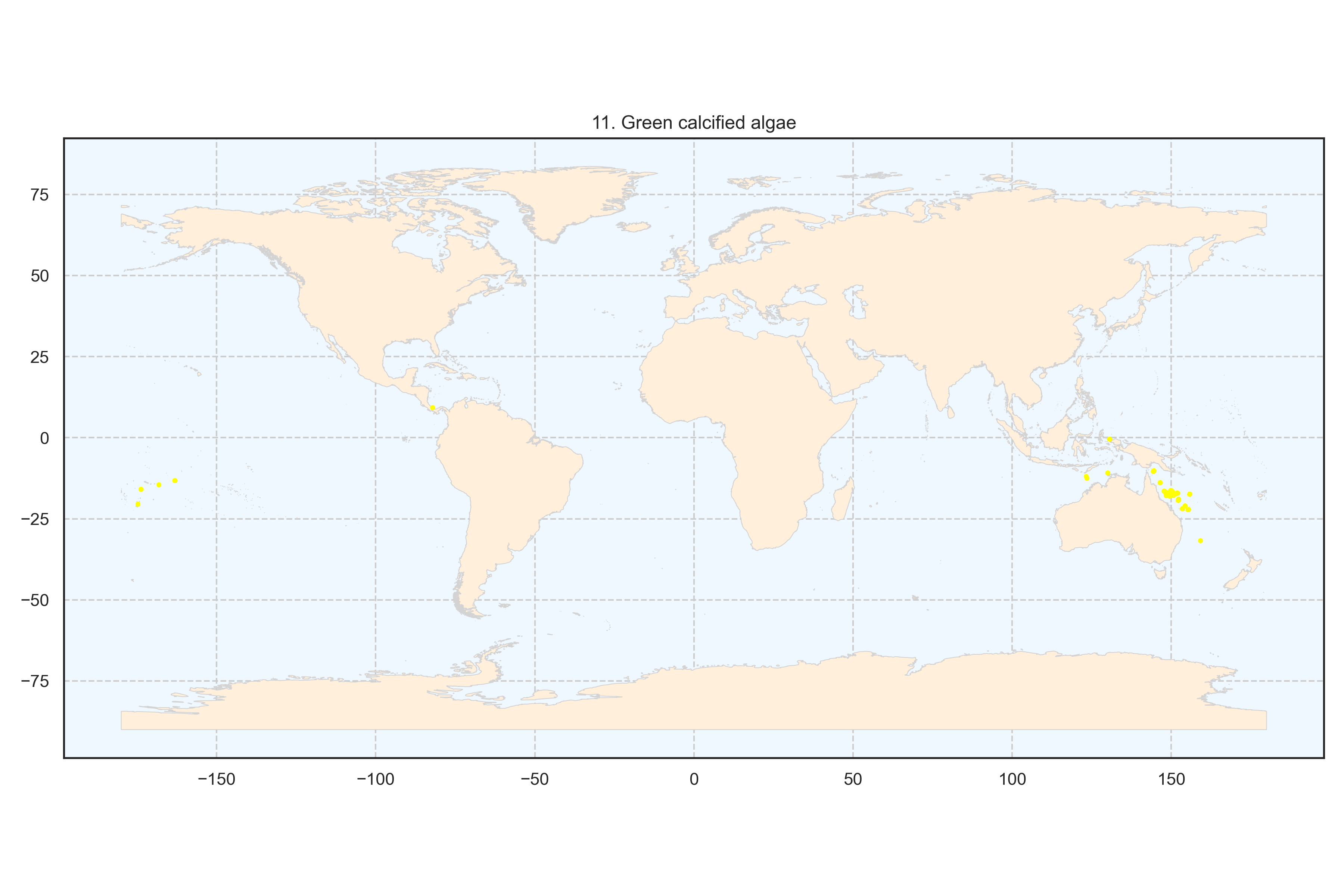

Figure 13: Spatial distribution of the cluster green calcified algae at the global scale. Each point represents a transect.

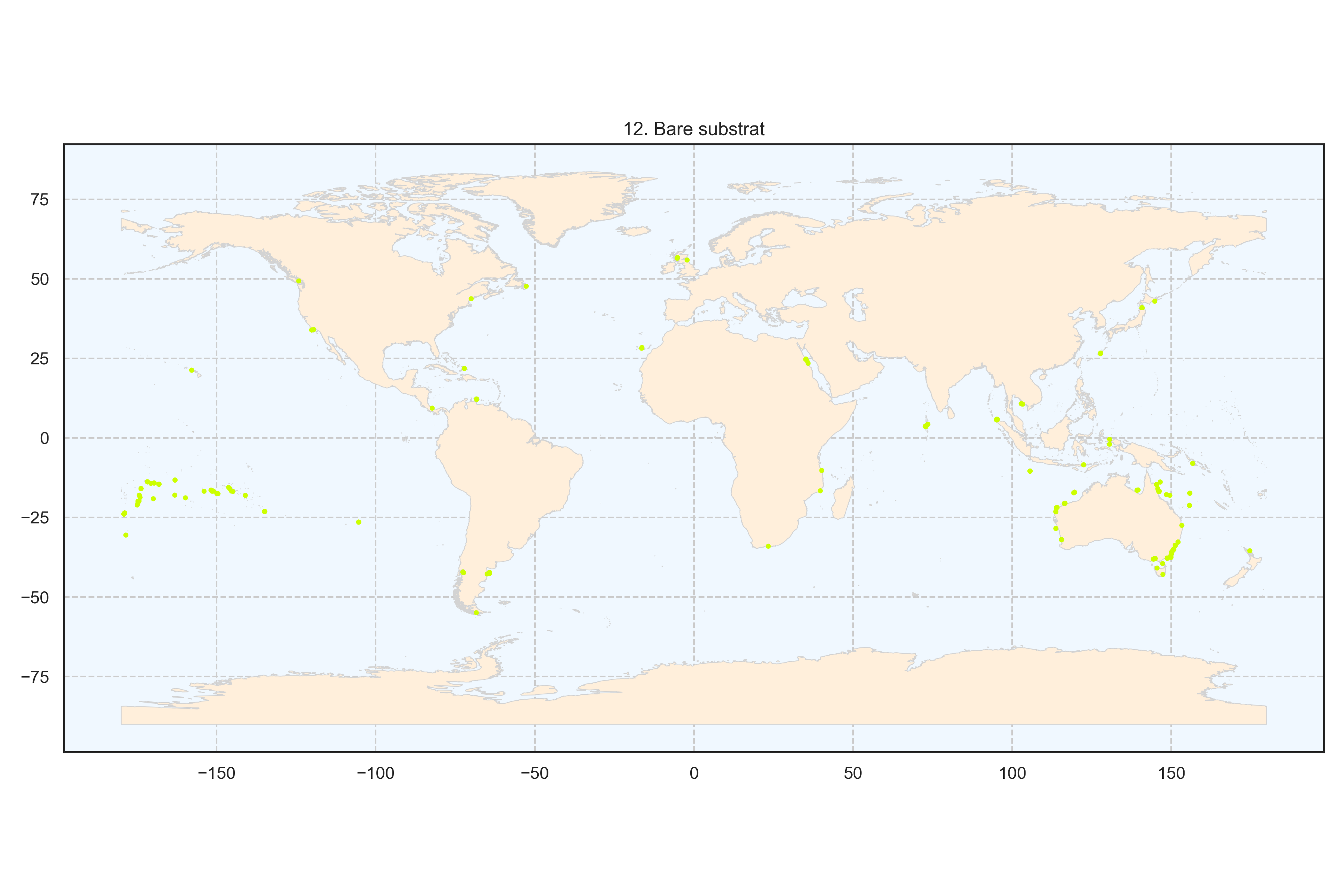

Figure 14: Spatial distribution of the cluster bare substrate at the global scale. Each point represents a transect.

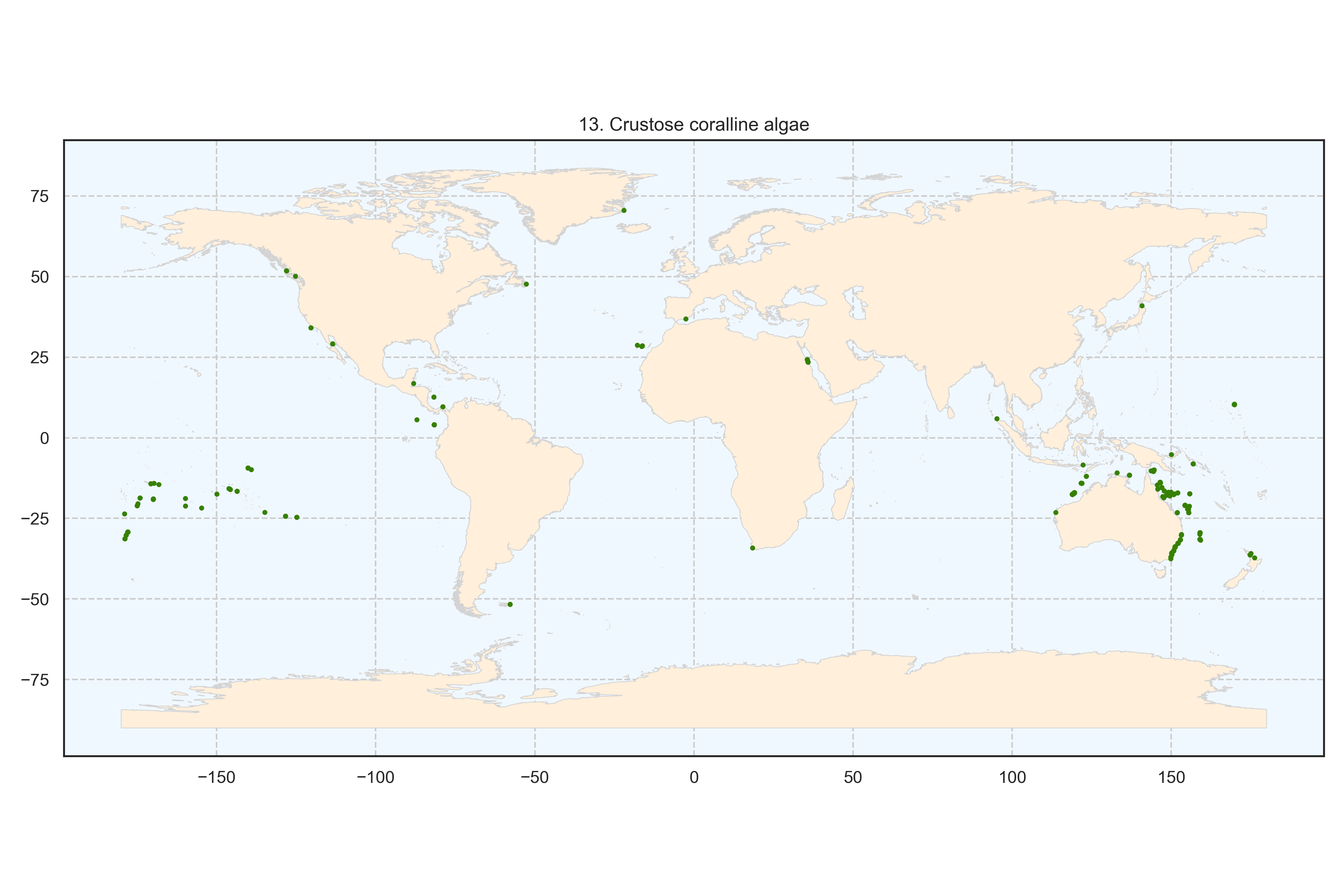

Figure 15: Spatial distribution of the cluster crustose coralline algae at the global scale. Each point represents a transect.

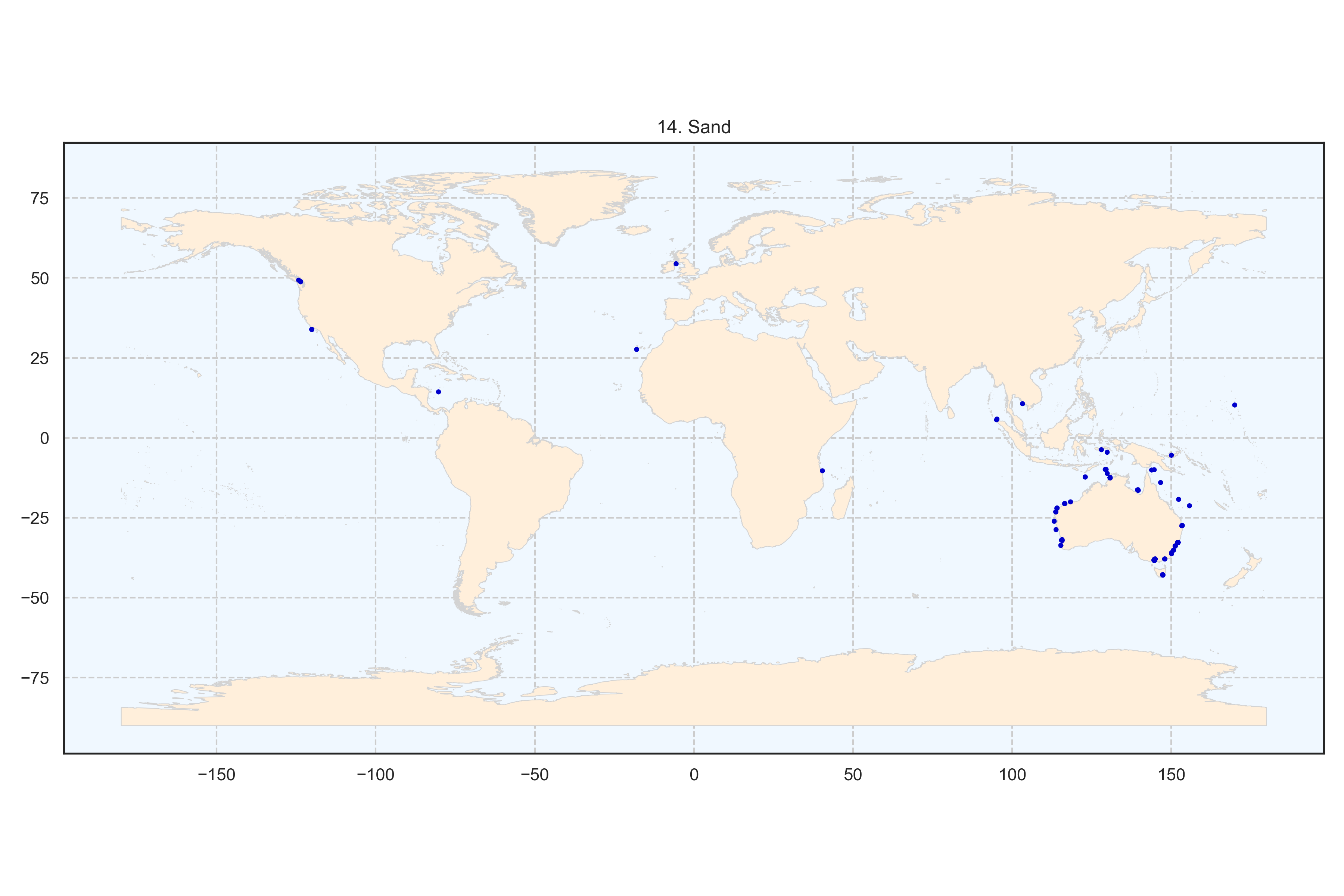

Figure 16: Spatial distribution of the cluster sand at the global scale. Each point represents a transect.

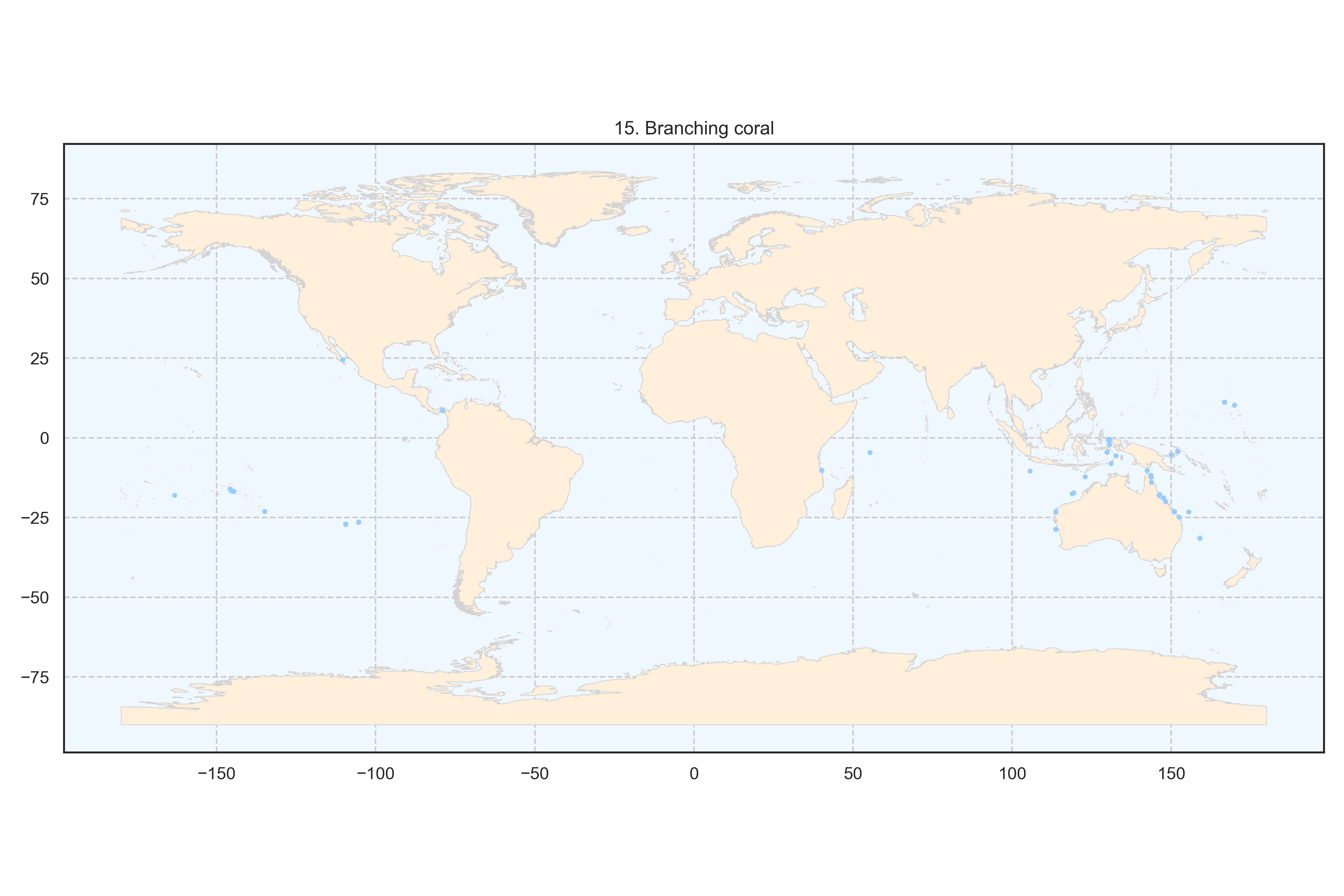

Figure 17: Spatial distribution of the cluster branching coral at the global scale. Each point represents a transect.

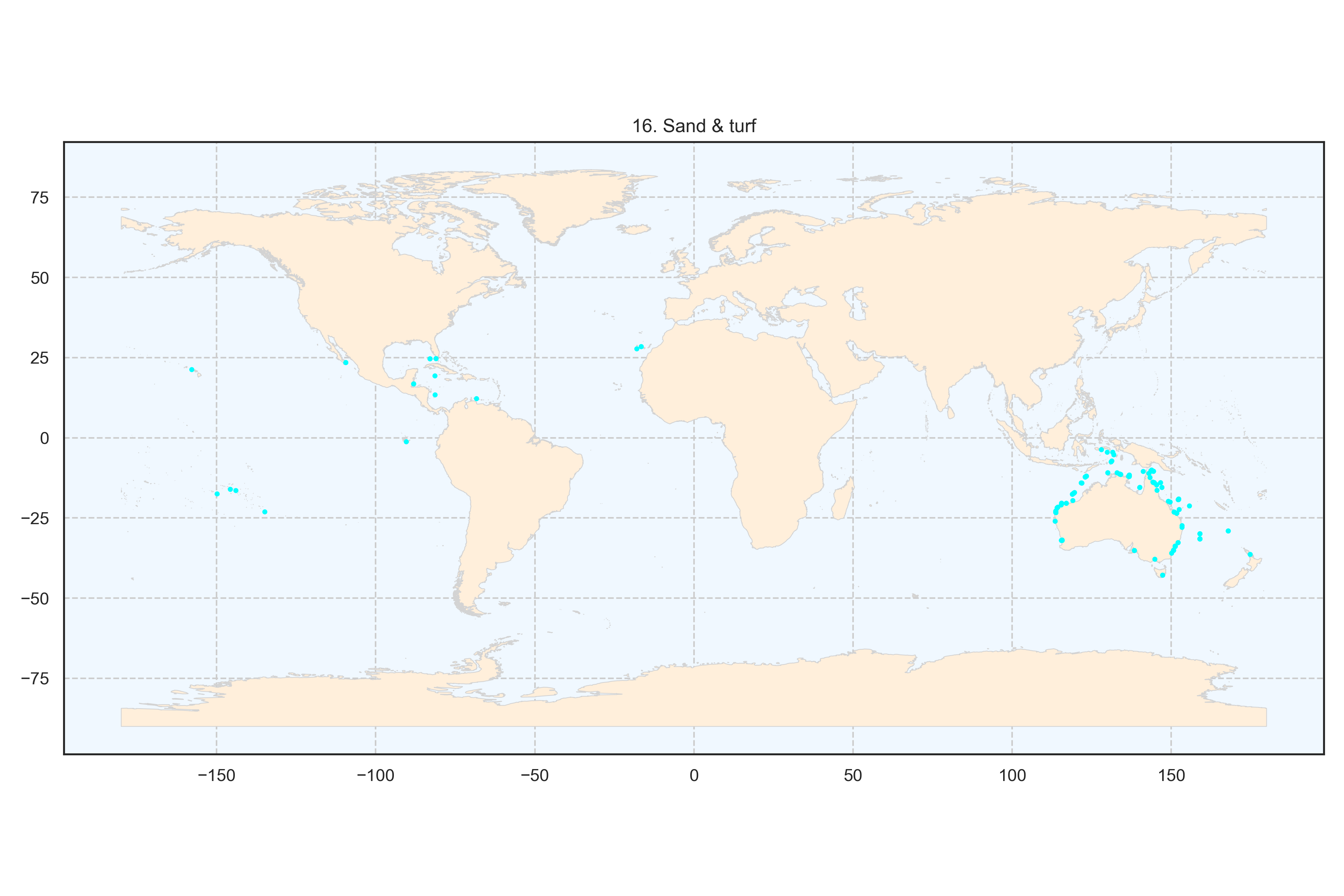

Figure 18: Spatial distribution of the cluster sand and turf algae at the global scale. Each point represents a transect.

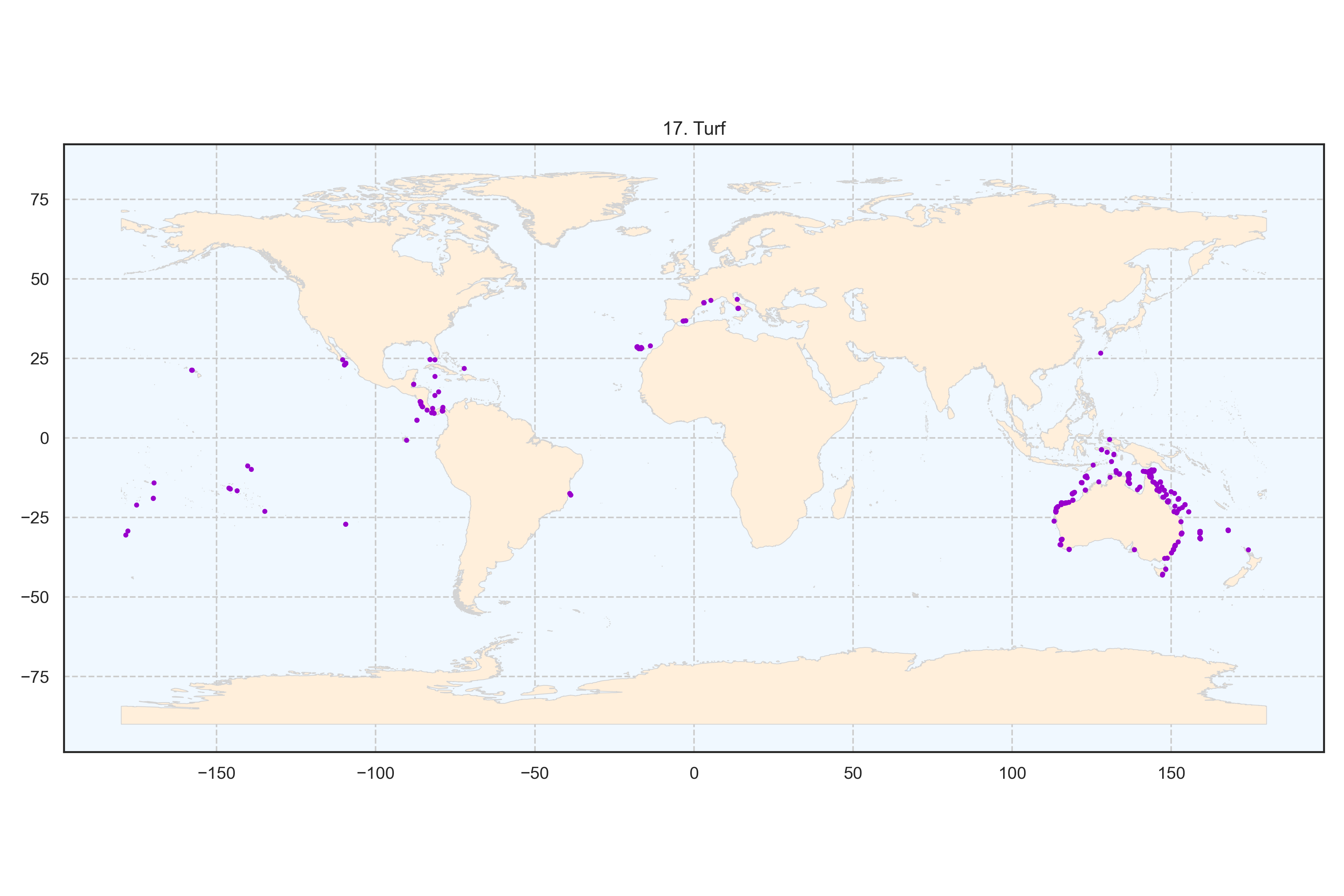

Figure 19: Spatial distribution of the cluster turf algae at the global scale. Each point represents a transect.

### Appendix C

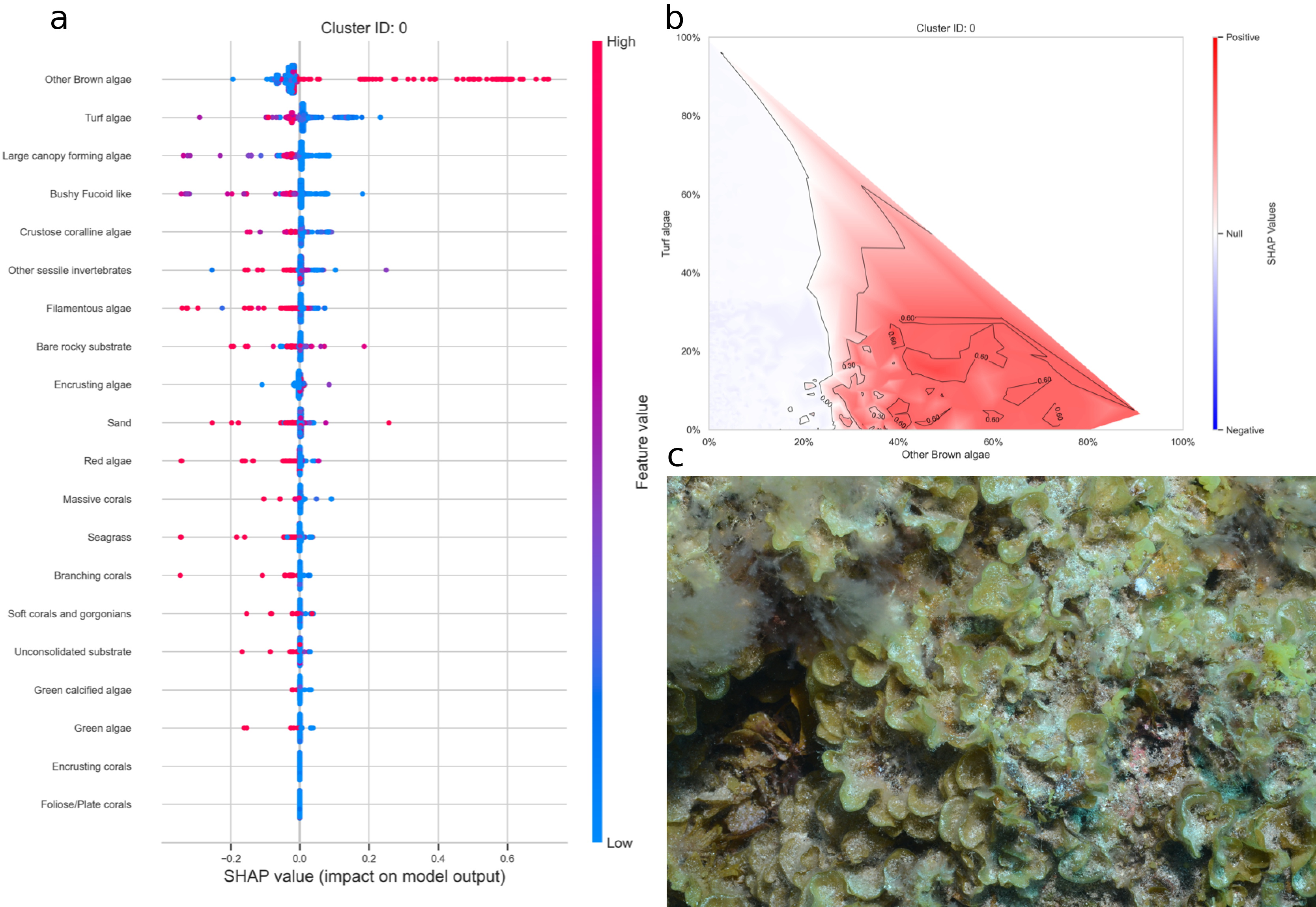

Figure 20: a. SHAP summary plot showing the impact of each habitat substrate on the classification. The position of each habitat group on the y-axis indicates its relative importance for the considered cluster. The position of each point along the x-axis indicates if the observation is associated with a lower or higher affinity with the cluster and the colour of the point indicates if the value of the cover of this habitat is rather high or low. b. Linear interpolation of the SHAP values for the two most influential variables for the cluster brown algae. c. Example of photoquadrat for one transect of the cluster brown algae categorised by HDBSCAN as exemplary.

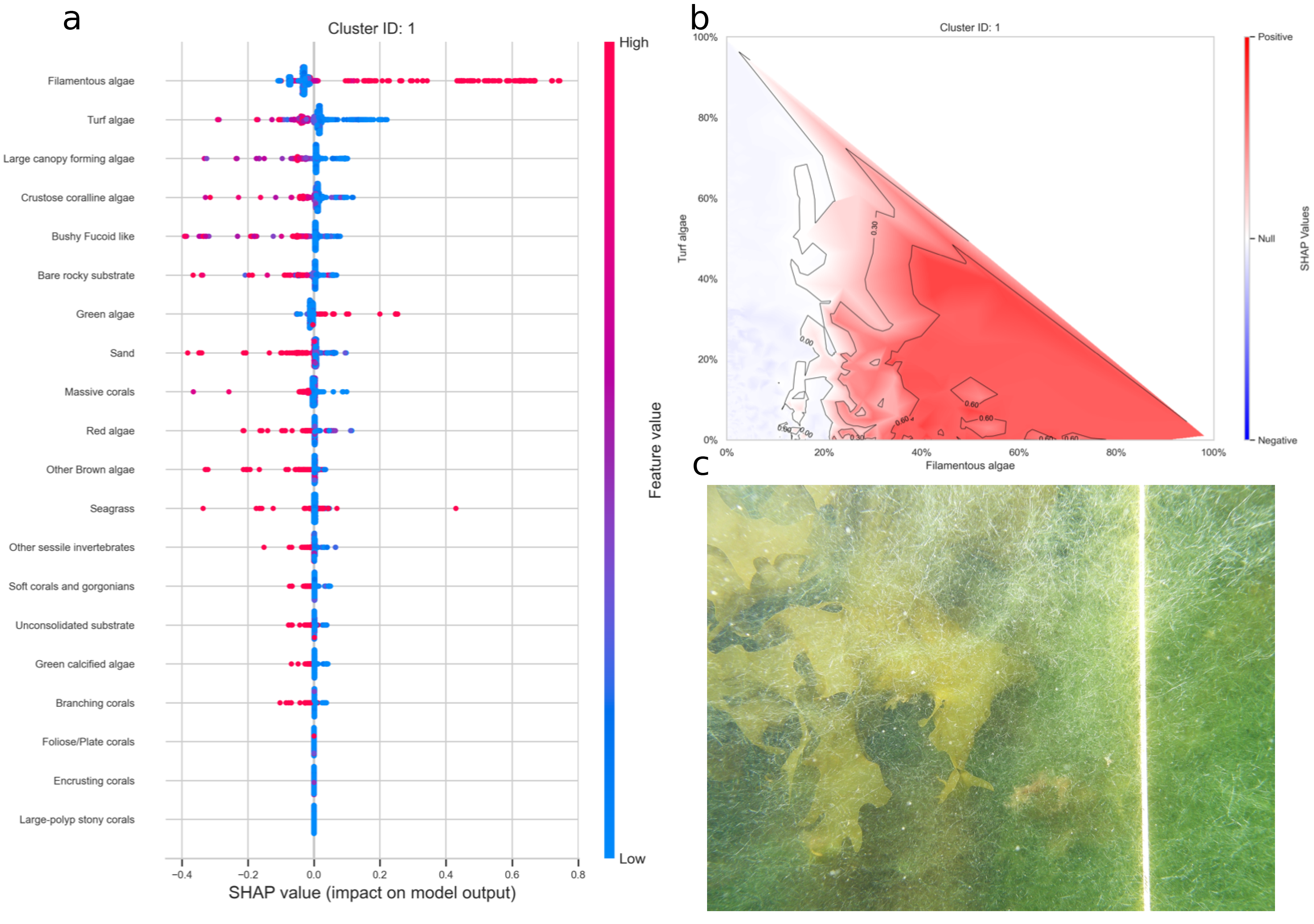

Figure 21: a. SHAP summary plot showing the impact of each habitat substrate on the classification. The position of each habitat group on the y-axis indicates its relative importance for the considered cluster. The position of each point along the x-axis indicates if the observation is associated with a lower or higher affinity with the cluster and the colour of the point indicates if the value of the cover of this habitat is rather high or low. b. Linear interpolation of the SHAP values for the two most influential variables for the cluster filamentous algae. c. Example of photoquadrat for one transect of the cluster filamentous algae categorised by HDBSCAN as exemplary.

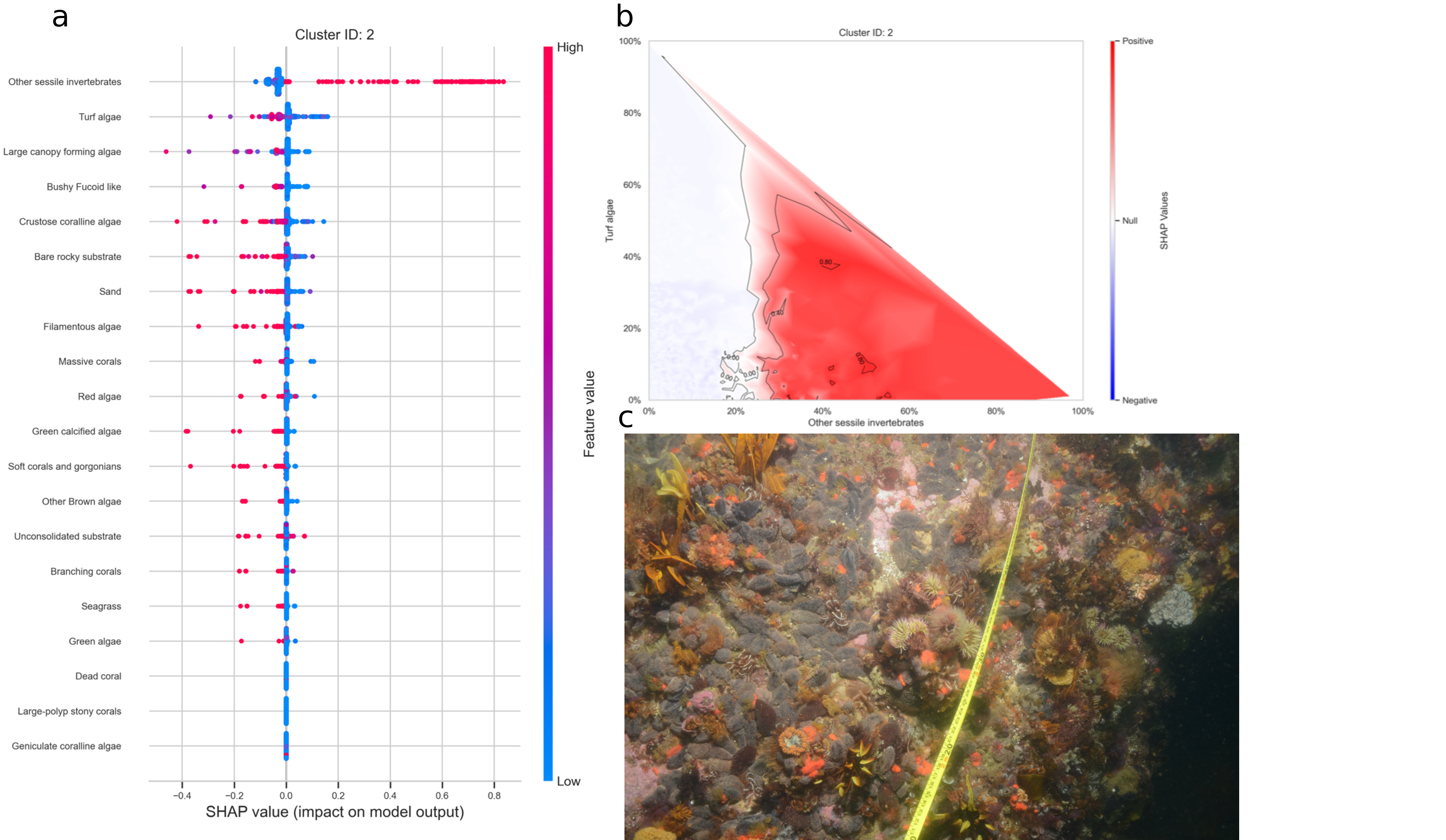

Figure 22: a. SHAP summary plot showing the impact of each habitat substrate on the classification. The position of each habitat group on the y-axis indicates its relative importance for the considered cluster. The position of each point along the x-axis indicates if the observation is associated with a lower or higher affinity with the cluster and the colour of the point indicates if the value of the cover of this habitat is rather high or low. b. Linear interpolation of the SHAP values for the two most influential variables for the cluster sessile invertebrates. c. Example of photoquadrat for one transect of the cluster sessile invertebrates categorised by HDBSCAN as exemplary.

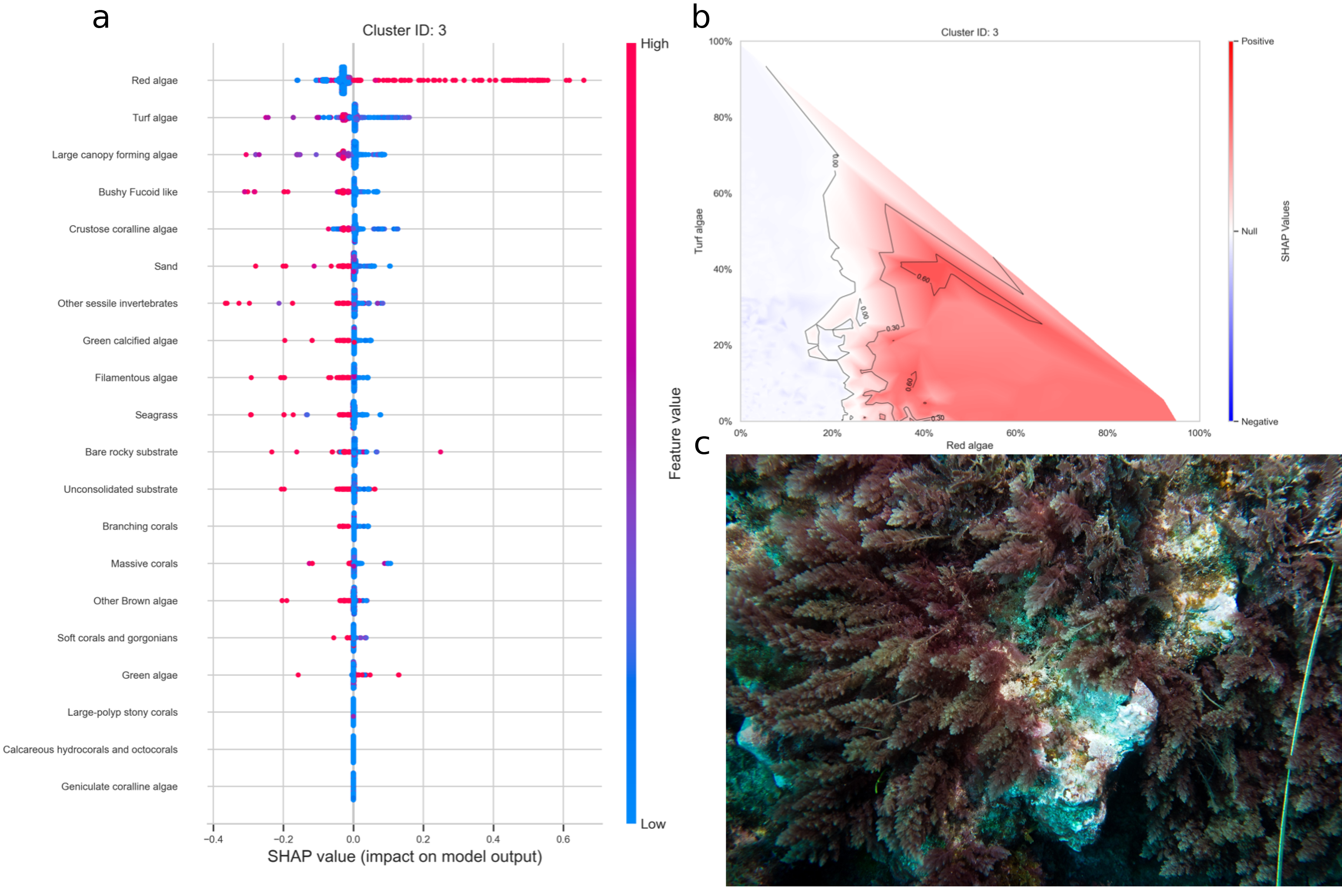

Figure 23: a. SHAP summary plot showing the impact of each habitat substrate on the classification. The position of each habitat group on the y-axis indicates its relative importance for the considered cluster. The position of each point along the x-axis indicates if the observation is associated with a lower or higher affinity with the cluster and the colour of the point indicates if the value of the cover of this habitat is rather high or low. b. Linear interpolation of the SHAP values for the two most influential variables for the cluster red algae. c. Example of photoquadrat for one transect of the cluster red algae categorised by HDBSCAN as exemplary.

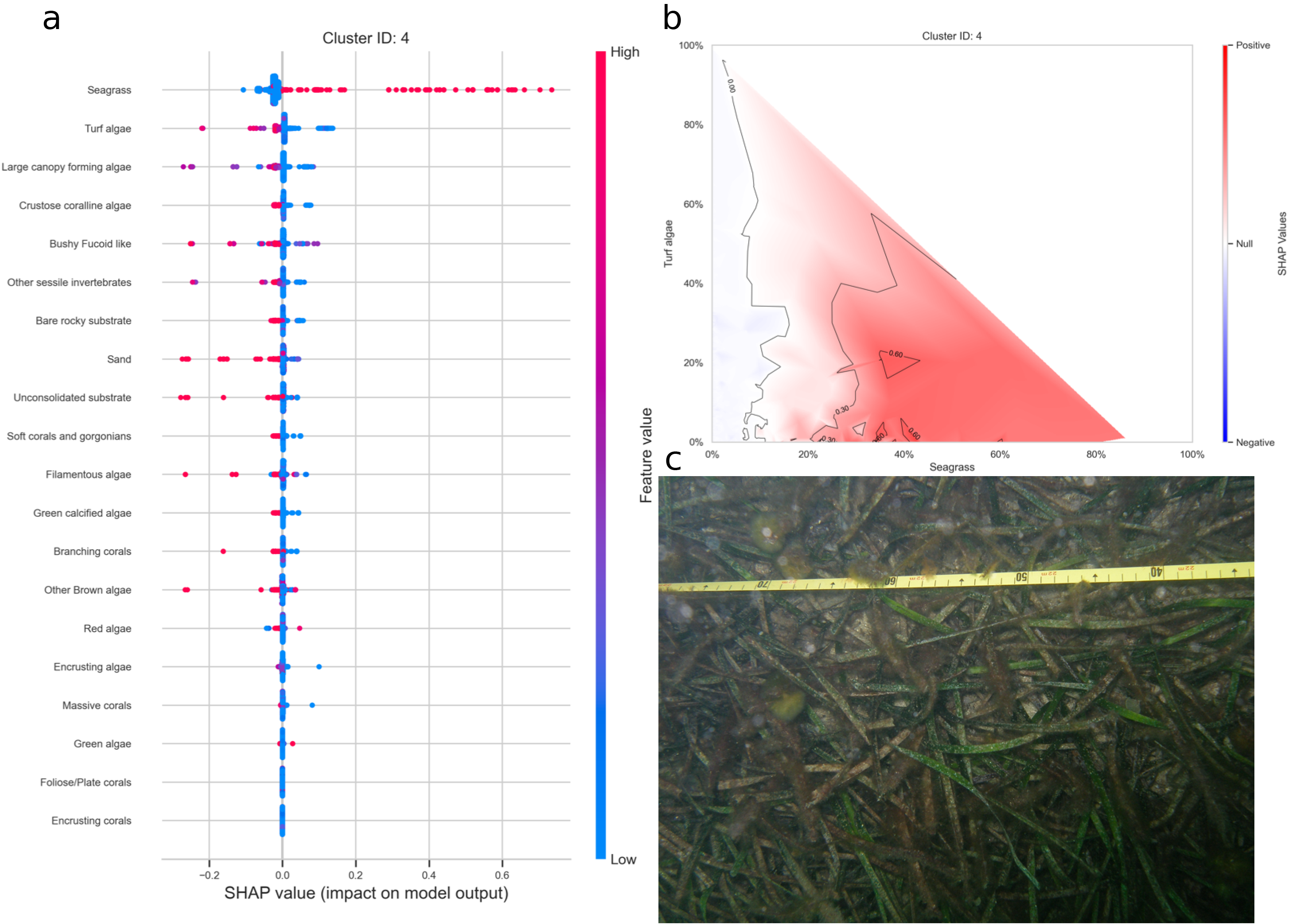

Figure 24: a. SHAP summary plot showing the impact of each habitat substrate on the classification. The position of each habitat group on the y-axis indicates its relative importance for the considered cluster. The position of each point along the x-axis indicates if the observation is associated with a lower or higher affinity with the cluster and the colour of the point indicates if the value of the cover of this habitat is rather high or low. b. Linear interpolation of the SHAP values for the two most influential variables for the cluster seagrass. c. Example of photoquadrat for one transect of the cluster seagrass categorised by HDBSCAN as exemplary.

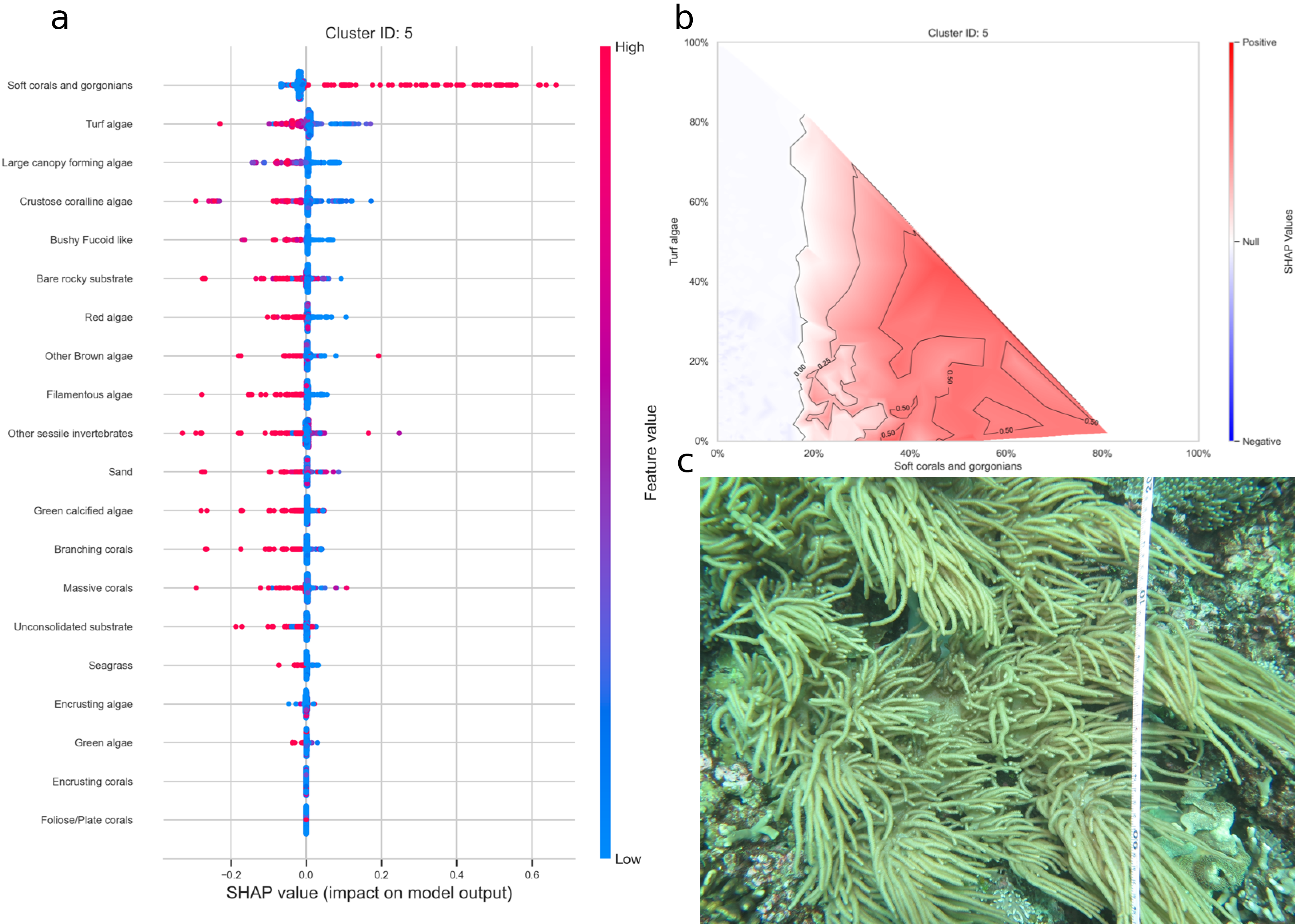

Figure 25: a. SHAP summary plot showing the impact of each habitat substrate on the classification. The position of each habitat group on the y-axis indicates its relative importance for the considered cluster. The position of each point along the x-axis indicates if the observation is associated with a lower or higher affinity with the cluster and the colour of the point indicates if the value of the cover of this habitat is rather high or low. b. Linear interpolation of the SHAP values for the two most influential variables for the cluster soft coral and gorgonians. c. Example of photoquadrat for one transect of the cluster soft coral and gorgonians categorised by HDBSCAN as exemplary.

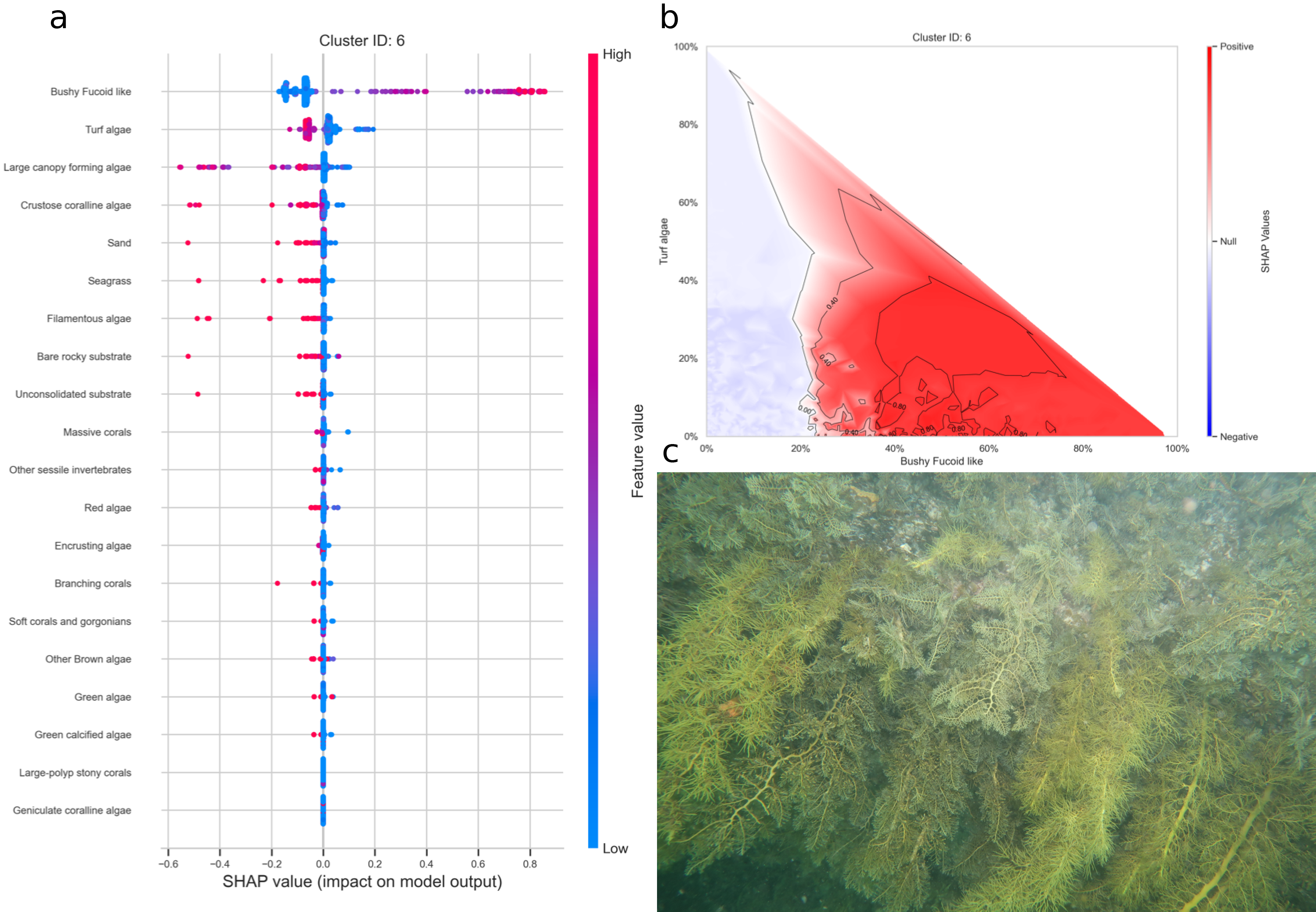

Figure 26: a. SHAP summary plot showing the impact of each habitat substrate on the classification. The position of each habitat group on the y-axis indicates its relative importance for the considered cluster. The position of each point along the x-axis indicates if the observation is associated with a lower or higher affinity with the cluster and the colour of the point indicates if the value of the cover of this habitat is rather high or low. b. Linear interpolation of the SHAP values for the two most influential variables for the cluster bushy fucoid-like. c. Example of photoquadrat for one transect of the cluster bushy fucoid-like categorised by HDBSCAN as exemplary.

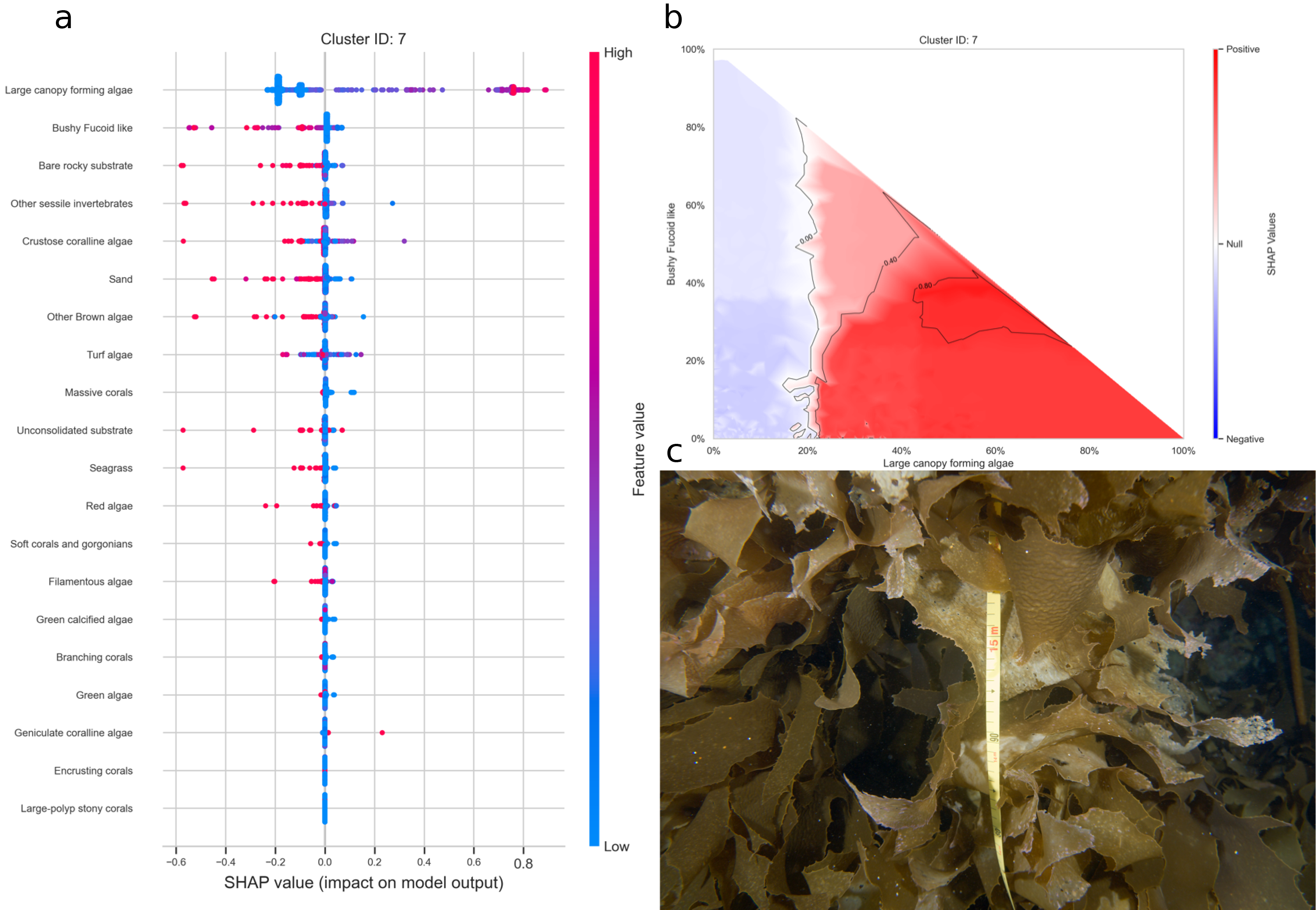

Figure 27: a. SHAP summary plot showing the impact of each habitat substrate on the classification. The position of each habitat group on the y-axis indicates its relative importance for the considered cluster. The position of each point along the x-axis indicates if the observation is associated with a lower or higher affinity with the cluster and the colour of the point indicates if the value of the cover of this habitat is rather high or low. b. Linear interpolation of the SHAP values for the two most influential variables for the cluster large canopy forming algae. c. Example of photoquadrat for one transect of the cluster large canopy forming algae categorised by HDBSCAN as exemplary.

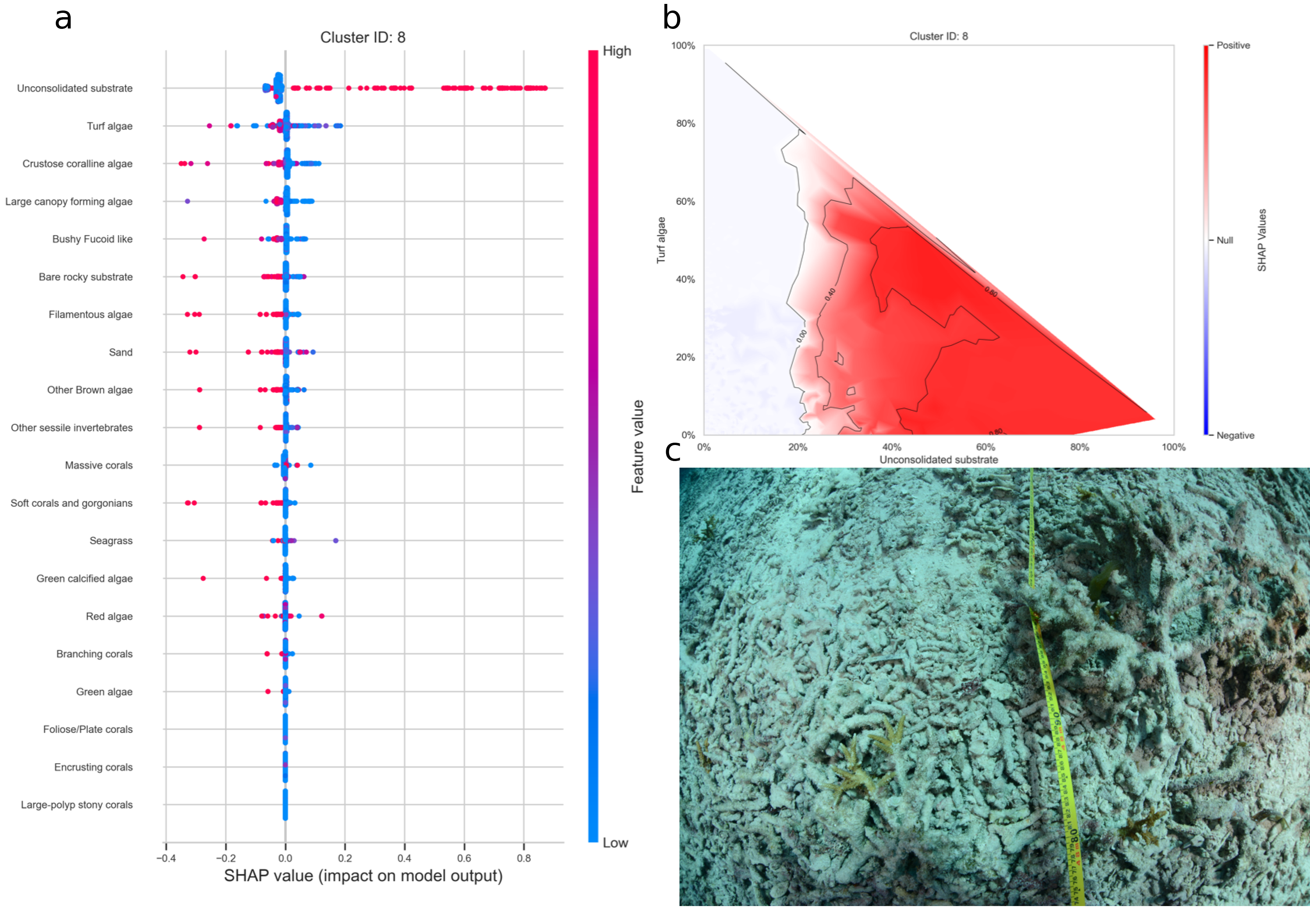

Figure 28: a. SHAP summary plot showing the impact of each habitat substrate on the classification. The position of each habitat group on the y-axis indicates its relative importance for the considered cluster. The position of each point along the x-axis indicates if the observation is associated with a lower or higher affinity with the cluster and the colour of the point indicates if the value of the cover of this habitat is rather high or low. b. Linear interpolation of the SHAP values for the two most influential variables for the cluster unconsolidated substrat. c. Example of photoquadrat for one transect of the cluster unconsolidated substrat categorised by HDBSCAN as exemplary.

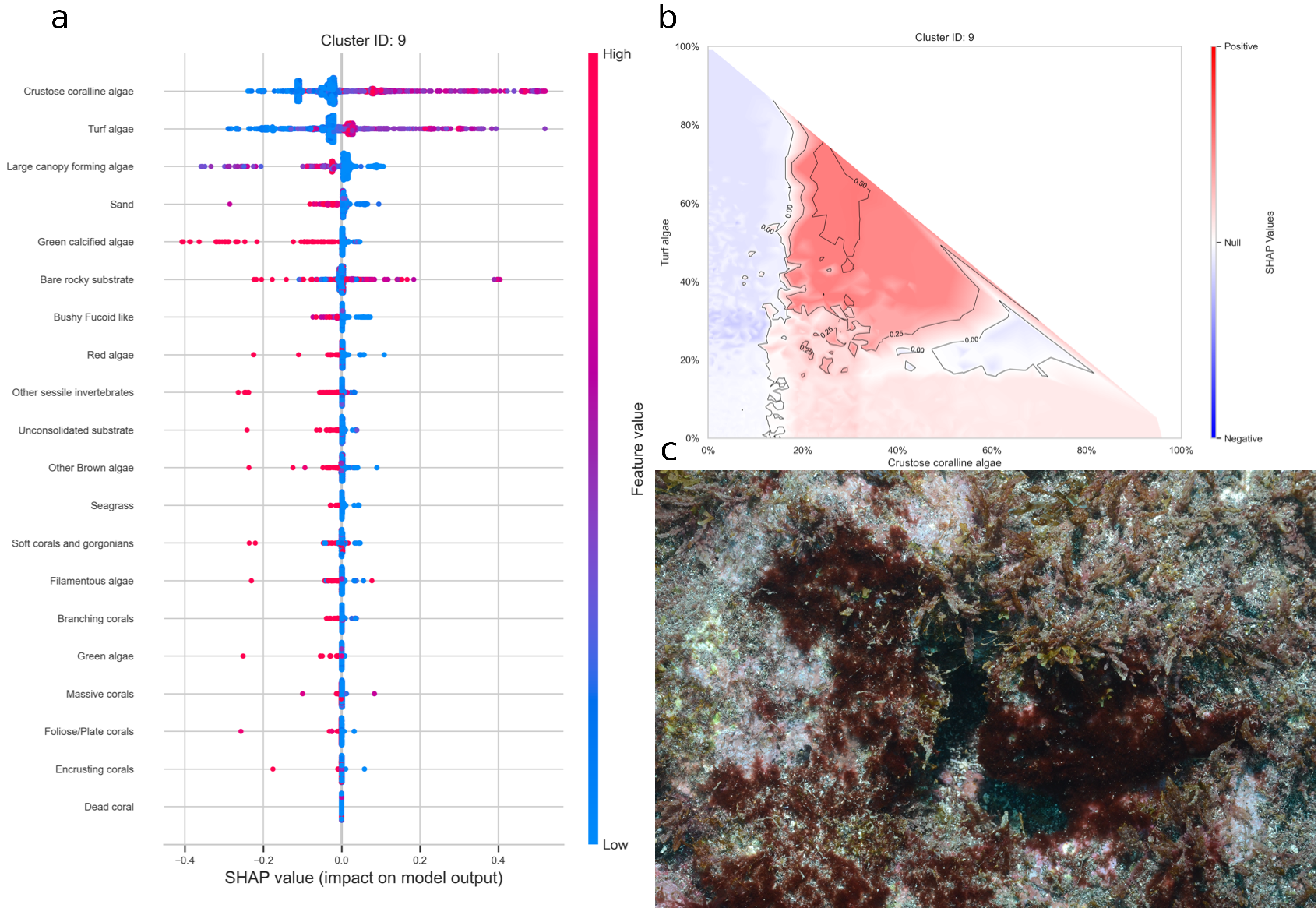

Figure 29: a. SHAP summary plot showing the impact of each habitat substrate on the classification. The position of each habitat group on the y-axis indicates its relative importance for the considered cluster. The position of each point along the x-axis indicates if the observation is associated with a lower or higher affinity with the cluster and the colour of the point indicates if the value of the cover of this habitat is rather high or low. b. Linear interpolation of the SHAP values for the two most influential variables for the cluster crustose coralline algae and turf. c. Example of photoquadrat for one transect of the cluster crustose coralline algae and turf categorised by HDBSCAN as exemplary.

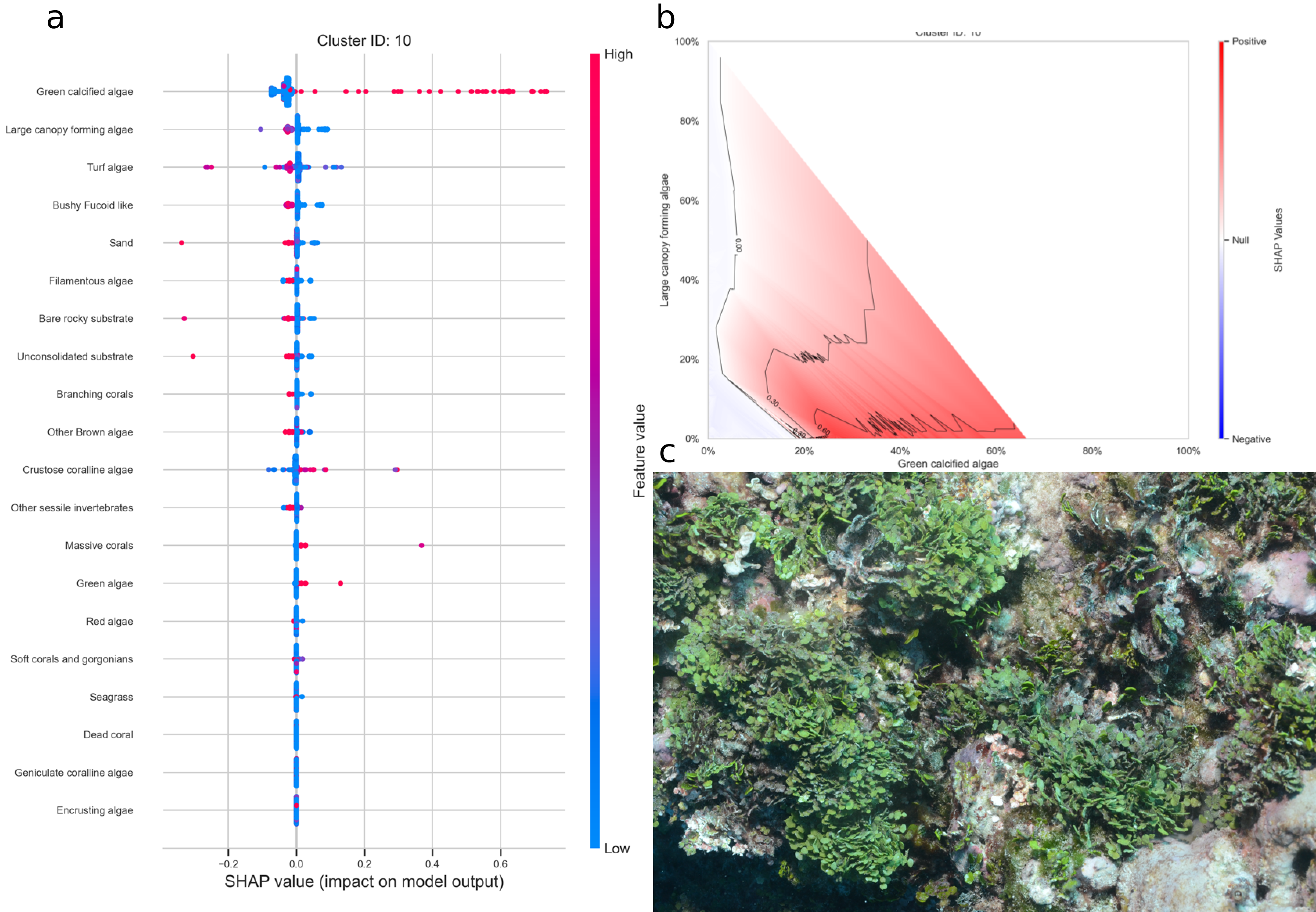

Figure 30: a. SHAP summary plot showing the impact of each habitat substrate on the classification. The position of each habitat group on the y-axis indicates its relative importance for the considered cluster. The position of each point along the x-axis indicates if the observation is associated with a lower or higher affinity with the cluster and the colour of the point indicates if the value of the cover of this habitat is rather high or low. b. Linear interpolation of the SHAP values for the two most influential variables for the cluster green calcified algae. c. Example of photoquadrat for one transect of the cluster green calcified algae categorised by HDBSCAN as exemplary.

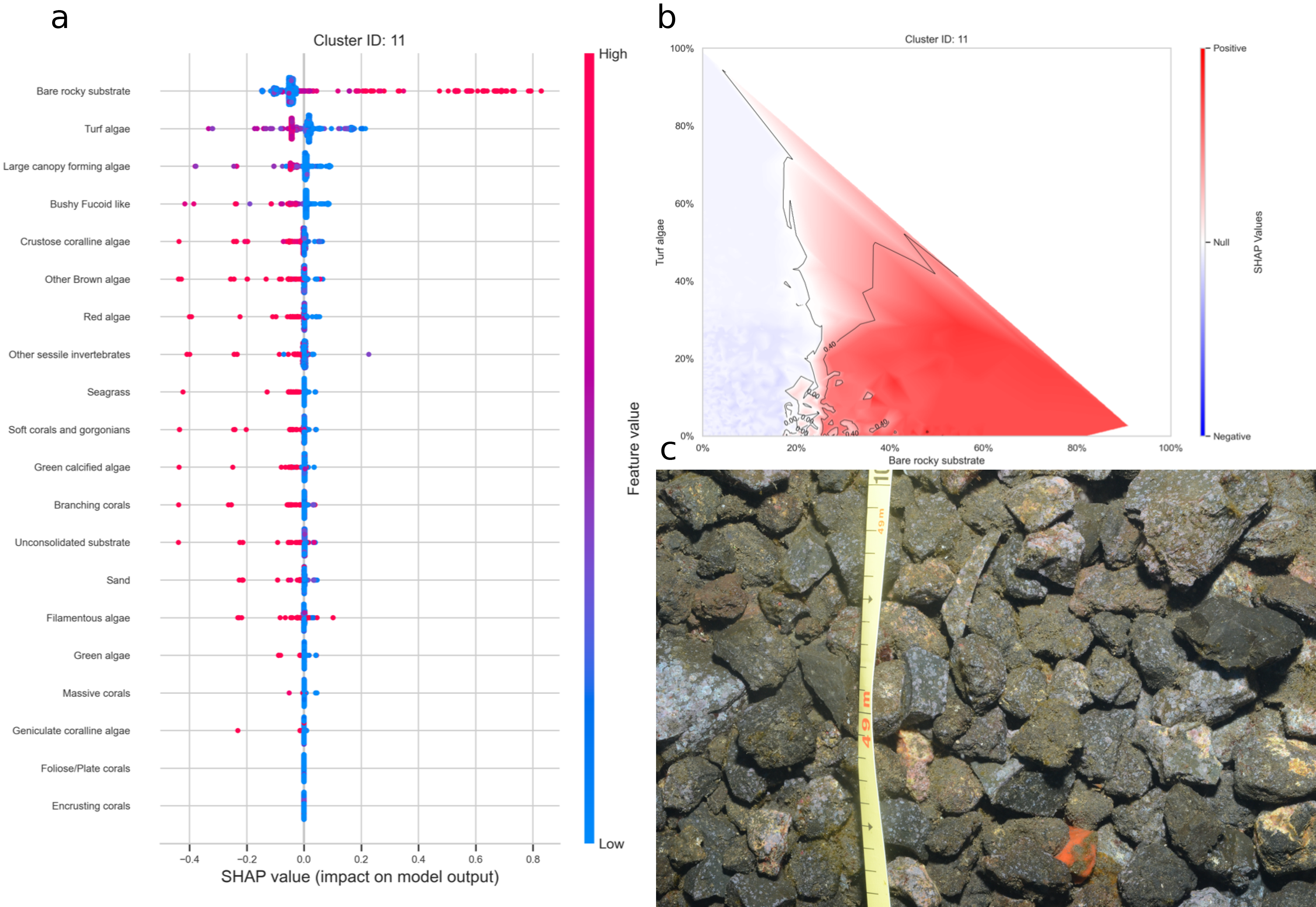

Figure 31: a. SHAP summary plot showing the impact of each habitat substrate on the classification. The position of each habitat group on the y-axis indicates its relative importance for the considered cluster. The position of each point along the x-axis indicates if the observation is associated with a lower or higher affinity with the cluster and the colour of the point indicates if the value of the cover of this habitat is rather high or low. b. Linear interpolation of the SHAP values for the two most influential variables for the cluster bare substrate. c. Example of photoquadrat for one transect of the cluster bare substrate categorised by HDBSCAN as exemplary.

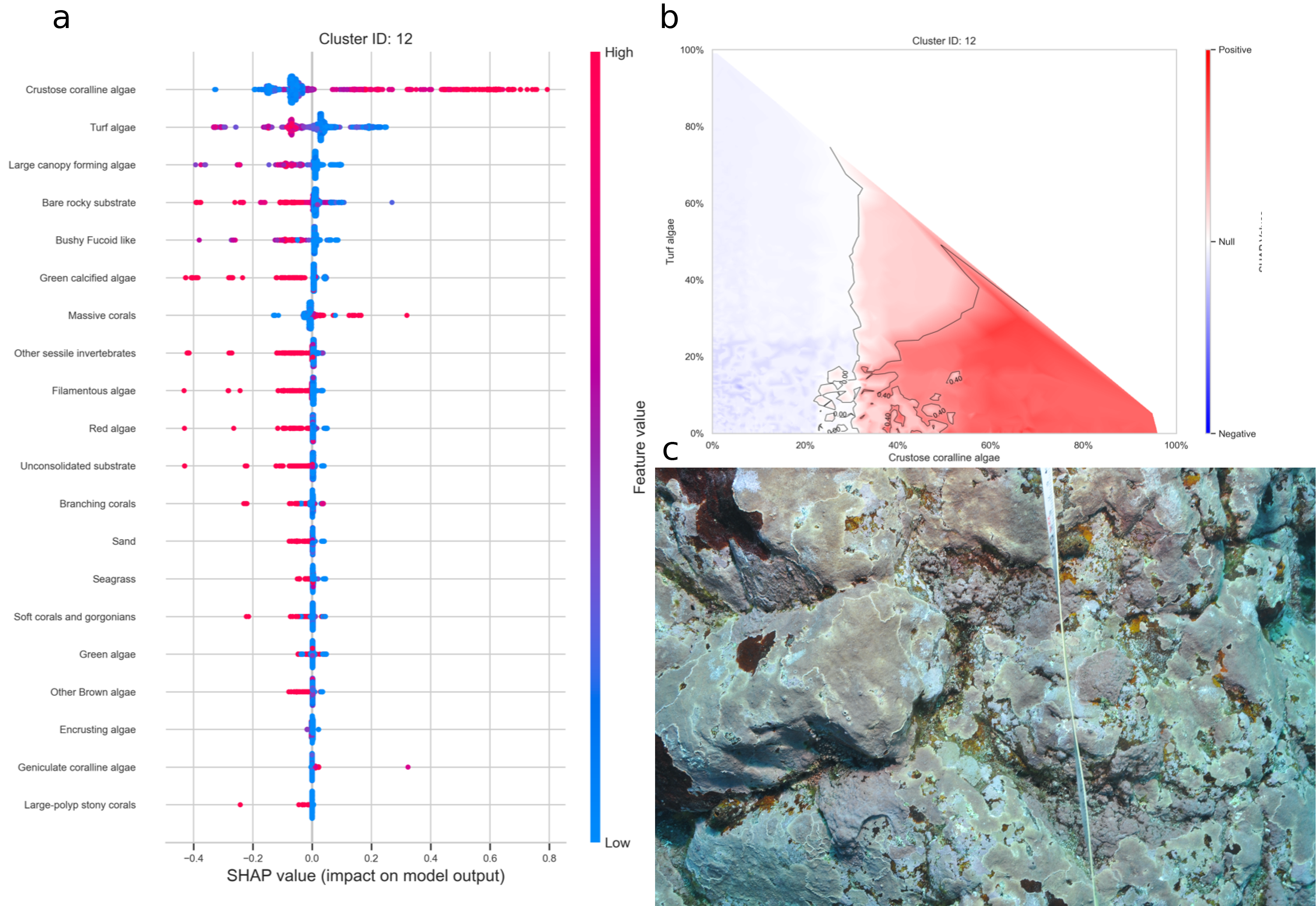

Figure 32: a. SHAP summary plot showing the impact of each habitat substrate on the classification. The position of each habitat group on the y-axis indicates its relative importance for the considered cluster. The position of each point along the x-axis indicates if the observation is associated with a lower or higher affinity with the cluster and the colour of the point indicates if the value of the cover of this habitat is rather high or low. b. Linear interpolation of the SHAP values for the two most influential variables for the cluster crustose coralline algae. c. Example of photoquadrat for one transect of the cluster crustose coralline algae categorised by HDBSCAN as exemplary.

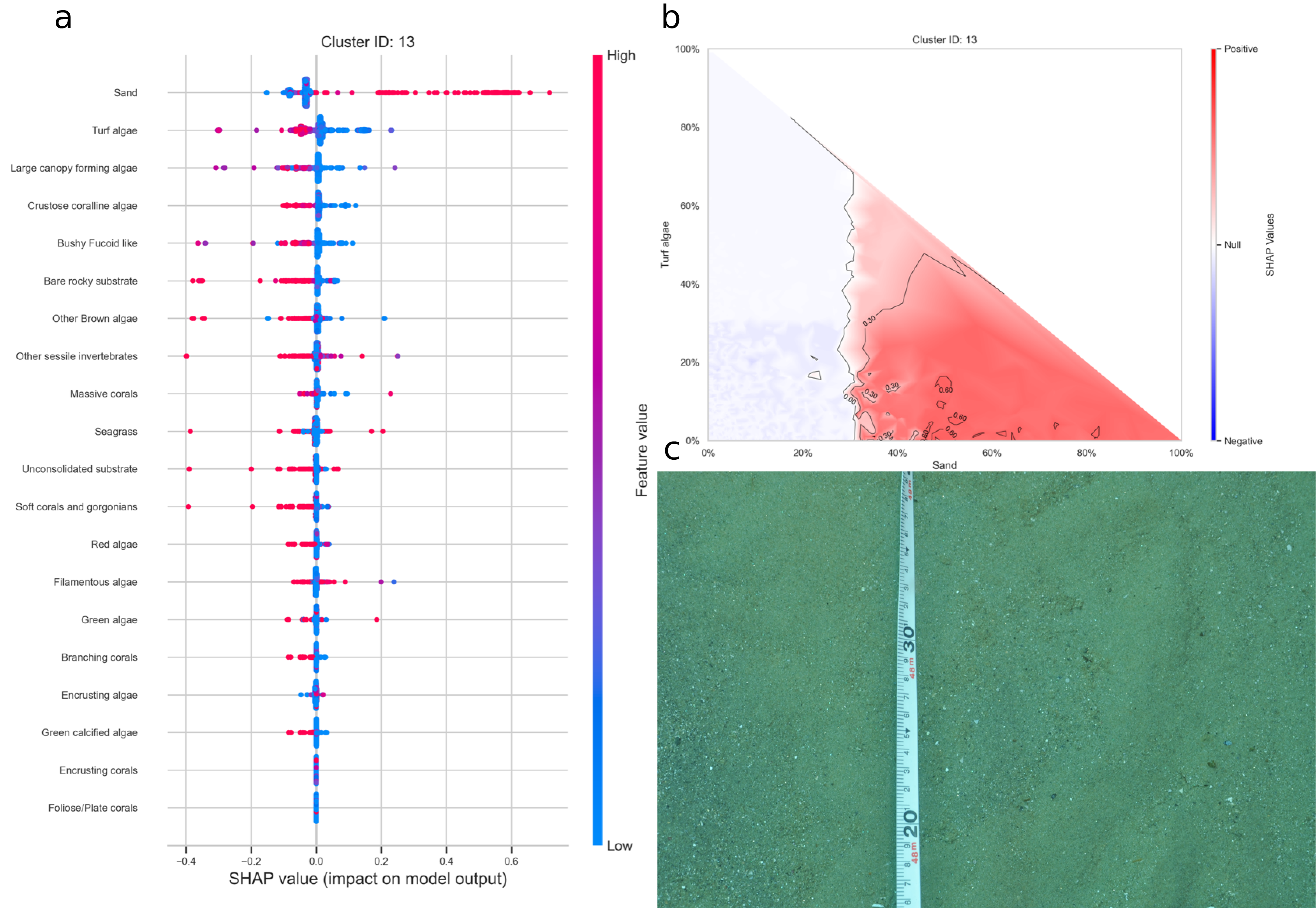

Figure 33: a. SHAP summary plot showing the impact of each habitat substrate on the classification. The position of each habitat group on the y-axis indicates its relative importance for the considered cluster. The position of each point along the x-axis indicates if the observation is associated with a lower or higher affinity with the cluster and the colour of the point indicates if the value of the cover of this habitat is rather high or low. b. Linear interpolation of the SHAP values for the two most influential variables for the cluster sand. c. Example of photoquadrat for one transect of the cluster sand categorised by HDBSCAN as exemplary.

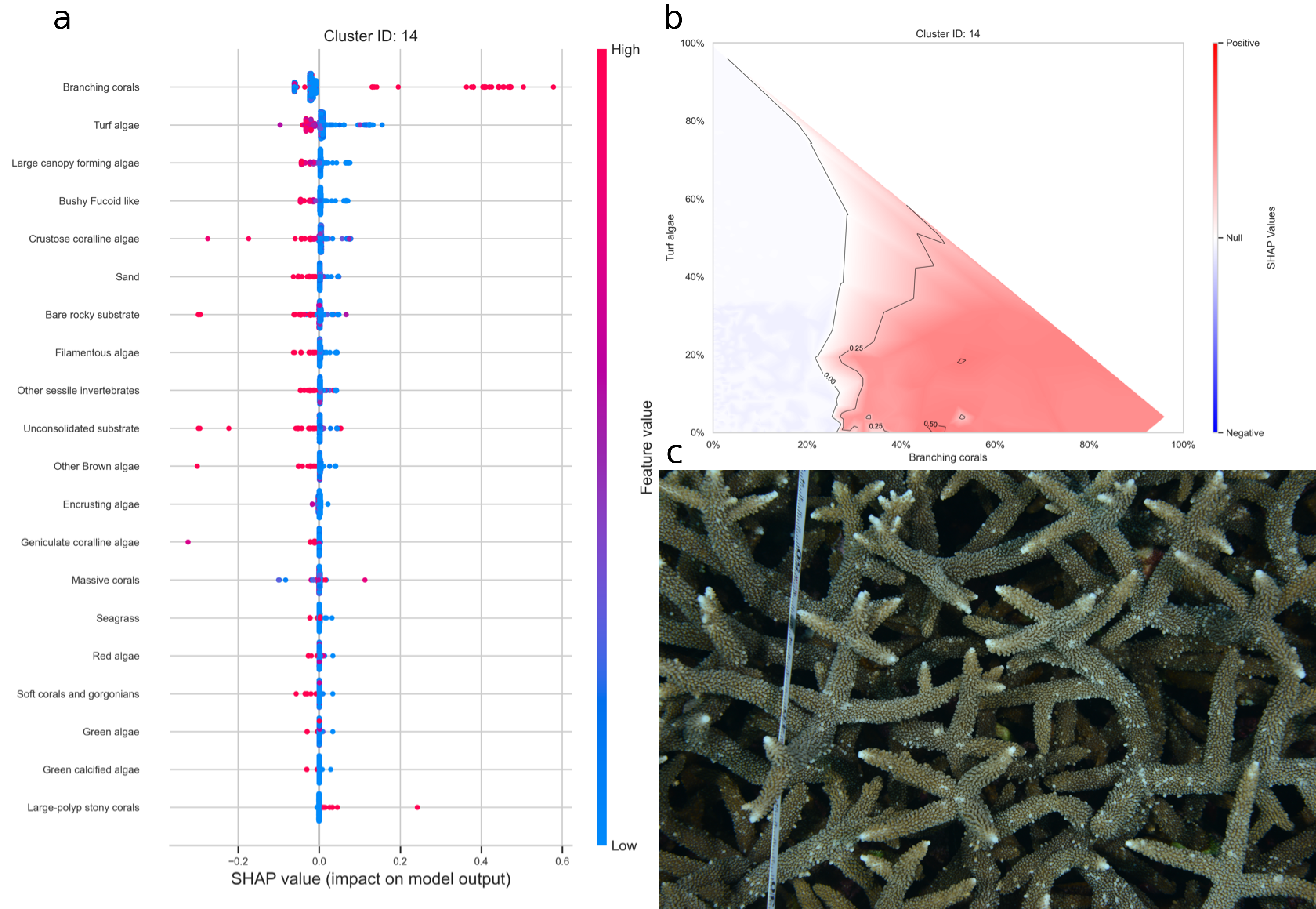

Figure 34: a. SHAP summary plot showing the impact of each habitat substrate on the classification. The position of each habitat group on the y-axis indicates its relative importance for the considered cluster. The position of each point along the x-axis indicates if the observation is associated with a lower or higher affinity with the cluster and the colour of the point indicates if the value of the cover of this habitat is rather high or low. b. Linear interpolation of the SHAP values for the two most influential variables for the cluster branching coral. c. Example of photoquadrat for one transect of the cluster branching coral categorised by HDBSCAN as exemplary.

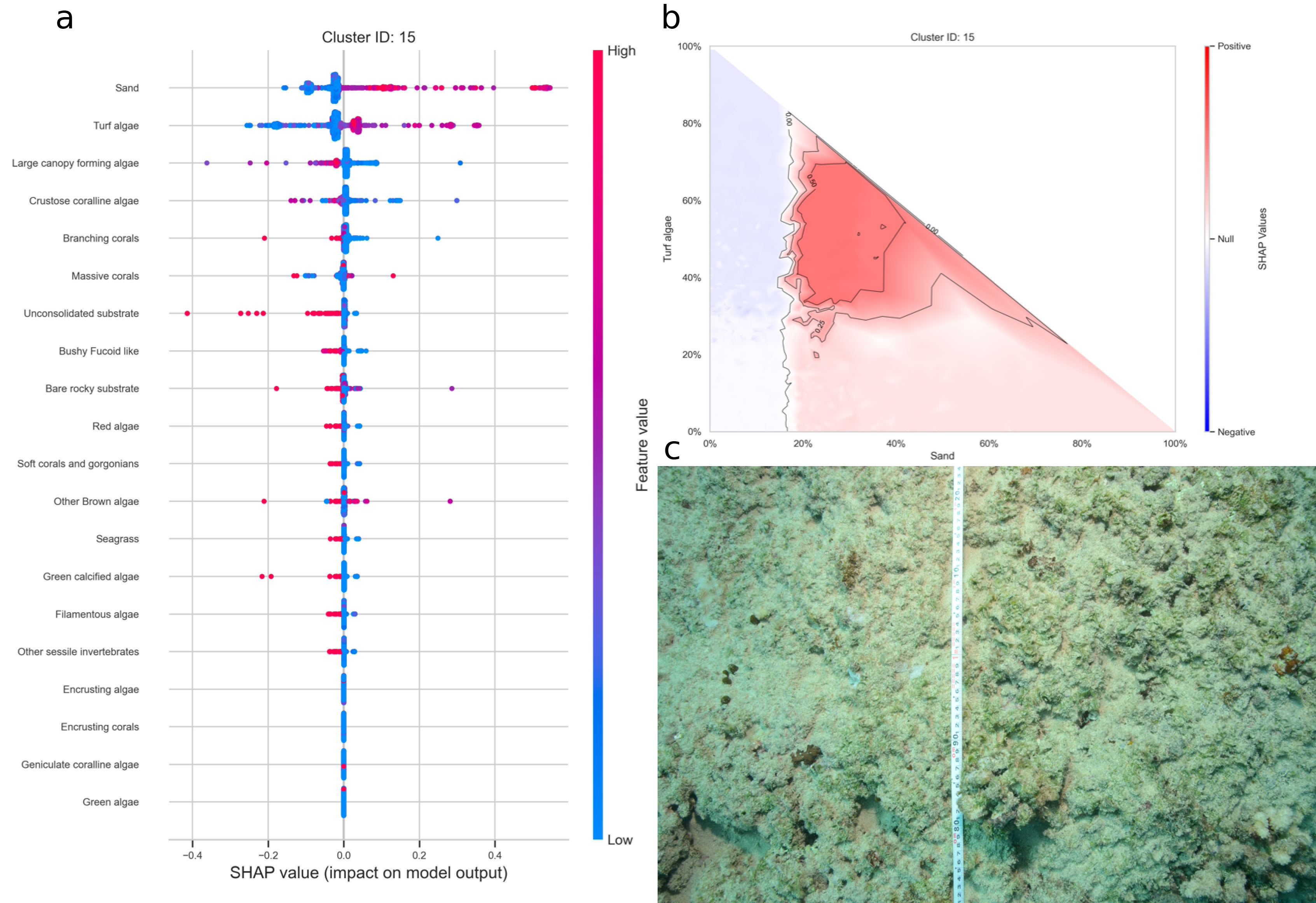

Figure 35: a. SHAP summary plot showing the impact of each habitat substrate on the classification. The position of each habitat group on the y-axis indicates its relative importance for the considered cluster. The position of each point along the x-axis indicates if the observation is associated with a lower or higher affinity with the cluster and the colour of the point indicates if the value of the cover of this habitat is rather high or low. b. Linear interpolation of the SHAP values for the two most influential variables for the cluster sand and turf algae. c. Example of photoquadrat for one transect of the cluster sand and turf algae categorised by HDBSCAN as exemplary.

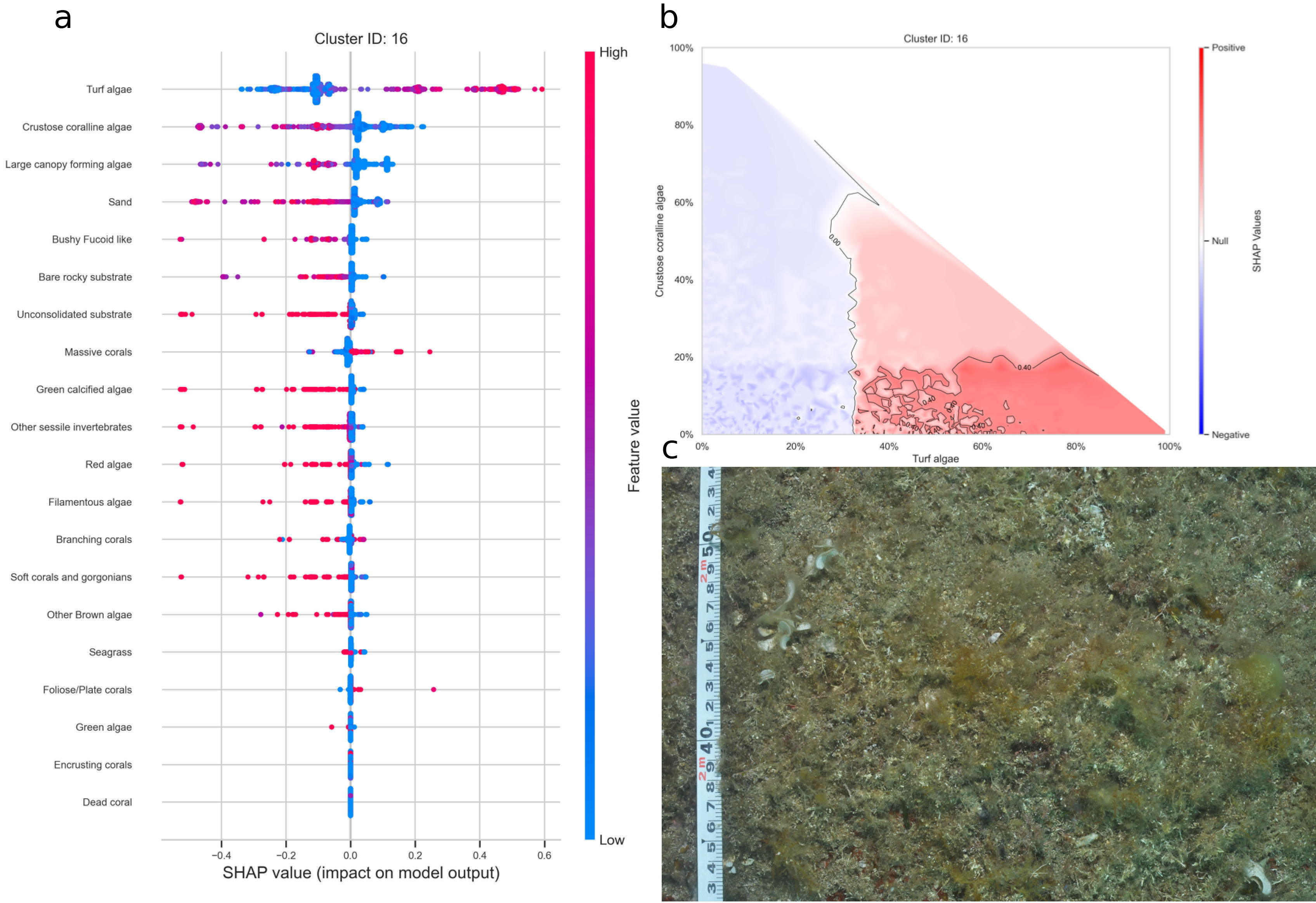

Figure 36: a. SHAP summary plot showing the impact of each habitat substrate on the classification. The position of each habitat group on the y-axis indicates its relative importance for the considered cluster. The position of each point along the x-axis indicates if the observation is associated with a lower or higher affinity with the cluster and the colour of the point indicates if the value of the cover of this habitat is rather high or low. b. Linear interpolation of the SHAP values for the two most influential variables for the cluster turf algae. c. Example of photoquadrat for one transect of the cluster turf algae categorised by HDBSCAN as exemplary.

### Appendix D

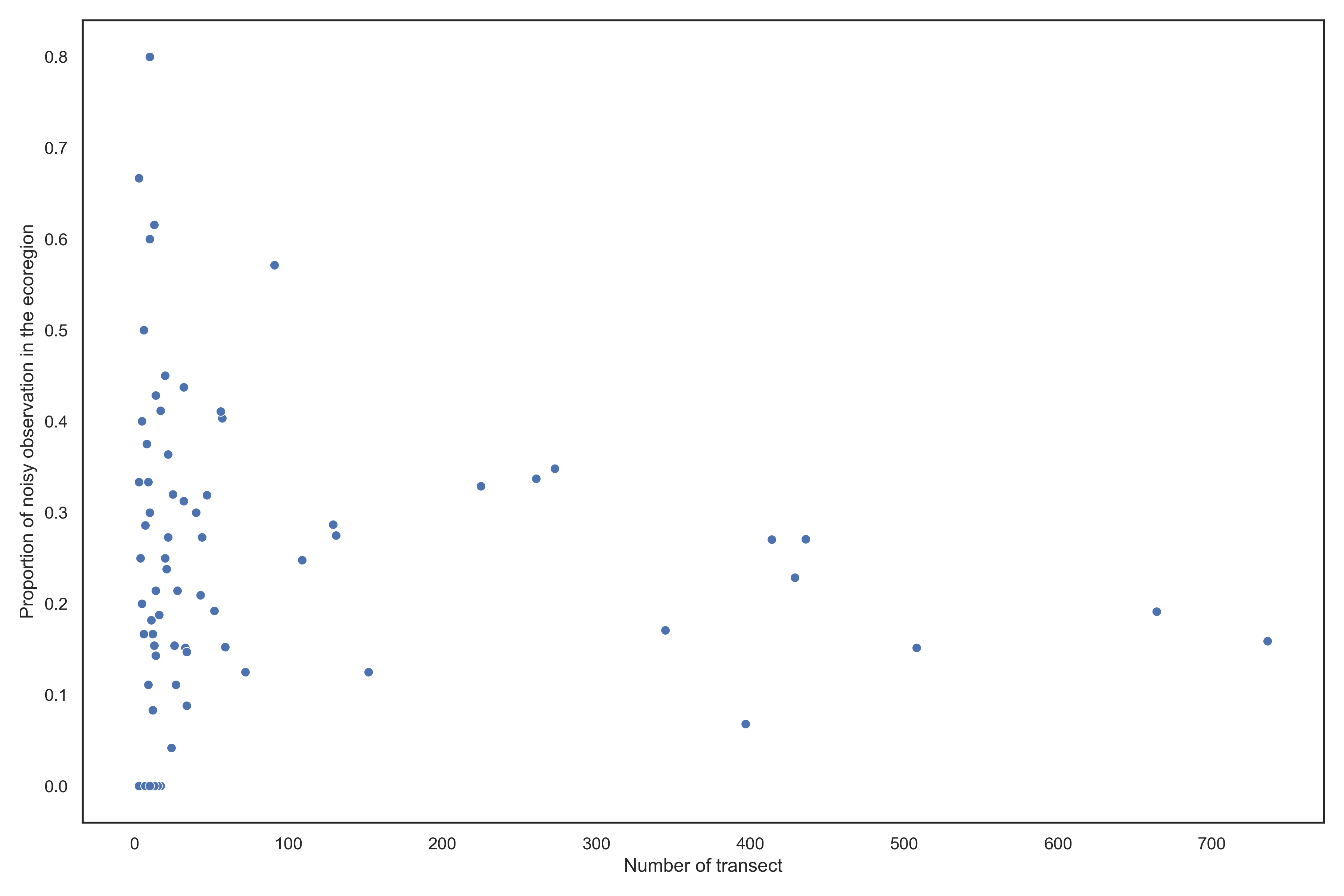

Figure 37: Plot of the proportion of transect classified as noisy by the UMAP-HDBSCAN pipeline as a function of the log10(number of transects) sampled within an ecoregion

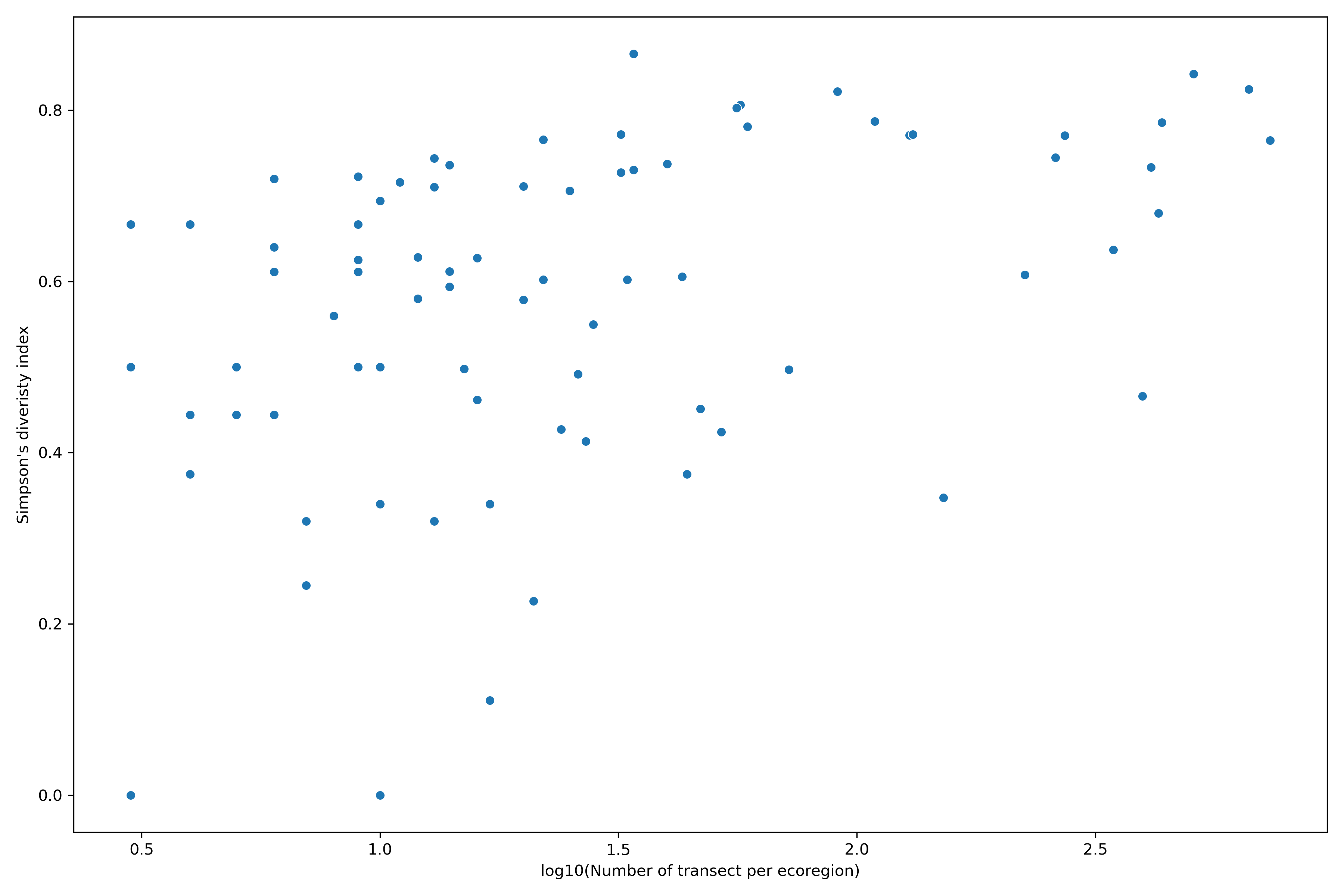

Figure 38: Plot of the Gini-Simpson diversity index as a function of the log10(number of transects) sampled within an ecoregion

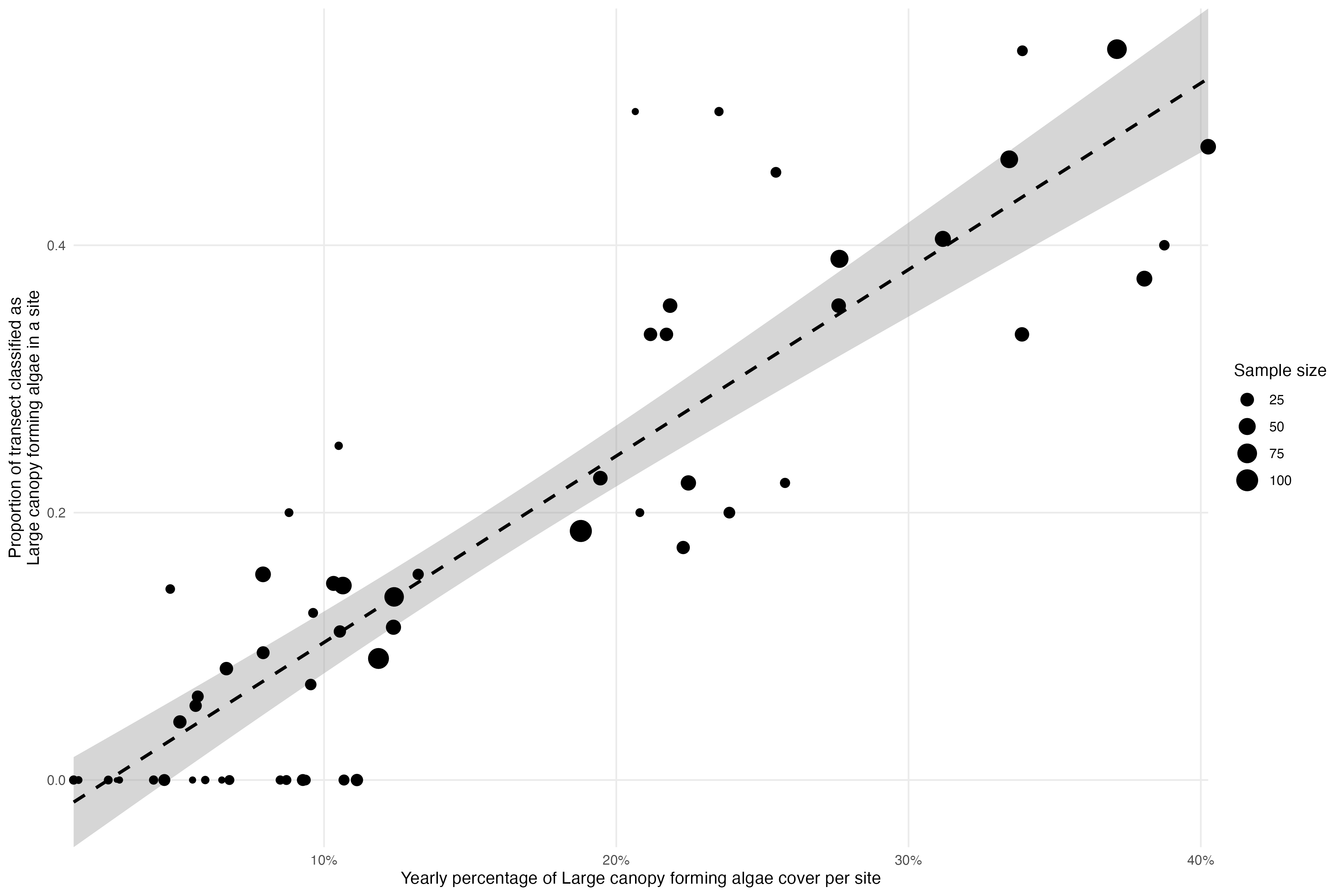

Figure 39: Proportion of transect classified as Canopy forming algae each year as a function of yearly average cover of Canopy forming algae. The size of the points depends on the number of transects sampled that year for the considered site. The dashed line represent a linear regression model fitted to the data with its 95% confidence interval.

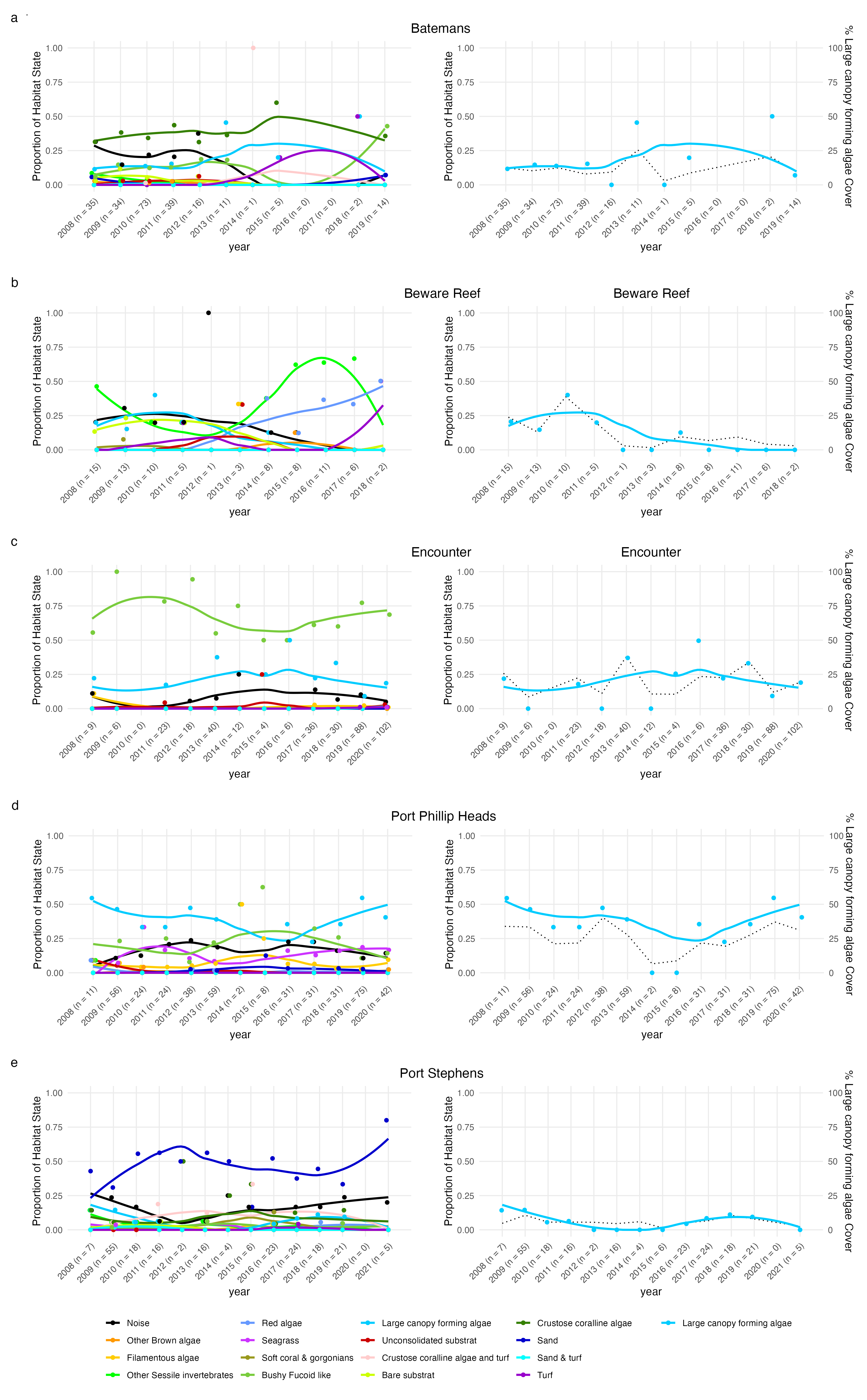

Figure 40: Same plot as Fig. 5 in the main text with all habitat states displayed. See Fig. 5 in main text for more details.
